# A causal model of human growth and its estimation using temporally-sparse data

**DOI:** 10.1101/2022.10.10.511559

**Authors:** John A. Bunce, Catalina I. Fernández, Caissa Revilla-Minaya

## Abstract

Existing models of human growth provide little insight into underlying mechanisms responsible for inter-individual and inter-population variation in children’s growth trajectories. Building on general theories linking growth to metabolic rates, we develop a causal parametric model of height and weight growth incorporating a representation of human body allometry and a process-partitioned representation of ontogeny. This model permits separation of metabolic causes of growth variation, potentially influenced by nutrition and disease, from allometric factors, potentially under stronger genetic control. We estimate model parameters using a Bayesian multilevel statistical design applied to temporally-dense height and weight measurements of U.S. children, and temporally-sparse measurements of Indigenous Amazonian children. This facilitates a comparison of the contributions of metabolism and allometry to observed cross-cultural variation in the growth trajectories of the two populations, and permits simulation of the effects of healthcare interventions on growth. This theoretical model provides a new framework for exploring the causes of growth variation in our species, while potentially guiding the development of appropriate, and desired, healthcare interventions in societies confronting growth-related health challenges, such as malnutrition and stunting.

## Introduction

Between birth and adulthood, humans generally increase in height and weight following trajectories unique to our species, and notably different from those of other mammals (von Bertalanffy 1957; Bogin 1999; Hamada and Udono 2002). However, like many widely-distributed animals, the typical adult body form, and thus the typical shape of the growth trajectory, differs between populations in different parts of the world (Eveleth and Tanner 1990). Some aspects of body form are heavily influenced by genes (Lello et al. 2018; Zoccolillo et al. 2020). Thus, both drift and natural selection may contribute to this intra-species variation through their action on genes at the population level over generations in different terrestrial ecosystems (e.g., Allen’s and Bergmann’s Rules: Katzmarzyk and Leonard (1998)). Other important contributions to variation in human body form come from environmental factors, such as food availability, pathogen exposure, and chronic stress (Stewart et al. 2013; Li et al. 2020; Tanner et al. 1982; Bogin 2022), that directly, and differentially, affect the growth of individuals in different societies around the world. Distinguishing among these contributions to population-level variation in growth is a major challenge (Hruschka 2021), both for our understanding of human ontogeny, and for the provision of appropriate healthcare support to individuals whose particular patterns of growth result from exposure to health challenges that can be alleviated (e.g., stunting and wasting as indicators of malnutrition: World Health Organization and United Nations Children’s Fund (2009); de Onis and Branca (2016)). Here we present a new approach to exploring these different causes of variation by deriving a model of human growth from first principles of metabolism and allometry.

### Growth models

Over the last century, auxology, the study of human growth, has produced a rich set of mathematical models to describe the complex growth trajectories of children. Growth models are often classified as “parametric” or “non-parametric”. Parametric models define a specific range of possible forms for the growth trajectory, implemented as a mathematical relationship between a (relatively small) number of parameters, the values of which (and the shape of the resulting trajectory) may vary from person to person. Ideally, the mathematical relationship is derived from some causal theory of growth, and the parameter values thus have a mechanistic interpretation. However, for most popular parametric models (e.g., Jolicoeur et al. (1992)), generally neither is true (Hauspie and Molinari 2004; Gasser et al. 2004). Non-parametric models, e.g., splines, are explicitly descriptive, and can be made to fit growth data arbitrarily well. However, their (generally numerous) parameters are, by design, biologically uninterpretable, offering no causal insight. A third class, called “shape invariant” models, employ either a parametric (e.g., QEPS: Nierop et al. (2016)) or non-parametric (e.g., SITAR: Cole et al. (2010)) function to represent the population mean trajectory, and then use a smaller set of individual-level parameters to account for each individual’s deviation from this mean trajectory (Gasser et al. 2004). All of these models are valuable tools for representing and comparing individual growth trajectories given the temporal resolution of most available data (where higher resolution data motivates structurally different models: Lampl (2012); Suki and Frey (2017)). Importantly, however, none of these models is derived from a causal theory of human growth, i.e., a theory that identifies biological quantities measurable independently of growth, and proposes a relationship between them that causes an observable growth trajectory. Existing models therefore provide little insight into the mechanisms through which individual characteristics (e.g., genes) and experiences (e.g., diet, illness) interact to produce an observed pattern of growth.

Though rarely applied specifically to humans, general causal models of organismal growth have a long history in biology. Pütter (1920) proposed a theory, further developed and popularized by von Bertalanffy (1938, 1957), such that organisms grow when the energy released by breaking down molecules acquired from the environment (via catabolism) exceeds the energy required for homeostasis (e.g., cellular maintenance). The excess energy can be used to synthesize new structural molecules (via anabolism) using both the products of catabolism and other substrates acquired from the environment. Because all living cells perform catabolic reactions, an organism’s total mean catabolic rate is a function of its mass. However, in an environment with abundant energy-containing substrates, the production of the excess energy required for anabolic structural growth is a function of the surface area of the body’s interface with the environment through which structural and energy-containing molecules are absorbed. In recent years, this basic principle has been incorporated into more general theories, detailing how environmental substrates of a given energy density, acquired by cells and transformed into reserves, are then allocated to maintenance, growth, and reproduction (Kooijman 2001, 2010), how anabolic and catabolic processes (and therefore growth) depend on temperature (Gillooly et al. 2001), and how anabolism (and therefore growth) is limited by the structure of the supply network (e.g., capillaries) transporting environmental substrates to internal cells (West et al. 1997, 2001). We develop a causal theory of growth tailored to humans by combining the basal model of Pütter (1920) and von Bertalanffy (1938) with a representation of human body allometry, and a process-partitioned representation of human ontogeny. This results in a new parametric model of the metabolic processes and allometric relationships that underlie both the unique species-typical pattern, and the individual variations, of human growth.

### Model estimation

We apply our new model to height and weight measurements from two populations that, on average, exhibit different patterns of growth: affluent U.S. children born in the late 1920s and Indigenous Matsigenka children from Amazonian Peru measured in 2017-2019. Importantly, these two datasets also differ in temporal resolution: each U.S. child was measured (at least) annually from birth to age 18 (Tuddenham and Snyder 1954), while most Matsigenka were measured (by authors C.R.M. and J.A.B.) only once or twice between the ages of one and 24. We believe the temporal sparsity of the Matsigenka dataset is typical of data feasibly collected in many rural, isolated, or marginalized societies around the world. Attention to such societies is essential, both to understand the range of growth variation in our species, and, even more importantly, to provide desired and appropriate support to address growth-affecting health challenges faced by many of these populations.

To accommodate the differing temporal resolutions of the two datasets, we design a Bayesian multilevel (mixed-effects) estimation strategy (McElreath 2020; Vincenzi et al. 2020; Johnson 2015). Given our theoretical model, this allows us to quantify differences between U.S. and Matsigenka children in metabolic rates, plausibly influenced by population-level differences in diet and/or disease exposure, as well as differences in allometric relationships, potentially (though not necessarily: Tanner et al. (1982)) under stronger genetic influence. Additionally, by modifying a fit model’s metabolic parameters, we can simulate population-specific effects on growth of a healthcare intervention affecting metabolic rates, such as dietary supplementation and/or disease reduction. An important goal of this approach is a tool to suggest when, and what form of, healthcare interventions would increase child well-being in a particular population, using data feasibly collected across a broad range of human societies.

## Methods

### Derivation of the theoretical growth model

In the Pütter (1920) and von Bertalanffy (1938) theory of growth, mass increases when the rate of anabolism (using energy to construct molecules) is greater than the rate of catabolism (breaking down molecules to release energy). All cells perform catabolic reactions. Anabolism is contingent on the acquisition of molecules (containing both energy and building materials) from the environment. Only parts of the organism in contact with the environment can acquire such molecules. Therefore, anabolism is a function of an organism’s absorbing surface area (under the assumption of abundant food: Kooijman (2010)), while catabolism is a function of an organism’s mass:

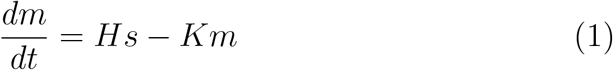

where *m* is mass, *s* is surface area, *H*≥0 is synthesized mass (i.e., substrate molecules chemically bonded via anabolism) per unit of an organism’s absorbing surface, and *K*≥0 is destructed mass (i.e., molecules whose chemical bonds are broken via catabolism) per unit of an organism’s mass.

We are interested in the rate of longitudinal (i.e., height) growth per unit time. The shape of the body is stylized as a cylinder with height *h*, radius *r*, and substrate-absorbing (i.e., intestinal) surface area 2*πrh* (Appendix A, with modification in Appendix D.1.2). Equation 1 can be used to approximate the rate of growth in mass (= volume *·* density):

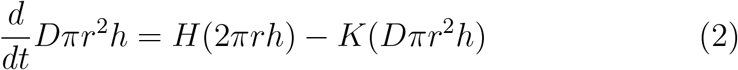

where *D* is density. Solving Equation 2 for the longitudinal growth rate 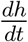 yields:

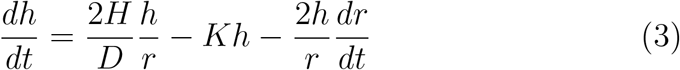

We assume a power law relationship between *r* and *h* such that *r* = *h*^*q*^, where 0 *< q <* 1 represents the allometric relationship between radius and height as the body grows in both dimensions (Appendix A). Substituting for *r* in equation 3 yields:

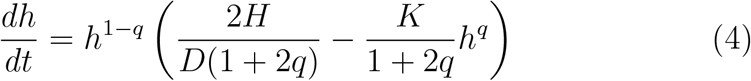

Under the same assumptions (Appendix B), re-writing *s* in equation 1 in terms of *m* yields the rate of growth in mass:

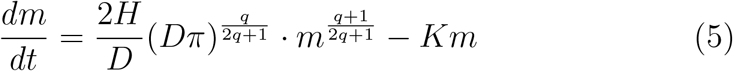

Assuming *D* = 1 g/cm^3^, solving differential equations 4 and 5 for *h*(*t*) and *m*(*t*) yields:

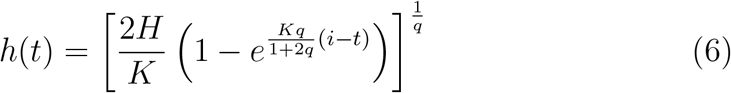

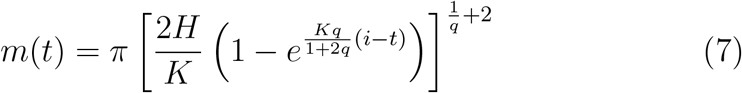

for all *t* ≥ *i*, where *t* is total age in years since conception, and *i* ≥ 0 is the age at which growth is initiated, such that, for growth in early life, *i* = 0 at conception. Detailed derivations and solutions, as well as justifications for assumptions, are provided in Appendices A and B.

It is well-established that different tissues in the human body, e.g., bones of the leg versus torso, grow with different acceleration profiles, attaining high growth velocities at different ages during ontogeny (Eveleth and Tanner 1990, pg 35, 82). There is also evidence that groups of tissues may undergo coordinated cycles of faster and slower growth on time scales of several years (Butler et al. 1990), several months (Togo and Togo 1982), and several days (Lampl et al. 1992; Lampl and Johnson 1993), likely due to hormonal regimes whose effects and interactions are still poorly understood (Appendix C). The additive effect of these partially-overlapping growth processes contributes to the complexity of the human growth trajectory, such that greater complexity (e.g., frequency of changes in the sign of acceleration) becomes more apparent the higher the temporal resolution of measurements.

In principle, equations 6 and 7 are general enough to apply at any time scale to any growing tissue linked via blood vessels to the intestinal surface. Total height (or weight) can thus be modeled as the sum of multiple equation 6s (or 7s), where each component equation represents one cycle of the growth of a particular tissue (or group of tissues). The number of components to add together within such a composite model is as much an empirical as a theoretical question, and depends on the temporal resolution of the data to which the model will be fit, as well as the expected measurement error. Here we construct a composite model with five additive components (justified in Appendix C). Each component function may have different values for the parameters *H, K, q*, and *i*, represented by subscripts 1 to 5. This yields the following 20-parameter composite growth models for height (equation 8) and weight (equation 9) at total age *t*:

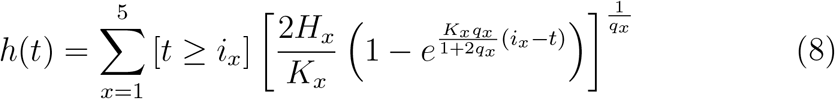

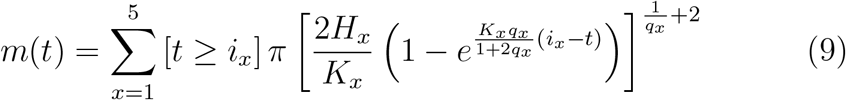

such that *i*_1_ = 0 represents conception, and [*t* ≥ *i*_*x*_] is Iverson notation (Knuth 1992), taking the value of one if the condition within brackets evaluates to True, and is zero otherwise.

Figure 1 presents a graphical illustration of the composite growth model for height and weight. For convenience, the five component processes are labeled according to the ontogenetic phase in which each has the largest effect on overall growth. The relative positions of the five component functions with respect to age are estimated empirically (with guidance from priors) through fitting the model, and are not fixed *a priori*. Thus, use of this growth model does not require us to define the five growth processes in terms of age. For illustration, the height and weight plots in Figure 1 are generated with different model fitting parameters to better match each type of growth data (described in Appendices D.3 and F.1, respectively). As explained in Appendix D.1.1, the model is fit to skeletal cell weight rather than total body weight.

**Figure 1.**
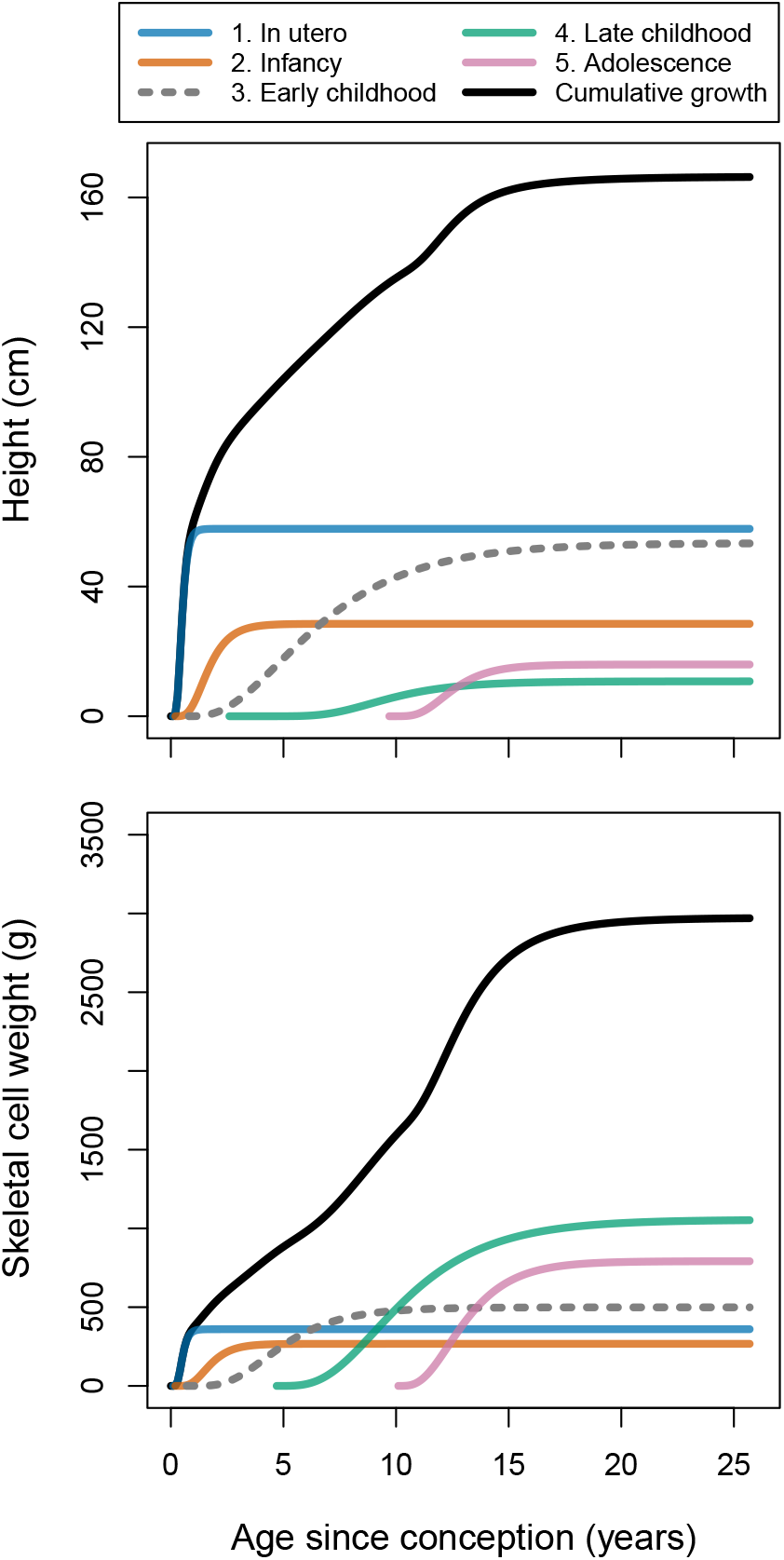
Illustration of the composite growth model comprising the sum of five component growth processes, fit to height and skeletal cell weight (Appendix D.1.1) of U.S. girls. Height (upper) and weight (lower) plots are generated from two variants of the model, explained in Appendices D.3 and F.1, respectively.

### Empirical analysis

We fit the composite growth model (equations 8 and 9) to two datasets: 1) temporally-dense height and weight measures from 70 girls and 66 boys born in 1928 and 1929 to parents of relatively high socioeconomic status in Berkeley, California, U.S.A. (Tuddenham and Snyder 1954); and 2) temporally-sparse measures from 196 girls and 179 boys collected by C.R.M. and J.A.B. between 2017 and 2019 in four Matsigenka Native Communities inside Manu National Park in the lowland Amazonian region of Peru. Matsigenka in these remote communities are horticulturalists, gatherers, fishers, and hunters who produce most of their own food and have relatively little (though increasing) dependence on broader Peruvian society (Bunce and McElreath 2017; Revilla Minaya 2019; Shepard et al. 2010). We refer to these as the U.S. and Matsigenka datasets, respectively.

The U.S. dataset comprises data collected at regular intervals between birth and (at most) 21 years of age (shorter intervals before age two), with an average of 30 measurements per individual. The original authors then manually smoothed the data by taking a moving weighted average of the actual measurements (Tuddenham and Snyder 1954, pg 193–198). Additional extrapolated measurements reported in the original dataset are excluded. To aid in fitting the theoretical model, we duplicated each individual’s last reported height and weight measurements yearly until the age of 26. We also set each individual’s height to 0.012 cm and weight to 1.02*·*10^−6^ g (i.e., that of an egg cell: Leary et al. (2014); Hatton et al. (2023)) at nine months prior to birth (i.e., conception).

The Matsigenka dataset comprises data collected using an electronic balance (Tanita BC-351) and a stadiometer (Seca 213) from all individuals who were between two and 24 years of age and residing in the Matsigenka communities or attending one of three boarding secondary schools in neighboring communities at the time of our visits between 2017 and 2019. One Matsigenka community and the three boarding schools were visited twice in this period. Five one-year-old children who were eager to be measured are also included in the dataset. Due to our uncertainty about age in this population, ages are recorded to the nearest year. A maximum of three, and an average of 1.3, measurements per person were collected. The majority (70%) of Matsigenka children are represented by measurements at a single time point. As with the U.S. dataset, we set each individual’s height and weight to that of an egg cell at conception. Raw data are plotted in Appendix Figure A.6.

To obtain more accurate parameter estimates (Appendix D.1.3), we simultaneously fit the model to both height and weight data in the combined U.S. and Matsigenka datasets. Observed height *h*_*jt*_ and weight *m*_*jt*_ of person *j* at total age *t* years since conception are given by:

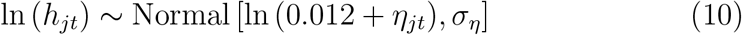

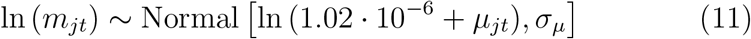

where *η*_*jt*_ and *µ*_*jt*_ are equations 8 and 9, respectively, which, when each parameter is indexed by *j*, represent height and weight trajectories of individual *j* in the five growth processes. These equations are added to the height (diameter) and weight of a human ovum. To represent the fact that observed heights and weights can never be negative, we take the log of both the observations and the mean of the Normal likelihood. The standard deviations *σ*_*η*_ and *σ*_*µ*_ represent measurement error, on the log scale, around individual *j*’s actual height and weight, respectively, at total age *t*. Females and males are fit with separate models.

We use a multilevel model design in a Bayesian framework (McElreath 2020), such that individuals’ height or weight trajectories in a given component growth function are estimated as the sum of ethnic group-level and person-level offsets to a baseline growth trajectory (i.e., random effects). We allow these offsets to covary across growth processes of an individual. Complete model structure, derivation, and priors are provided in Appendix D. Models were fit in R (R Core Team 2022) and Stan (Stan Development Team 2022) using the cmdstanr package (Gabry and Cesnovar 2021). Data and analysis scripts are provided at https://github.com/jabunce/bunce-fernandez-revilla-2022-growth-model.

Posterior parameter estimates from fit models are then systematically modified to simulate the effects on growth of specific healthcare interventions likely to affect metabolism.

## Results

### Theoretical model comparison

The composite model presented here was compared against the parametric JPA-1 model of Jolicoeur et al. (1992) (Appendix F.3) and the spline-based shape invariant SITAR model of Cole et al. (2010) (Appendix F.4) by fitting all three models to measurements of U.S. boys. As described above, the composite model was fit simultaneously to heights and weights (Appendix D.1.3), while the latter two models were fit only to height, as they are not designed to be fit to both data types simultaneously. Visual inspection suggests that estimates of the mean height trajectory by the JPA-1 and SITAR models are more similar to each other than they are to estimates of the composite model (Appendix Figure A.20).

However, estimates of all three models generally coincide well, giving us confidence that the composite model is appropriate for population-level comparisons of growth. Detailed model comparison and analysis of residuals is presented in Appendix G.1.

### Empirical estimates

Figure 2 shows differences in mean posterior estimates of population-level and individual-level height trajectories for U.S. and Matsigenka girls and boys between conception and adulthood. Weight trajectories from this model are shown in Appendix Figure A.9. Comparing trajectories of the five component processes across U.S. and Matsigenka girls and boys, the model predicts that population-mean differences in overall height are primarily the result of processes beginning during infancy and early childhood (and, for boys, in utero). Appendix Figure A.10 incorporates uncertainty in the estimated U.S. and Matsigenka mean growth trajectories in order to compute descriptive characteristics of these trajectories: maximum achieved height, maximum achieved growth velocity during puberty, and the age at maximum velocity during puberty. Analysis of residuals is presented in Appendix G.2.

**Figure 2.**
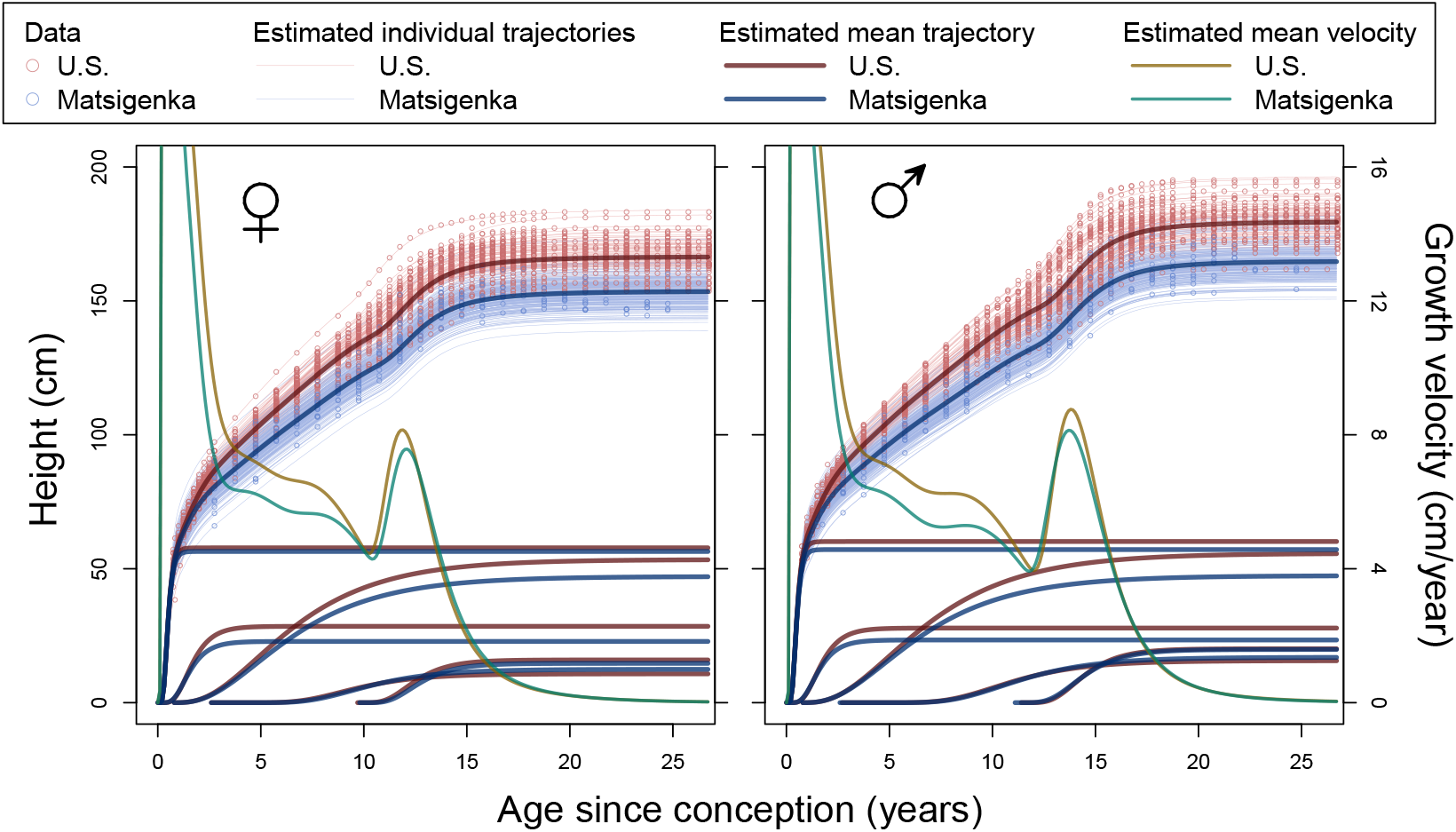
Posterior estimates from the composite growth model fit to U.S. (red) and Matsigenka (blue) height. Thin lines are mean posterior trajectories for each individual. Thick red and blue lines are mean posteriors for the mean cumulative trajectories of each ethnic group, as well as the mean trajectories for each of the five component growth processes. Corresponding mean posterior mean cumulative velocity trajectories in green and orange are decreasing (after age 14) from left to right.

Figure 3 presents a comparison of population-mean parameter estimates for U.S. and Matsigenka children summed for each age over the entire period of growth. For a given age, parameters in each of the five growth processes are weighted by the contribution of that process to overall height. Metabolic parameters *K* and *H* are further weighted by an expected decrease in metabolic activity as each process approaches its asymptote (Appendix D.4). This analysis suggests that, during most of ontogeny, Matsigenka girls and boys tend to have lower values of the allometric parameter *q* (i.e., smaller intestinal surface area, and concomitant body circumference, for a given height), and higher rates of catabolism (*K*) and anabolism (*H*) than U.S. children. These trends all reverse in the perinatal period. Objective values of metabolic parameters *K* and *H* shown in Figure 3 coincide reasonably well with estimates of catabolic and anabolic rates calculated independently from contemporary U.S. dietary intake data (Appendix D.5). Estimates of *i* parameters are shown in Appendix Figure A.8.

**Figure 3.**
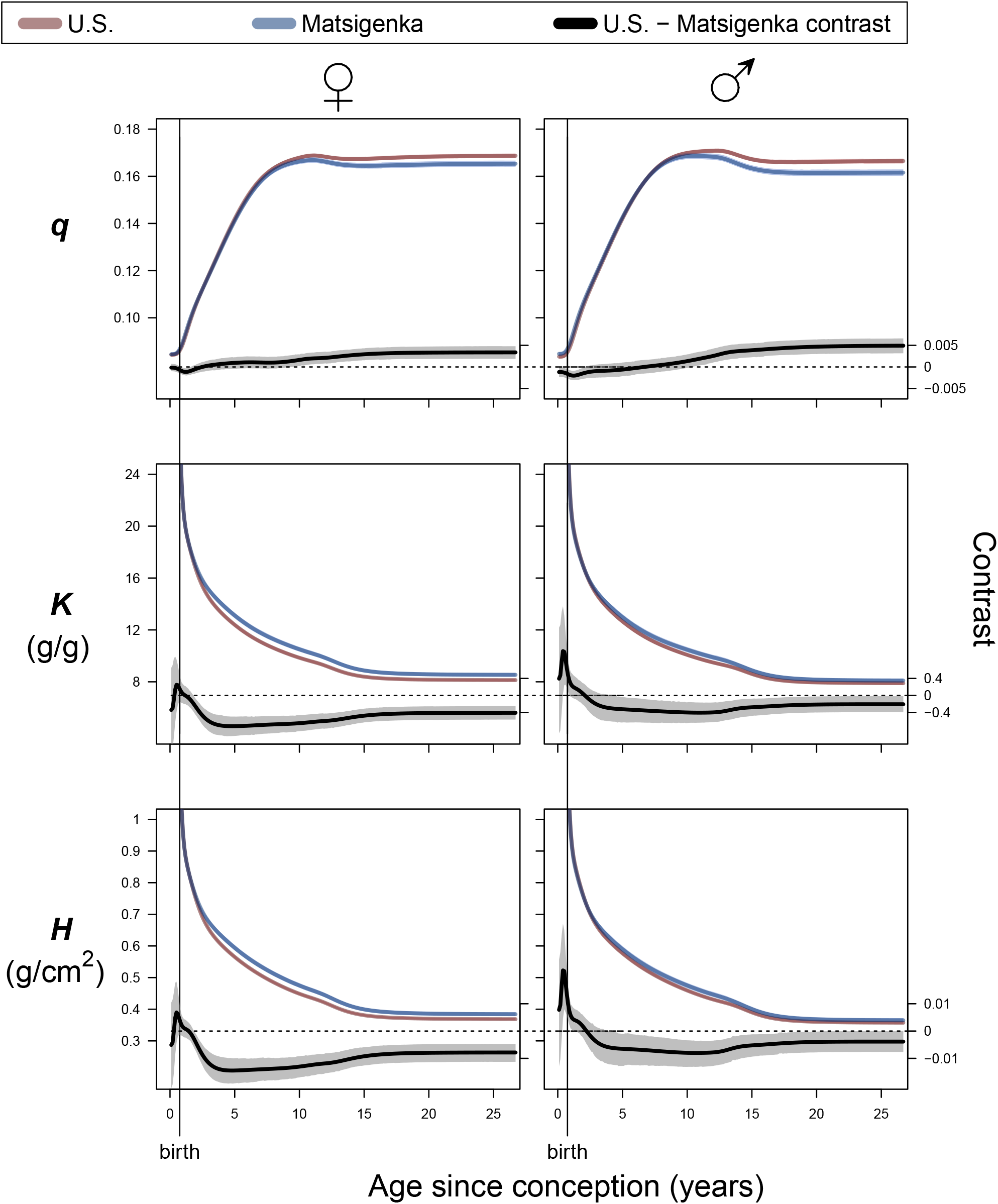
Estimates of mean parameter values by age for each ethnic group. Lines are means and shaded areas are 90% highest posterior density intervals (HPDIs: McElreath (2020)) for posterior distributions of sums of model parameters *q* (allometry), *K* (catabolism), and *H* (anabolism) across the five growth processes (Figure 1) at a given age, weighted by the contribution of each process to overall height and metabolism at that age (Appendix D.4), for U.S. (red) and Matsigenka (blue) children. HPDIs for parameter trajectories are often too narrow to be visible around the mean lines. The 90% HPDI of the U.S. - Matsigenka contrast (difference) is shown in grey. HPDIs of contrasts that do not overlap zero (dotted horizontal lines) indicate a detectable ethnic-group difference in parameter estimates.

### Intervention simulation

Figure 4 shows the predicted effects of two metabolic interventions on mean Matsigenka height trajectories. In the first intervention (second row of Figure 4), values of all Matsigenka metabolic parameters (*H*’s and *K*’s) are changed to match the mean estimates for U.S. children. This allows us to see the contribution of allometry (*q* parameters) to the observed mean height differences between Matsigenka and U.S. children.

**Figure 4.**
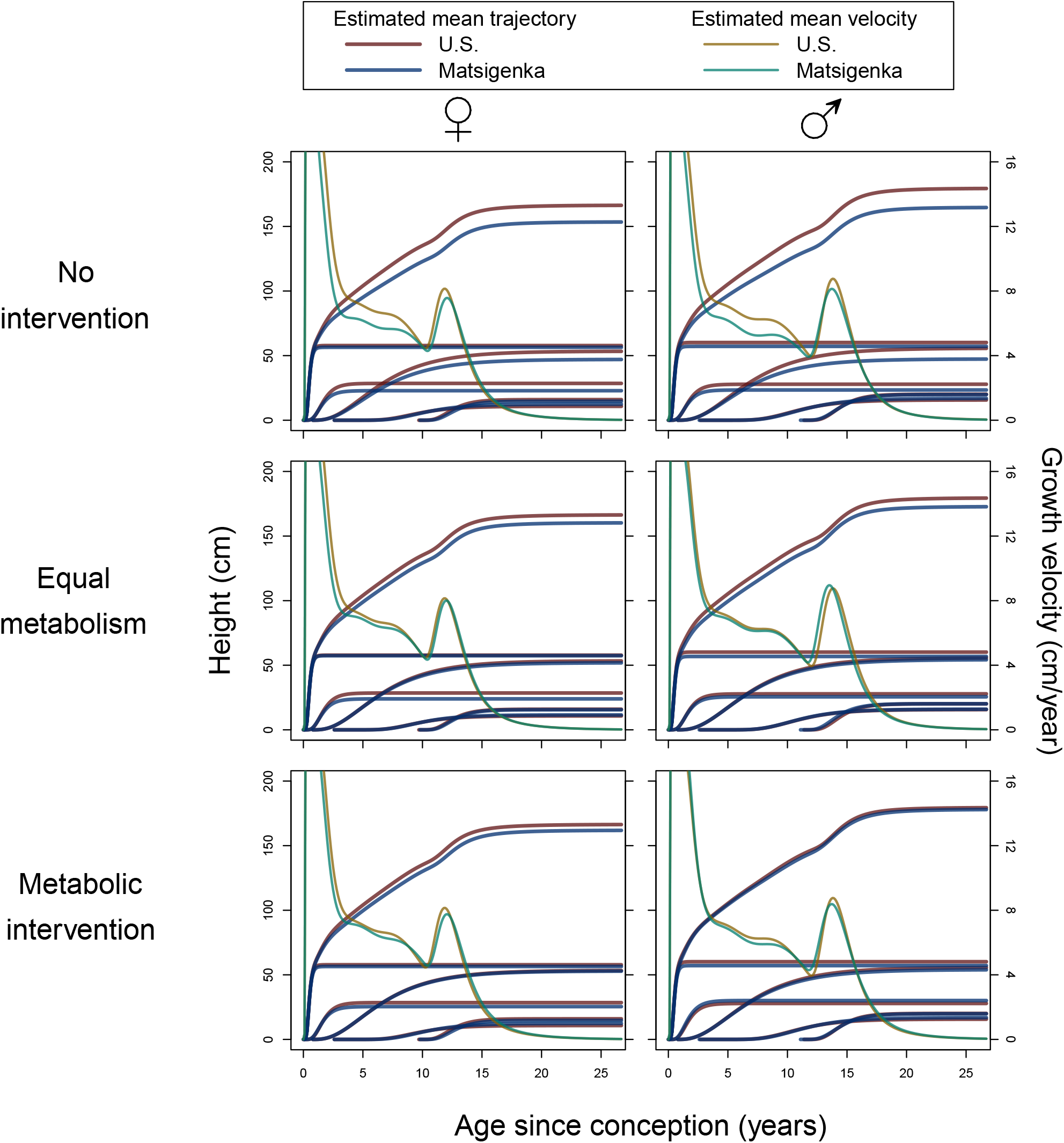
Simulated effects of healthcare interventions on Matsigenka mean height. Shown are mean posterior trajectories for mean overall height and five component growth processes for U.S. (red) and Matsigenka (blue) children. Corresponding mean growth velocities are decreasing (after age 14) from left to right. The first row reproduces Figure 2, and represents the pre-intervention state. In the second row, values of all metabolic parameters (*K*’s and *H*’s) for Matsigenka are modified to match mean estimates of those for U.S. children. In the third row, only Matsigenka values of *K*_2_ and *H*_3_ (girls) and *K*_2_ and *K*_3_ (boys) are modified to match mean estimates of those for U.S. children. Further analysis of interventions is presented in Appendix E.

The second is a targeted metabolic intervention (third row of Figure 4) meant to simulate dietary supplementation (presumed to increase *H*) and a decrease in immune system activation due to disease (presumed to decrease *K*) in the Matsigenka population during infancy and early childhood (the second and third component growth processes: Figure 1). As shown in Appendix Figure A.7, in the infancy process, both Matsigenka girls and boys are estimated to have higher mean values of *K*_2_ and *H*_2_ than U.S. children. Thus, we decreased *K*_2_ for Matsigenka girls and boys to match that of U.S. girls and boys, but did not modify *H*_2_. In the early childhood process, Matsigenka girls are estimated to have lower mean *H*_3_ and indistinguishable *K*_3_ compared to U.S. girls, while Matsigenka boys were estimated to have higher mean *K*_3_ and indistinguishable *H*_3_ compared to U.S. boys. Thus, we increased *H*_3_ for Matsigenka girls to match that of U.S. girls, and decreased *K*_3_ for Matsigenka boys to match that of U.S. boys. Because diet and disease are probably not solely responsible for population-level differences in average metabolism (e.g., genes may also contribute), these simulated changes likely represent an upper limit on the effect of an intervention targeting diet and disease.

Comparing maximum achieved height in the first row of Figure 4 against that in the last two rows suggests that, under both interventions, Matsigenka children would grow taller. However, with the possible exception of boys in the second intervention, Matsigenka children are still predicted to be shorter than U.S. children, on average. This suggests that allometric (i.e., body shape) differences, represented by the *q* parameters, are at least partly responsible for observable mean height differences between these two populations. Appendix E presents additional analyses of these interventions, including their effects on weight, and shows how modifying metabolic parameters *H* and *K* can contribute to changing overall body allometry, as well as the age at maximum growth velocity, even if *q* and *i* parameters are unmodified.

## Discussion

We have developed a new causal model of human growth and estimated its parameters using temporally-dense data from a population of U.S. children, and temporally-sparse data from a population of Indigenous Matsigenka children. The model predicts that these two ethnic groups differ in terms of overall mean rates of anabolism and catabolism, and the allometric relationship between body height and intestinal surface area. Furthermore, analysis of the model suggests that both metabolism and allometry contribute to observed mean size differences between the populations. Below we present possible explanations for these differences, implications for health interventions, limitations of this new approach, and future directions.

### General trends

Figure 3 suggests that total rates of catabolism (*K*) and anabolism (*H*) are very high at conception and then decrease toward an asymptote by age 20 (resembling a horizontal reflection of height growth: Figure 2). This general pattern can be compared to the findings of Pontzer et al. (2021), who use a doubly labeled water method with international participants of a range of ages to calculate size-adjusted total daily energy expenditure (TEE), conceptually similar to catabolic rate *K*. Adjusted TEE at birth resembles that of adults, but increases rapidly during the first year, after which it declines toward an asymptote at age 20. At birth, the relatively low estimate of TEE contrasts with our relatively high estimate of *K*. This could result from a (plausibly adaptive: Vasak et al. 2015) slowing (negative acceleration) of fetal growth prior to birth (Kiserud et al. 2018), which would lower perinatal estimates of TEE and *K*. To represent such a slowing, and a lower *K* (and *H*) at birth, the new composite model could be modified to include more than one in utero component growth process, and then fit to a dataset that includes fetal (and placental) size measures.

### Inter-group comparisons

Figure 3 suggests that, just prior to birth, Matsigenka tend to have lower (boys) or indistinguishable (girls) average rates of catabolism (*K*) and anabolism (*H*) compared to U.S. children. However, by approximately two to five years after birth, Matsigenka metabolic rates are estimated to be higher than those of U.S. children, a pattern that persists into adulthood. Perinatal comparisons must be interpreted with caution, as in utero and infant component growth functions are fit to very few measurements of Matsigenka children younger than two years (Appendix Figure A.6), and studies in other populations have found little cross-cultural variation in weight-adjusted infant energy expenditure (Prentice and Paul 2000). However, with this caveat in mind, the marked postnatal increase in Matsigenka metabolic rates compared to U.S. children could result, in part, from Matsigenka infants consuming more energy during the first years of life, perhaps relating to the timing and type of supplemental foods introduced during infancy, practices that were changing rapidly in the U.S. of the late 1920s (Bentley 2014; Castilho and Barros Filho 2010; Stevens et al. 2009). This hypothesis requires future systematic study. At the same time, Matsigenka infants may also expend relatively more energy due to immune system activation in response to parasites or other pathogens in the tropical forest environment (Urlacher et al. (2018); Gurven et al. (2016); Garcia et al. (2020), though see Urlacher et al. (2021)), e.g., intestinal pathogens accompanying the introduction of solid food.

The higher mean Matsigenka rates of catabolism (*K*) relative to U.S. children are estimated to continue through ontogeny (Figure 3). As with infants, this could be due to Matsigenka immune system activation in response to tropical pathogens and parasites, which is associated with shorter stature in other Amazonian Indigenous populations (Urlacher et al. 2018; McDade et al. 2008). Mean rates of anabolism (*H*) are also estimated to be higher for Matsigenka than for U.S. children. This suggests that Matsigenka synthesize more mass per unit intestinal surface, which, for a given intestine size, can be achieved through consumption of more energy-dense foods. This is unexpected, given that Matsigenka children’s foods are produced through their parents’ daily fishing, hunting, gathering, and horticultural activities, that, on average, likely have more fiber and lower energy density than foods available to upper-class U.S. children in the 1930s, a time when commercially processed foods were gaining in popularity (Bentley 2014). One potential explanation is that Matsigenka have greater intestinal surface area for their body size than do patients in Euro-American clinics from whom our surface area calculations are derived (Appendix D.1.2). Indeed, Weaver et al. (1991) reports considerable variation in human small intestine length for a given height, which suggests that intestinal surface area may adjust during ontogeny as a plastic response to disease and dietary changes. Underestimating Matsigenka intestinal surface area *s* would lead to an overestimate of the anabolic rate *H* (equation 1). This hypothesis requires future study.

Figure 3 suggests that, starting around age 10, Matsigenka children have lower values of the allometric parameter *q* than do U.S. children. Because *r* = *h*^*q*^ (equation 4), this is interpreted to mean that, for a given height, Matsigenka have a narrower skeleton, defining a smaller abdominal cavity, housing an intestine with smaller surface area (Appendix D.1.1). This result coincides with our impression that Matsigenka tend to have ectomorphic bodies. However, it is unexpected given the presumably lower energy density (e.g., higher fiber content) of typical Matsigenka foods, and the negative relationship between gut surface area and dietary energy density (fauna *>* fruit *>* foliage) observed in other mammals (Chivers and Hladik 1980). Underestimation of the Matsigenka *q* parameter could be produced through a disproportionate underestimation of Matsigenka weight (Appendix G.3), or, as above for *H*, an underestimation of Matsigenka intestinal surface area for a given body size. A future challenge will be to determine (noninvasively) whether populations like the Matsigenka tend to surpass urban populations in intestinal surface area per abdominal volume.

### Healthcare interventions

Figure 2 suggests that observed differences in stature between Matsigenka and U.S. children are primarily the result of component growth processes that make their most substantial contributions to overall height in utero, during infancy, and during early childhood (i.e., the first three component growth processes). This suggests that, if such differences signal growth-related health challenges, interventions should be targeted toward children in these earlier developmental stages. This highlights an advantage of process-partitioned composite growth models, like that developed here (as well as the ICP (Karlberg 1989) and QEPS models (Nierop et al. 2016; Holmgren et al. 2022)): they allow exploration of the ages at which population-level differences in growth contribute to observed size differences. This is not possible with other growth models (e.g., JPA-1: Jolicoeur et al. (1992) and SITAR: Cole et al. (2010)), that represent growth over all of ontogeny as a single process (Mansukoski et al. 2019). A unique advantage of the causal theoretical model developed here compared to previous models is the ability to simulate interventions, which can inspire hypotheses about underlying causes of growth variation. For example, Figure 4 suggests that the shorter mean stature of Matsigenka children relative to U.S. children is partly due to metabolic differences between these populations. These may be responsive to environmental manipulation. For instance, Matsigenka are predicted to grow taller after an intervention that: 1) increases anabolic rates (*H*), e.g., through the provision of supplemental foods dense in energy and substrates for corporal construction; and 2) decreases catabolic rates (*K*), e.g., through disease treatment and prevention to decrease immune system activation. Importantly, however, this analysis suggests that allometry (*q*) also contributes to height differences between Matsigenka and U.S. children, as simulated elimination of average metabolic differences is not predicted to equalize their average heights (Figure 4). Compared to metabolism, allometry (body shape) may be under stronger genetic control, and thus more responsive to drift and natural selection (Katzmarzyk and Leonard 1998) at the population level on generational time scales. For instance, there is genetic evidence that short stature may provide a selective advantage in tropical forest environments (Perry et al. 2014; Perry and Verdu 2017; Zoccolillo et al. 2020), though the nature of any such height adaptation, or linkage to another adaptive trait, is still poorly understood (Perry and Dominy 2009). If allometric differences of genetic origin contribute substantially to population differences in average height, it would lend support to recent calls for caution (Hackman and Hruschka 2020; Hruschka 2021) when diagnosing stunting, a form of malnutrition defined as low height for age, by comparing children’s growth against a single universal human standard, as is currently the norm in international contexts (de Onis and Branca 2016; WHO Multicentre Growth Reference Study Group and de Onis 2006).

Interestingly, interventions on metabolic rates can affect overall body shape, even if the allometric *q* parameters of each component growth process are unchanged (e.g., under complete genetic control). Appendix Figure A.11 shows that a simulated intervention to equalize mean Matsigenka and U.S. metabolic rates results in nearly identical mean weight trajectories, even while height trajectories remain distinct (Figure 4). This suggests that the intervention would cause Matsigenka to become heavier for a given height. Appendix Figure A.12 shows that the simulated intervention has indeed increased overall Matsigenka *q*, and by extension intestinal surface area and body circumference for a given height, above that of U.S. children. This effect is due to the fact that metabolic rates affect the shapes of the five component growth functions, and thus their weighted contributions to the overall value of *q* at a given age (Appendix E). Metabolism-induced changes in the shape of component growth functions result in a similar effect on the timing of growth: the simulated metabolic intervention decreases Matsigenka age at maximum pubertal growth velocity (compare Figure A.10 and Appendix Figure A.14), despite the fact that initiation parameters *i* for each component growth process are unchanged. These effects parallel Tanner et al. (1982)’s analysis of a secular trend whereby growth patterns of cohorts of Japanese children changed over the course of 20 years, during which living (and presumably nutritional and metabolic) conditions improved: children’s legs (but not trunk) grew longer, and they reached maximum pubertal growth velocity earlier. In the context of our new model, we would hypothesize that improving living conditions in Japan increased anabolism (*H*) and decreased catabolism (*K*) in a component growth process that contributed more to leg length than to expansion of the abdominal cavity (and intestinal surface area), and thus had a low value of the allometric *q* parameter. Changing the shape of this component process (i.e., increasing its contribution to overall growth), relative to that of the other processes, resulted in a shallower overall *q* trajectory and an earlier timing of maximum pubertal growth.

### Limitations and future directions

In developing this theory of human growth, we make a number of simplifying assumptions to facilitate derivation and fitting of the mathematical model. These assumptions, detailed in Appendices A–D, place important limits on interpretation of model estimates, and, at the same time, serve as a guide for future work. For instance, in the absence of cross-cultural data, we assume a constant and universal relationship between skin surface area and intestinal surface area (Appendix D.1.2), which, as explained above, may distort estimates of *H* and *q* for populations like the Matsigenka. We represent components of the body as cylinders (Appendix Figure A.1), the radii and heights of which are constrained to grow according to a simple power function (*r* = *h*^*q*^, equation 4). While better than alternative simple functions (Appendix Figures A.2 and A.3), this assumption precludes any change in radius (and by extension, weight) resulting from soft tissue (e.g., fat and muscle) development that does not correspond to a height increase. When fitting the model simultaneously to both height and weight measurements, we compensate for this assumption by defining unequal constraints on variance due to measurement error between the two data types (Appendix D.1.3). Appendices F.1 and F.2 present alternative, less satisfactory approaches. A mechanistic theory that explicitly incorporates soft tissue growth would improve on the present model.

In computing overall mean parameter values by age (Figure 3), we assume that metabolic rates decrease as a component growth process approaches its asymptote, primarily due to replacement of metabolically active red marrow cells by less active marrow fat cells as the skeleton ages. In the absence of data, the proportion by which metabolic rates decrease at the asymptote is arbitrarily set to 0.75 (Appendix D.4). Future work is needed to determine if, and by how much, mean rates of anabolism and catabolism change with the replacement of such cell types during ontogeny.

In this study, Matsigenka children are represented by a temporally-sparse dataset (Appendix Figure A.6) typical of those collected in remote non-urban societies often neglected in clinical growth studies. While the Bayesian analytical techniques employed here facilitate estimation of growth trajectories from sparse datasets, more longitudinal data usually result in more accurate estimates. Thus, it will be of interest to know if and how model estimates of Matsigenka growth parameters change as more data are collected in coming years. Given the sparse dataset, and the many assumptions of this new growth model that have yet to be systematically explored, the above comparison of U.S. and Matsigenka children’s growth should not be viewed as a definitive analysis upon which to base healthcare decisions. Rather, it illustrates the promise and potential of this new approach to serve as a framework to: 1) relate height, weight, and intestinal growth to body allometry and rates of anabolism and catabolism; 2) determine the relative contributions of metabolism (likely strongly affected by diet and disease) and allometry (likely more strongly affected by genetic factors) to observed size differences between individuals and populations; 3) simulate the effect of healthcare interventions on children’s growth; and 4) complement the strengths of existing growth models, such as SITAR (Cole et al. 2010), QEPS (Nierop et al. 2016), and JPA-1 (Jolicoeur et al. 1992), by providing a theoretical foundation for the design of desired healthcare support strategies better tailored to the needs of particular communities, and, importantly, to guide exploration of the causes of growth variation in our species.

## Ethics statement

Research was conducted with authorization from the Servicio Nacional de Áreas Naturales Protegidas por el Estado del Perú (SERNANP). Verbal informed consent from adult Matsigenka participants, and informed assent from their minor children, was obtained following standards for the protection of research subjects implemented by the Department of Human Behavior, Ecology, and Culture at the Max Planck Institute for Evolutionary Anthropology, with approval from the Ethics Council of the Max Planck Society (Application No. 2019 27). Results were presented to, and discussed with, Matsigenka participants in community-wide meetings prior to publication.

## Acknowledgments

We thank participants in the Matsigenka Native Communities of Tsirerishi, Tayakome, Sarigeminiki, and Yomibato. M. Navarra of Chaska Wasi (Salvación), P. Rey and C. Llana of the Misión san Miguel Arcangel Shintuya, E. Herrera, K. Mejía, and M. Chinchiquiti of Maganiro Matsiguenka (Boca Manu), N. Oyeyoyeyo, R. Chiqueti and family of Tayakome, C. Huamantupa, J. Poma and family of Atalaya, V. Chávez, C. Flores, and F. Rayan of the Cocha Cashu Biological Station, N. Santullo and C. Matos of Rainforest Flow, J. Florez and E. Meza of SERNANP Cusco, O. Espinosa of the Pontificia Universidad Católica del Perú, M. Minaya, O. Revilla, N. Revilla, G. Lugon, and L. Revilla provided support with fieldwork. S. Atmaca, B. Beheim, A. Büchner, and A. Bublikova helped with data transcription. C. Ross, A. Dalla Ragione, R. McElreath, B. Beheim, D. Lukas, W. Church, E. Ready, N. Blurton-Jones, T. Cole, M. Guilfoyle, J. Cullin, A. Wiley and members of the Department of Human Behavior, Ecology, and Culture (HBEC) at the Max Planck Institute for Evolutionary Anthropology provided critique on the theoretical model, statistical analyses, and/or earlier versions of this paper. All errors are ours. Funding was provided by the Max Planck Society.

## APPENDICES

### Appendix A. Derivation of the height growth model

We require a geometrical representation of the human body that incorporates both height and intestinal surface area, and is simple enough to make Equation 1 in the main text solvable. We employ the special case where an everted tube becomes a solid cylinder of length *h* and radius *r*, such that intestinal surface area is represented by the external surface of the cylinder (minus the circular ends), and the length of the cylinder represents height (Figure A.1). Incorporating this geometry into Equation 1 yields Equation 2 in the main text. Solving the latter for the longitudinal growth rate 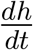 yields Equation 3 in the main text.

**Figure A.1.**
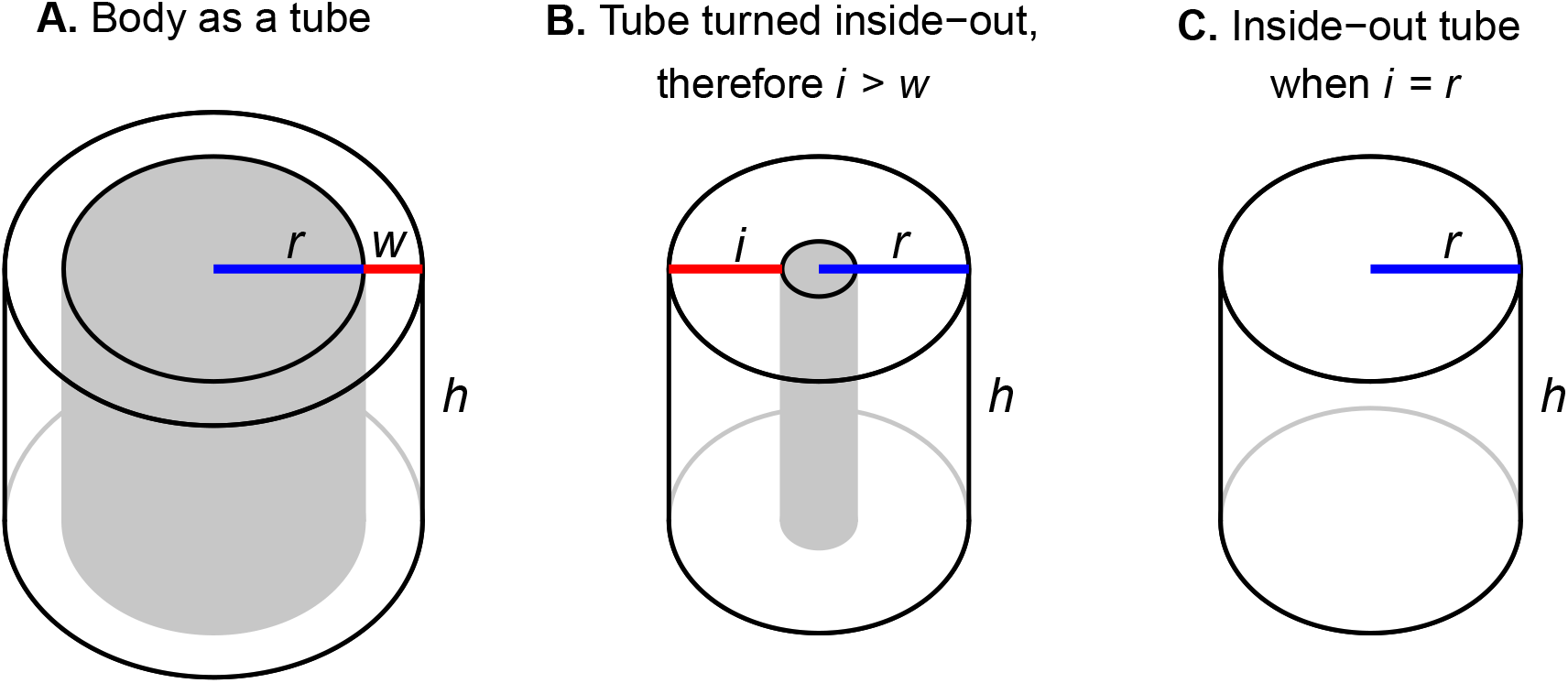
Approximating the geometry of the human body. **A)** The body (or a part thereof) is represented as a tube of length (i.e., height) *h*. The nutrient-absorption surface (i.e., intestine) is on the inside and has radius *r*. The volume of the body is contained in the wall of the tube, which has thickness *w*. In reality, due to villi and microvilli, the absorbing surface area of the human digestive tract is more than an order of magnitude larger than the external (skin) surface area (Mosteller 1987; Helander and Fändriks 2014). Thus, a tube, where internal surface area is necessarily less than that of the exterior, is an imperfect representation (see Appendix D.1.2). **B)** The tube is everted (turned inside out), such that the absorbing surface is now on the outside. If *r* and the solid volume are held constant, this requires the thickness of the new wall *i* to increase relative to *w*, such that 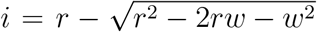. **C)** For convenience in the analysis that follows, we assume the special case where 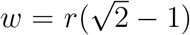, such that *i* = *r*, and the body can thus be approximated by a cylinder with height *h*, radius *r*, absorbing surface area 2*πrh* (i.e., not including the surface area of the cylinder’s ends), and volume *πr*^2^*h*. This implements an assumption that any increase in solid body volume (i.e., mass, given constant density) must be accompanied by a corresponding increase in absorbing surface area.

The theory of Pütter (1920) and von Bertalanffy (1938) makes different predictions depending on how an organism changes shape (or not) as it grows. If the cylindrical organism grows in only one dimension, e.g., it grows in length *h* but not in diameter, we can set radius *r* as a constant, which, from Equation 3, yields the rate of length growth:

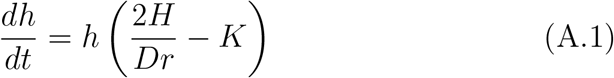

Because *H, D, r*, and *K* are assumed to be constant over time, and can therefore be consolidated into a single constant *B*, this equation has the form:

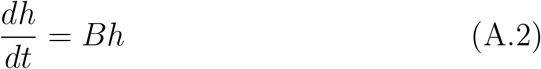

whose solution is the exponential function:

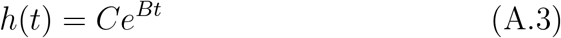

where *C* is a constant of integration. Thus, under the above assumptions, such an organism would grow continuously and exponentially in length, quickly becoming very long while retaining the same diameter. Now assume that the cylindrical organism grows in size and maintains the same shape (i.e., body proportions). In other words, the ratio of its radius *r* to its length *h* at any time *t* is a positive constant *F*. Setting *r*(*t*) = *F* · *h*(*t*), and substituting into Equation 3, yields the rate of length growth:

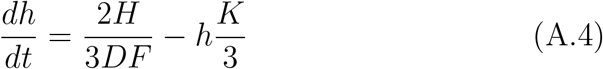

Consolidating constant terms into constants *G* and *J*, this rate has the form:

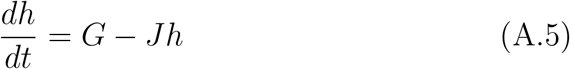

whose solution is the negative exponential function described by von Bertalanffy (1938):

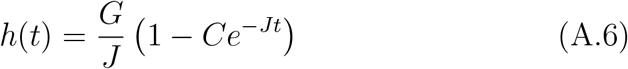

where *C* is a constant of integration, and 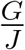 is a constant representing the asymptotic (e.g., maximum achieved) length. Thus, under the above assumptions, such an organism would grow at an ever-decreasing rate while maintaining the same shape, approaching a maximum size of length 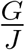 and radius 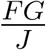. As was recognized early on (Pütter 1920), such a pattern of growth characterizes many fish species, and this model continues to be used as an excellent representation of fish growth (Essington et al. 2001; Vincenzi et al. 2020). One hypothesis for this model’s success is that a fish’s nutrient absorption surface (e.g., intestine) increases, like its external surface, at a constant proportion of the fish’s mass. In other words, the body proportions of most fish remain relatively constant between hatching and adulthood.

The growth of humans (and many other animals) falls between these two extreme patterns: The body changes in both size and shape over time, and no dimension can be reasonably assumed to be constant. Here, we are interested in five overlapping processes that affect growth in different ways at different ages during ontogeny (Appendix C). We assume that each of these growth processes contributes to changing overall body shape in a specific way, namely, (as a rough approximation) height and circumference (=radius*2*π*) both increase, but the ratio of circumference to height decreases over time, such that the body elongates.

Evidence for this comes from a comparison of height to both head circumference (Figure A.2) and waist circumference (Figure A.3). Note that, in these figures, circumference increases over time. Therefore *r* is not a constant, and, under the assumption that intestinal surface area is proportional to body circumference, we should not expect to observe pure exponential growth in height as described by equation A.3. Similarly, the ratio circumference/height decreases over time. Therefore 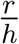 is not a constant *F*, and, under the same assumptions of intestinal surface area, we should neither expect to see pure negative exponential growth in height as described by equation A.6.

**Figure A.2.**
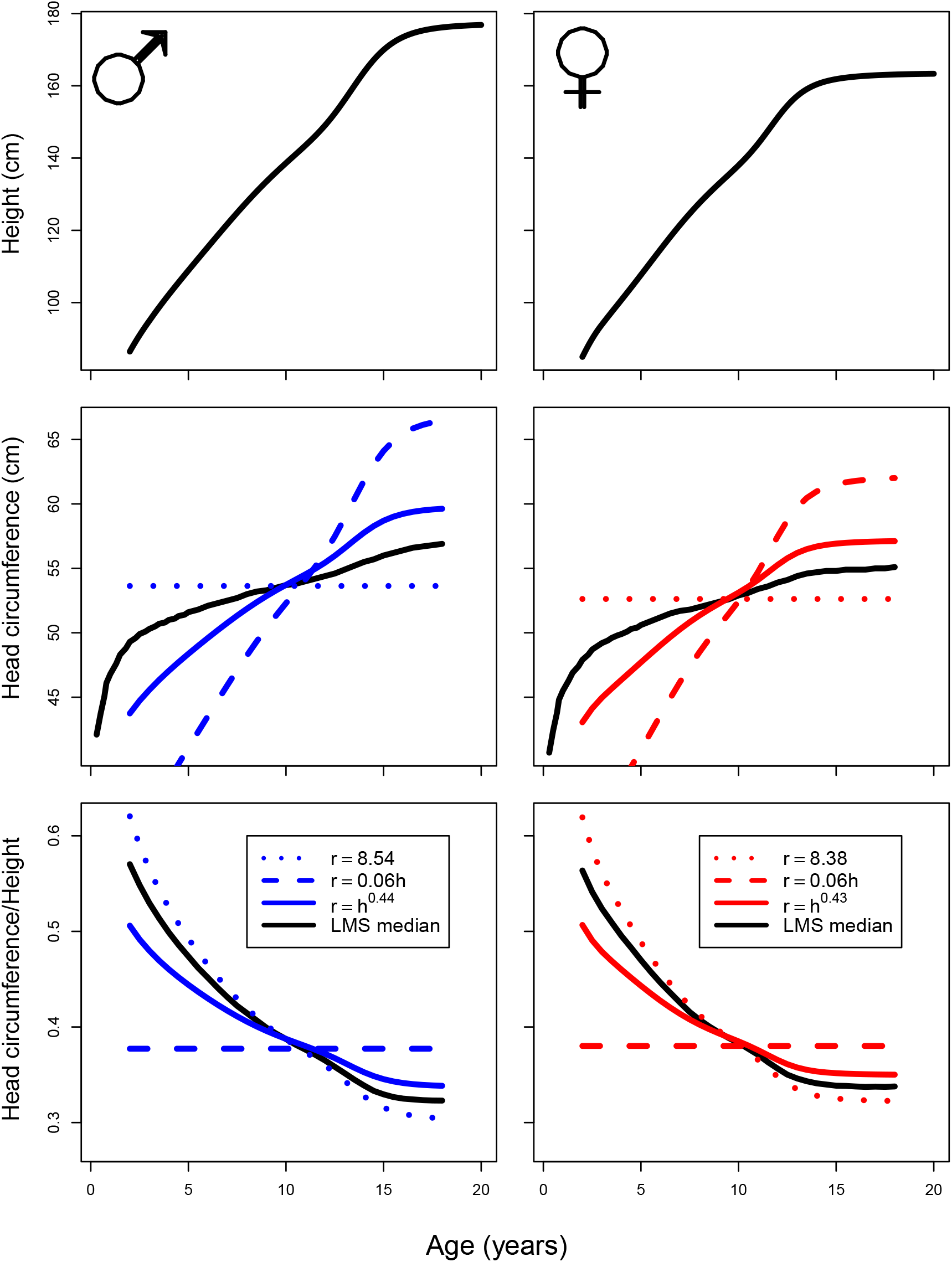
Changes in head circumference and height by age since birth. Reference trajectories of median height by age (top row) are taken from LMS values published by the CDC (https://www.cdc.gov/growthcharts/percentile_data_files.htm), derived from a cross-sectional sample of U.S. children. Reference trajectories of median head circumference by age (middle row) are taken from the LMS values of Schienkiewitz et al. (2011), from a nationally representative cross-sectional sample of German children. All reference trajectories are shown as black lines. Colored lines show predicted trajectories of head circumference and head circumference/height given median height trajectories, assuming the body is a cylinder with radius *r* and height *h*, and linear (height) growth is determined by Equation 3 in the main text, under three alternative relationships between *r* and *h* (dotted, dashed, and solid blue or red lines). Head circumference/height (bottom row) is calculated by dividing the respective median values for each age, thereby implementing a simplifying assumption that children with median height also have median head circumference. Functions relating head circumference to height were fit to simulated median head circumference trajectories using Rstan (Stan Development Team 2018). Data and analysis scripts are available at https://github.com/jabunce/bunce-fernandez-revilla-2022-growth-model.

**Figure A.3.**
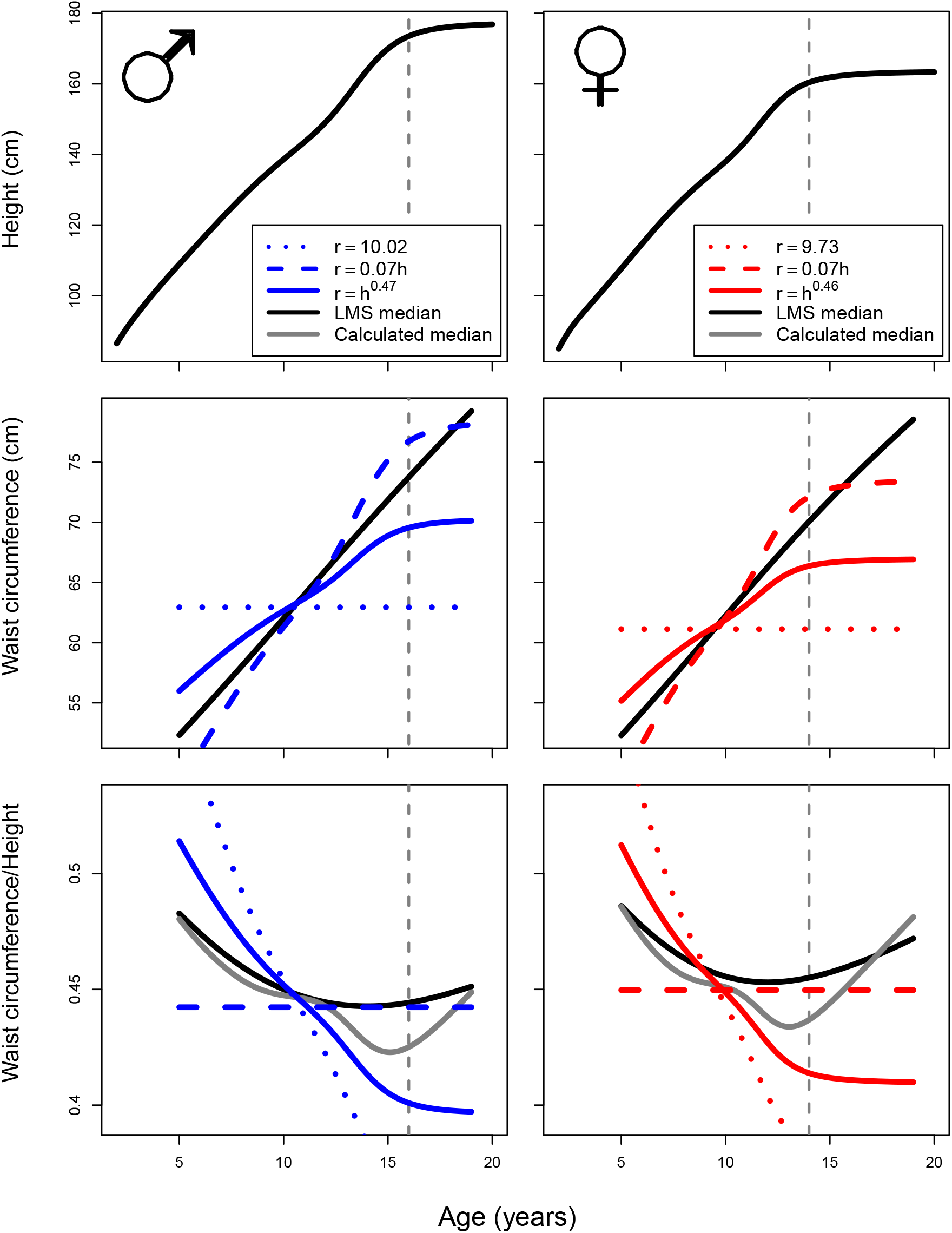
Changes in waist circumference and height by age since birth. Reference trajectories of median height by age (top row) are taken from LMS values published by the CDC (https://www.cdc.gov/growthcharts/percentile_data_files.htm), derived from a cross-sectional sample of U.S. children. Reference trajectories of median waist circumference (middle row) and median waist circumference/height (bottom row) by age are taken from the LMS values of Sharma et al. (2015), using CDC data. All reference trajectories are shown as black lines. Colored lines show predicted trajectories of waist circumference and waist circumference/height given median height trajectories, assuming the body is a cylinder with radius *r* and height *h*, and linear (height) growth is determined by Equation 3, under three alternative relationships between *r* and *h* (dotted, dashed, and solid blue and red lines). In the bottom row, observed median waist circumference/height is shown as a black line. Grey, blue, and red lines represent waist circumference/height calculated by dividing the respective median values for each age, thereby implementing a simplifying assumption that children with median height also have median head circumference. Functions relating waist circumference to height were fit (using Rstan (Stan Development Team 2018)) to simulated median waist circumference trajectories only until ages 14 and 16, for girls and boys, respectively (vertical dotted lines). After these ages, observed waist circumferences diverge notably from the predictions of all functions. Data and analysis scripts are available at https://github.com/jabunce/bunce-fernandez-revilla-2022-growth-model.

Given our simplifying assumption of a cylindrical human body (Figure A.1), *r* represents the intestinal diameter, intestinal length is equal to height *h*, and thus intestinal surface area is equivalent to skin surface area (an unrealistic assumption, requiring a conversion factor explained in Appendix D.1.2). Our objective here is to represent the general relationship between intestinal surface area and height as both increase. Specifically, we hypothesize that height *h* and intestinal surface area (represented as 2*πrh*) both increase during childhood, but that the proportional increase in intestinal surface from one time step to the next is less than the square of the proportional increase in height. This might occur, for instance, if, during some phases of growth, the body elongates, such that leg length increases without a substantial increase in intestinal surface (likely related to trunk volume)(see Eveleth and Tanner (1990), pg 186). In the cylindrical representation, this occurs if *r* increases slower than *h*, analogous to the relationship between height and body radius (circumference) in Figures A.2 and A.3.

A power function is a simple representation of a relationship between height and radius, where a power can be chosen such that both increase over time, but the ratio 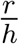 decreases over time. This is illustrated in Figures A.2–A.4.

**Figure A.4.**
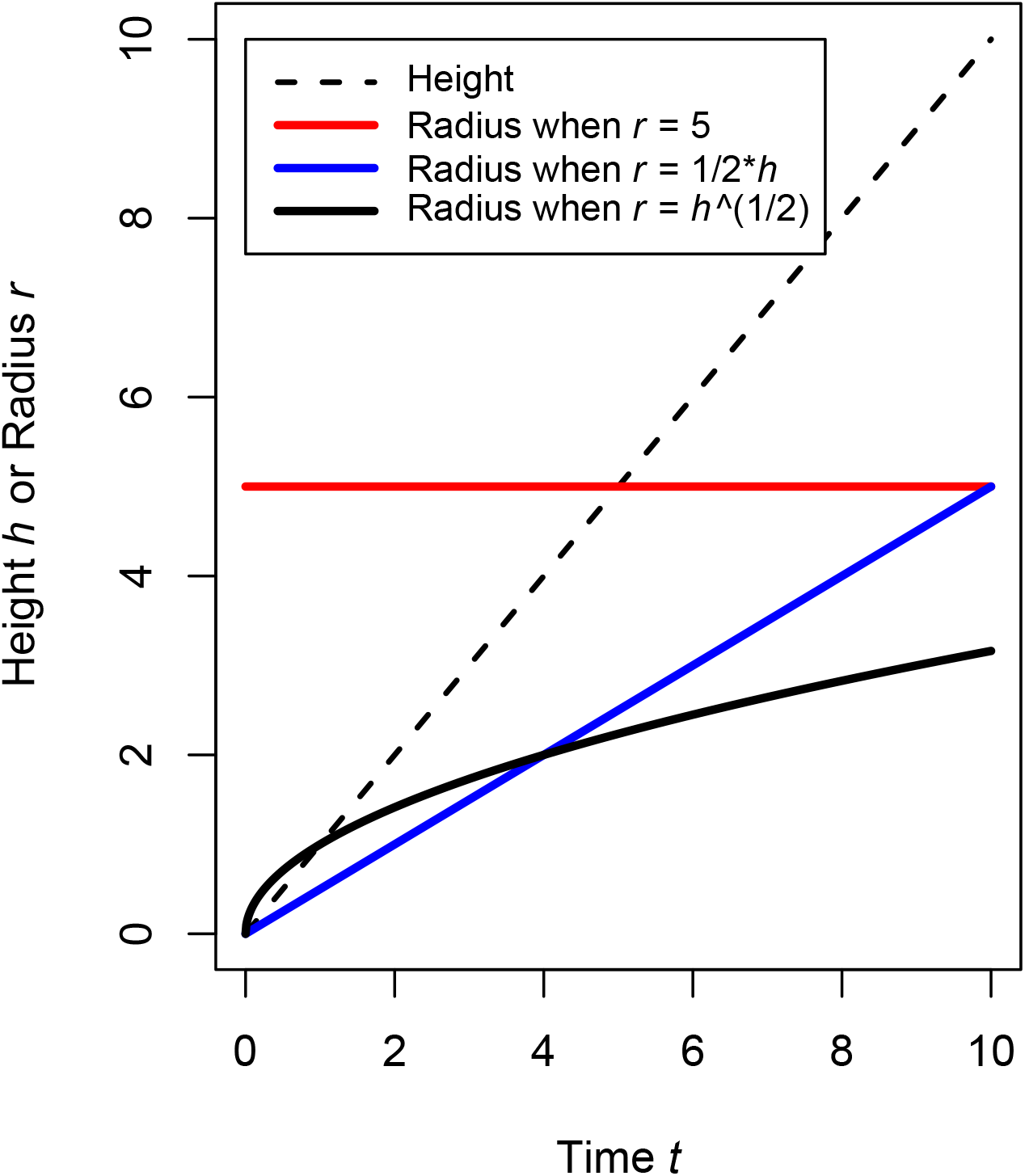
The effects of different relationships between radius *r* and height *h*. When *r* is a power function of *h* (such that the power is between 0 and 1), both *r* and *h* increase over time (unlike when *r* is a constant), while the ratio 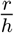 decreases over time (unlike when *r* is a constant multiple of *h*), such that the cylindrical body elongates.

To implement this assumption, we set *r*(*t*) = [*h*(*t*)]^*q*^, where 0 *< q <* 1. Substituting for *r* in Equation 3 yields Equation 4 in the main text. Substituting constants 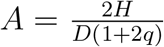 and 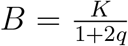, Equation 4 has the form:

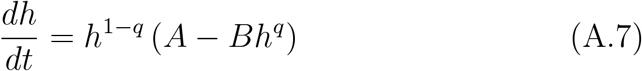

This is solved as follows:

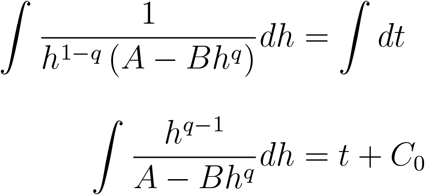

where *C*_0_ is a constant of integration. Substituting *u* = *A* − *Bh*^*q*^, and therefore 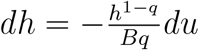, yields

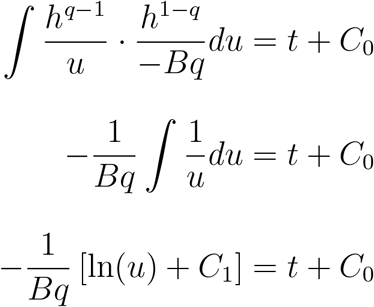

Re-substituting for *u* yields

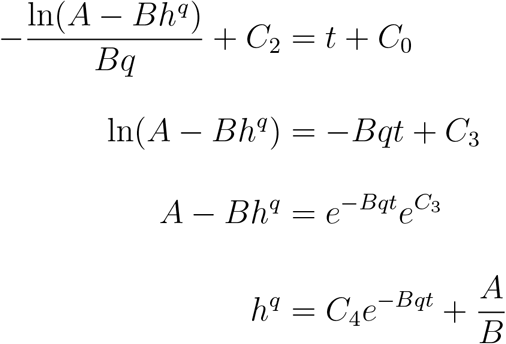

Re-substituting for *A* and *B* yields

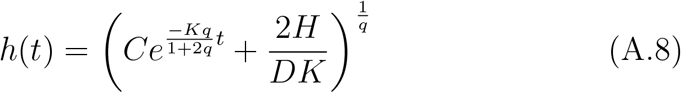

where *C*’s are modified constants of integration. This height function has the following derivatives:

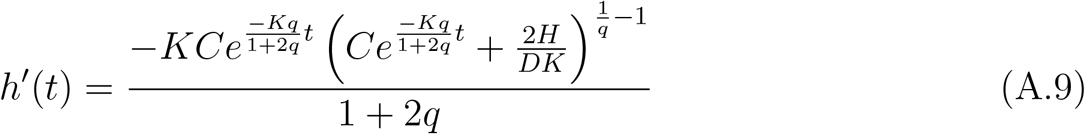

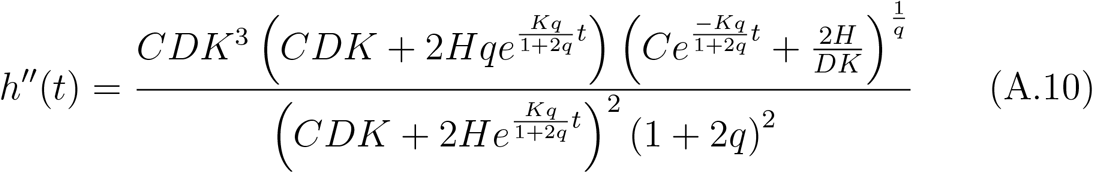

and the following characteristics:

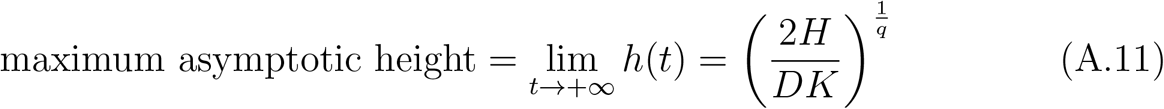

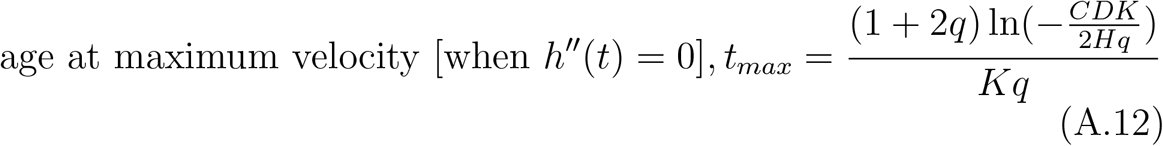

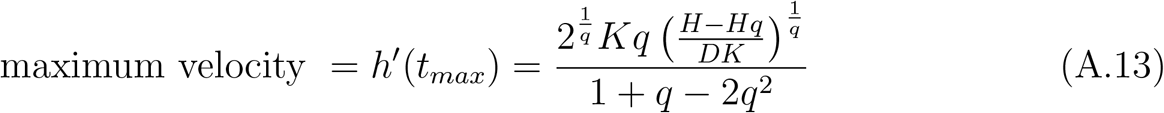

under the conditions that 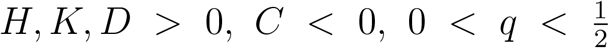, and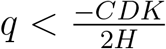. Note that changing the value of the constant *C* serves only to shift the function horizontally on the *x*-axis, i.e., it contributes only to changing the age at maximum growth velocity, not the shape of the growth function.

We define the constant *C* as follows:

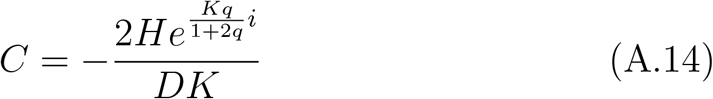

where *i* is the age at which the growth function initiates, i.e. when *h*(*t*) = 0. Substituting this value of *C* into equation A.8, and assuming, for simplicity, that density *D* = 1 g/cm^3^, yields equation 6 in the main text.

## Appendix B. Derivation of the weight growth model

The mass of a cylinder is density times volume = *Dπr*^2^*h*, where *h* is defined as equation 6, and also 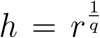 (from the definition of *q*).

Therefore

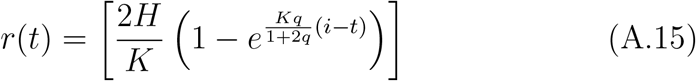

from which it follows that

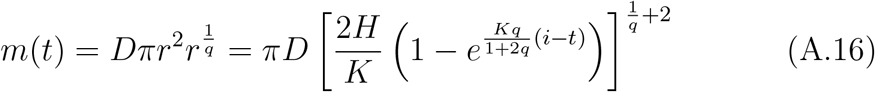

which, under the assumption that *D*=1 g/cm^3^, is equation 7. However, equivalently, we may also derive equation 7 directly from equation 1 as follows:

Using Equation 1 to approximate the rate of increase in mass over time, we again assume the organism’s shape can be roughly approximated by a cylinder of radius *r* and length *h* (Figure A.1). If *D* is density, then the cylinder has mass

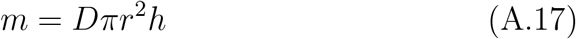

and surface area

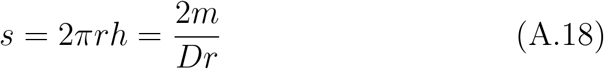

such that, like the height function, at any time *t*, 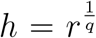, where 0 *< q <* 1. Substituting for *h* and solving Equation A.17 for *r* yields

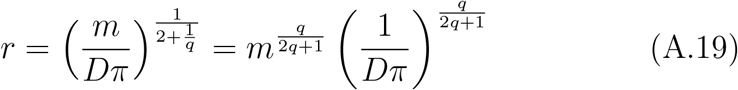

Substituting for *r* in Equation A.18 yields

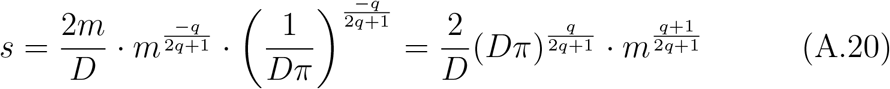

Substituting *s* into Equation 1 yields Equation 5 in the main text.

Substituting constant 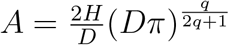, Equation 5 has the form

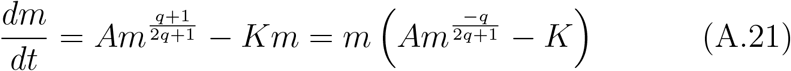

This is solved as follows:

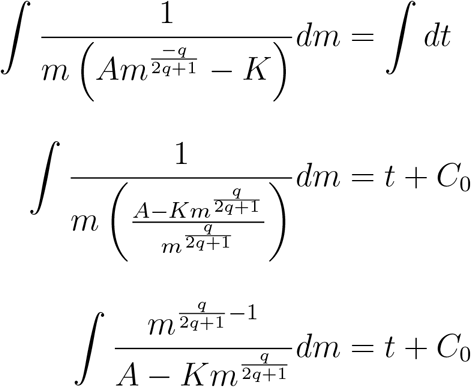

where *C*_0_ is a constant of integration. Substituting 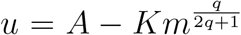, such that 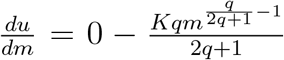 and therefore 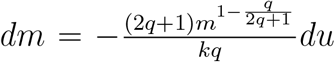,yields

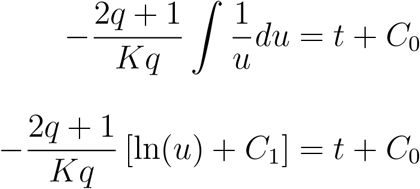

Re-substituting for *u* yields

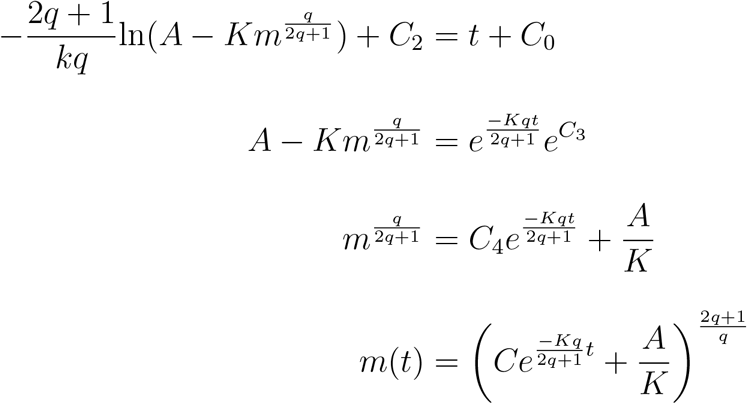

Re-substituting for *A* yields

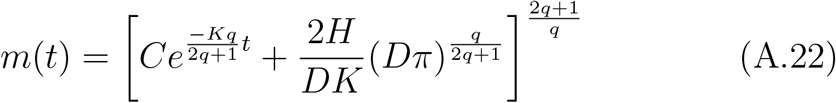

where *C*’s are modified constants of integration. This weight function has the following derivatives:

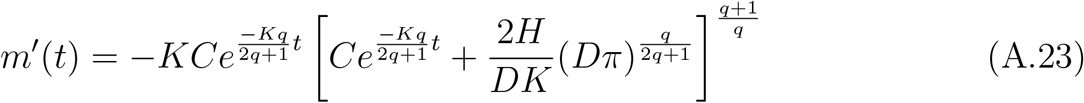

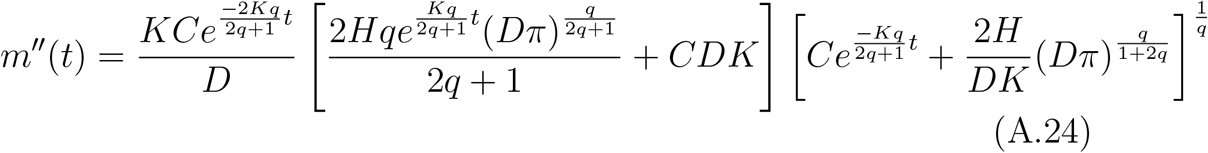

and maximum asymptotic weight

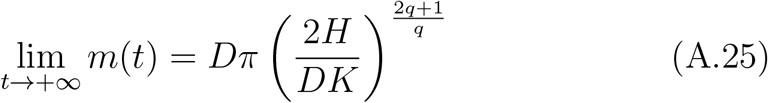

We have been unable to find a closed form solution to *m*^′′^(*t*) = 0, at which *t*_*max*_ is the age at maximum velocity and *m*^′^(*t*_*max*_) is the maximum velocity.

We define the constant *C* as follows:

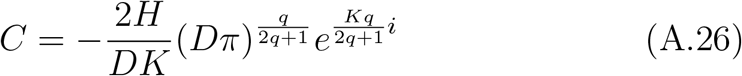

where *i* is the age at which the growth function initiates, i.e. when *m*(*t*) = 0. Substituting this value of *C* into equation A.22, and assuming, for simplicity, that density *D* = 1 g/cm^3^, yields equation 7 in the main text.

## Appendix C. Assumption of five growth processes

Change in human stature is primarily the result of: 1) mitosis and cell hypertrophy at bone articulations (epiphyses) (Hunziker 1994), which appear to occur in bursts (saltations) of several hours followed by irregular periods of stasis varying in length from days to months (Lampl et al. 1992; Lampl and Johnson 1993; Lampl 1993); 2) the effect of gravity, which compresses vertebrae during sustained upright (e.g., diurnal) posture, and is counteracted by stretching of the spinal column during horizontal (e.g., nocturnal) posture (Ashizawa and Kawabata 1990), though such nightly resilience presumably decreases over the course of senescence; and 3) the resorption of bone mass as a consequence of catabolic stress (e.g., immune system activation), especially in infancy and early childhood (Wales and Gibson 1994). Hormones, such as growth hormone (GH), insulin-like growth factors (IGFs), thyroid hormone (Robson et al. 2002), and hormones that regulate these hormones (e.g., adrenal androgens: Mühl et al. (1992), and other sex steroids: Devesa et al. (2016); Murray and Clayton (2013)), are believed to affect the amplitude and/or frequency of the mitosis- and hypertrophy-driven growth saltations (Lampl and Johnson 1993). Recent reviews (Devesa et al. 2016; Murray and Clayton 2013) suggest a consensus in the literature for the general pattern that, in all humans, fetal growth is regulated by IGFs (Gicquel and Le Bouc 2006), GH begins to take a primary role in growth regulation late in the first year after birth (Karlberg and Albertsson-Wikland 1988), and increased sex steroid production during puberty stimulates growth by increasing both GH and IGFs (Müller et al. 1999; Devesa et al. 1992; Murray and Clayton 2013). These qualitatively different phases of the hormonal regulation of growth accord well with Karlberg (1989)’s proposal that human growth comprises three superimposed (additive) components corresponding, respectively, to infancy, childhood, and puberty.

However, there are suggestions that a single growth component representing “childhood” (between approximately one year old and puberty) is insufficient. Rather, growth during this period may be regulated by several distinct, though poorly understood, (presumably) hormonal processes. In Northern hemisphere populations, the growth rates of many children exhibit seasonal patterns (often unique to the child) that repeat yearly, though the mechanisms responsible are still unclear (Marshall and Swan 1971; Marshall 1975; Togo and Togo 1982). Independent of these seasonal cycles, there is evidence that the legs and the trunk each go through three irregular cycles (with periods greater than one year) of rapid growth, followed by slow growth, between the ages of three years and puberty (Butler et al. 1990; Wales and Gibson 1994). For any particular child, these cycles of leg and trunk growth velocity may or may not coincide. When they do, one (or several) mid-childhood growth spurt(s) in overall stature may be detected (e.g., Tanner and Cameron 1980; Molinari et al. 1980; Gasser et al. 1985). When the leg and trunk growth velocity cycles are out of phase (e.g., a trough in leg growth velocity aligns with a peak in trunk growth velocity), they result in destructive interference (i.e., of waves), effectively smoothing an individual’s height velocity trajectory and masking mid-childhood growth spurt(s) in overall stature (Butler et al. 1990). It is currently unknown how these sequential cycles of childhood leg and trunk growth are regulated, though adrenal androgens may play a role in later childhood (Parker 1991; Molinari et al. 1980). Furthermore, while such sequential growth cycles have been detected in both European and Asian children (Butler et al. 1990), it is unknown whether or not they characterize the growth of all humans.

Because human growth appears to occur by saltations every few weeks (Lampl et al. 1992), it may, in theory, be appropriate to model overall growth as the sum of hundreds of overlapping processes starting at sequential ages between conception and adulthood. Overlain on these saltations are slower growth cycles of greater amplitude that occur on yearly and inter-yearly schedules, potentially regulated by the hormonal mechanisms described above. Thus, when designing a model of human growth, the choice of how many additive overlapping processes to include, if less than hundreds, becomes a practical, rather than theoretical, decision. If the model will be fit to empirical data, then the temporal resolution of the data may constrain the number of component processes. For example, if children’s height and weight are measured once each year between birth and adulthood, little insight may be gained by fitting a model in which twenty processes per year sum to give overall growth, compared to a simpler model in which only a few processes sum on an inter-yearly schedule.

The composite model in the main text incorporates five growth processes between conception and adulthood. This is more than the three processes proposed by Karlberg (1989), and adopted (with the addition of a fourth “stop” process) by Nierop et al. (2016). There are two reasons for our choice of five component processes. First, through our own exploration, we’ve found that our model with only three components (e.g., like Karlberg 1989) does not fit the data well, such that the resultant height growth trajectory qualitatively deviates in a non-trivial way from the shape of the trajectories found through fitting the JPA-1 and SITAR models (Appendices F.3 and F.4). Second, given that the Matsigenka dataset contains measurements collected no more frequently than once per year (and usually much less frequently), five component growth processes seems like a reasonable number to represent the most important of the larger-amplitude inter-yearly growth cycles suggested above. For convenience, we label these growth processes by the colloquial life stages in which each process tends to have the greatest impact on overall growth: 1. in utero; 2. infancy; 3. early childhood; 4. late childhood; and 5. adolescence (Figure 1). Note that the ages at which these processes initiate (*i* parameters, Figure A.8) do not necessarily fall within these colloquial life phases. Furthermore, given the current absence of independent knowledge regarding the onset of these growth processes, priors on the *i* parameters (equations A.41 and A.59) are manually chosen simply to facilitate model convergence.

Given that equations 6 and 7 represent the body as a cylinder (Figure A.1), equations 8 and 9 represent the body as a set of five cylinders. As explained in Appendices D.1.1 and D.1.2, each cylinder is envisioned as comprising all of the living skeletal cells that contribute to growth under the corresponding growth process (1 through 5). Importantly, the living skeletal cells are distributed throughout the body, and are not concentrated in a single cylinder. The cylindrical representation captures the intuition that growing skeletal cells increase both the height and the radius of the skeleton (and, by extension, the surface area of the intestine), such that the relationship between height and radius is *r* = *h*^*q*^ (Appendix A). One interpretation of the five distinct growth processes is that different skeletal cells are the targets of different regimes of growth-promoting hormones produced at the initiation of each process. These groups of cells grow in such a way that they may have different effects on the relationship between overall skeletal height and radius, which is expressed in different values of *q* for each growth process. Thus, the cylindrical representations of the skeletal cells active in the five growth processes should not be envisioned as stacked one on top of the other. Rather, each cylinder is perhaps better envisioned as embedded inside the growing cylinder from the previous growth process.

## Appendix D. Empirical estimation of the growth model

### D.1. Biological assumptions for model fitting

#### D.1.1. Relating intestine to living skeleton

Equations 6 and 7 represent growth of only those parts of the body whose increase in size increases the surface area through which molecules are absorbed. Therefore, in humans, these equations apply only to the intestine and the parts of the body that determine and constrain its surface area, which is a function of its length, diameter, and density of villi and microvilli. The intestine fills a large part of the lower abdominal cavity, which increases in size as the skeleton increases in size. We thus make the simplifying assumption that intestinal surface area is determined by the skeleton, whose size and weight can be (approximately) calculated from height and a proportion of total body weight.

The skeleton is composed of bone marrow cells, as well as other living cells embedded in metabolically-inactive bone matrix (composed primarily of collagen and calcium phosphate) and cartilaginous matrix (composed primarily of collagen) (International Commission on Radiological Protection 1995). Because the theory described here represents growth as a balance between anabolism and catabolism, equations 6 and 7 are applicable only to the metabolically-active cells of the skeleton. Table A.1 presents the different types of living cells in the skeleton as approximate proportions of total body weight at different ages. Note that these proportions are calculated by applying many generalizations and simplifying assumptions to extremely limited data, and thus serve as very rough approximations.

**Table A.1.** Proportion of total female body mass contributed by metabolically active cells in different parts of the skeletal system during development.

| Age<sup>a</sup> | Bone<sup>b</sup> | Marrow<sup>c</sup> | Cartilage<sup>d</sup> | Total |
| --- | --- | --- | --- | --- |
| Birth | 0.009 | 0.013 | 0.0037 | 0.026 |
| 1 | 0 | 0.017 | 0.0038 | 0.021 |
| 5 | 0 | 0.025 | 0.0033 | 0.028 |
| 10 | 0 | 0.038 | 0.0025 | 0.041 |
| 15 | 0 | 0.045 | 0.00017 | 0.045 |
| Adult | 0 | 0.045 | 0.00014 | 0.045 |

<sup>a</sup>Age in years since birth
<sup>b</sup>From International Commission on Radiological Protection \(1995\), skeletal weights (pg 14) - marrow weights (pg 67) = weights of bone matrix and osteocyte cells, which are then divided by body weights for the respective ages: 3.5, 9.5, 18, 33, 55, 65 kg, yielding the proportion of total body weight comprising bone matrix and osteocytes. Assume, at birth, 10% of human bone mass (assuming uniform density of 1 cm<sup>3</sup>/g) comprises osteocyte cells, like in embryonic chickens (Sugawara et al. 2005), which decreases by 82% (yielding a negligible proportion) at one year old, like in adult pigs (Mullender et al. 1996) (pig: 43000 osteocytes per mm<sup>3</sup>, chicken: 1 osteocyte / 0.00000418 mm<sup>3</sup> = 239234 osteocytes/mm<sup>3</sup>).
<sup>c</sup>Taken from Table 39 of International Commission on Radiological Protection \(1995\)
<sup>d</sup>Cartilage weights taken from pg 58 of International Commission on Radiological Protection \(1995\), multiplied by 0.1 cm<sup>3</sup> chondrocyte cell volume per 1 cm<sup>3</sup> cartilage (Bačenková et al. 2023) (assuming uniform density of 1 cm<sup>3</sup>/g), divided by the body weights above

Loosely guided by these data, we designed the function in equation A.27 to represent how the proportion *p* of total body mass comprising living structural cells (e.g., of the skeleton) changes with age *a*, in years since conception.

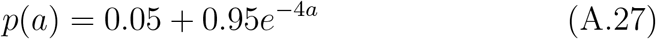

This function is plotted in Figure A.5. At conception, the proportion is one, as, after formation of the blastocyst, we assume that any new cell added in early embryonic development increases both (pre-)embryo mass, as well as (external) surface area. In other words, early in development, we assume the body comprises a cluster of cells with an external absorbing surface, i.e., all cells are structural. An important assumption of this model is that the proportion *p* decreases rapidly as the structural tissues that will become the skeleton differentiate and develop in the fetus, reaching a constant proportion of 0.05 at approximately one year of age (since birth) when the skeleton is sufficiently rigid to support upright posture and walking, and, we assume, has become the primary constraint on intestinal growth. The validity of this assumption requires future study.

#### D.1.2. Intestinal surface area and q

To approximate intestinal surface area, we use a cylindrical approximation of the surface area (derived from mass and height through equations 6 and 7) of the living cells of the skeleton, which we assume is equivalent to the corresponding proportion (i.e., from equation A.27) of skin surface area. To convert skin surface area to intestinal surface area, we use the following approximation: a skin surface area of 20,200 cm^2^ (Mosteller 1987) corresponds to an intestinal surface area of 320,000 cm^2^ (Helander and Fändriks 2014). Thus, when fit to data on height of the body and weight of living skeletal cells, the units of the parameter *H* are g of constructed molecules per cm^2^ surface of a cylindrical approximation of living skeletal cell mass. To convert to g per cm^2^ of intestinal surface, the value *H* is multiplied by 20, 200*/*320, 000 = 2*/*32. Note that this conversion from skin surface area to the surface area through which molecules are absorbed is only appropriate when that absorbing surface area is the intestine, i.e., after birth. In utero, the absorbing surface area is the interface between the placenta and the uterus, which may have a different relationship to skin surface area. Thus, the objective value of *H* should be interpreted with caution for ages prior to birth.

It is difficult to know what proportion of total body height is contributed by the living cells of the skeleton (without the bone matrix).

**Figure A.5.**
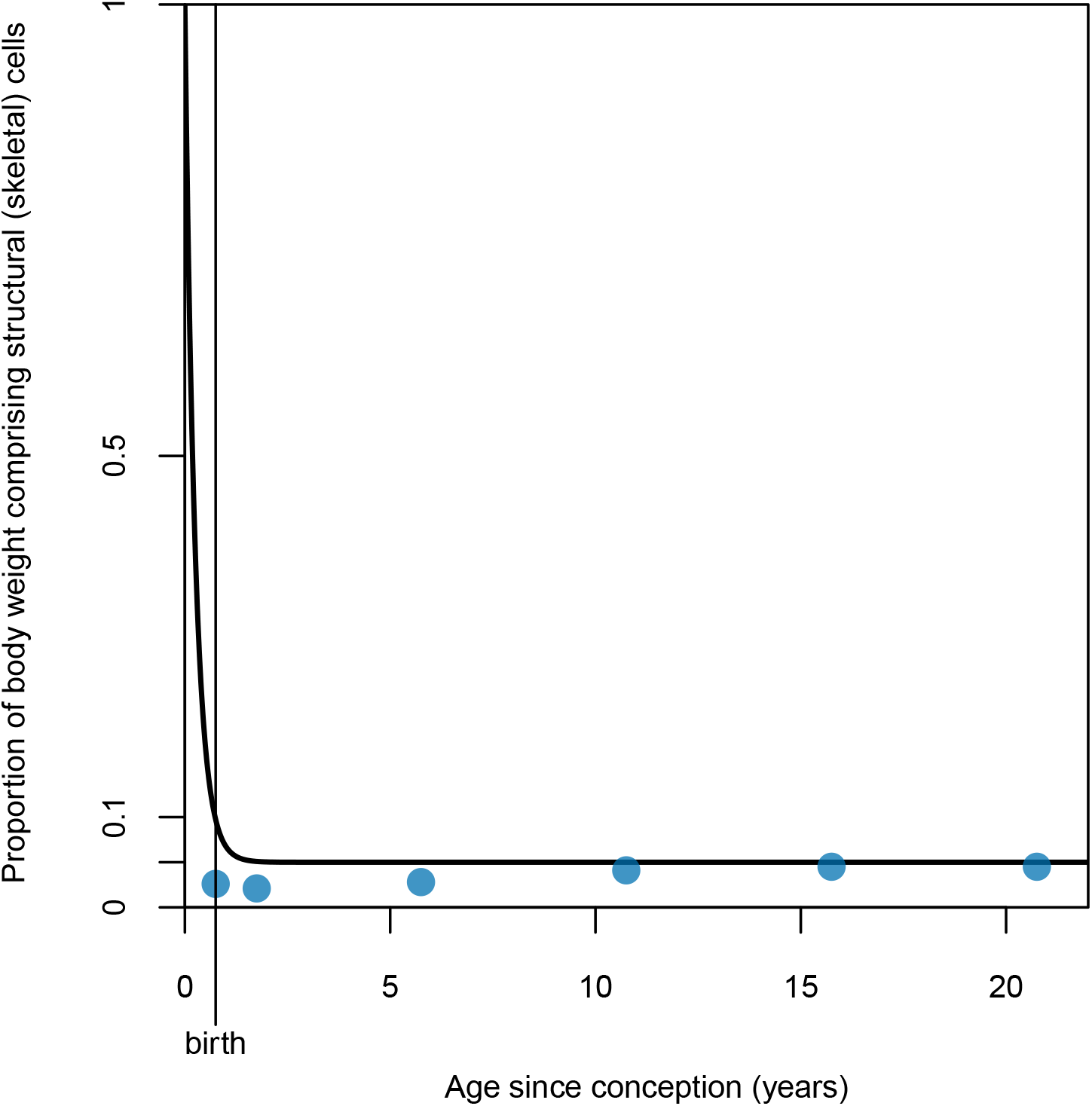
Proportion of total body weight comprising metabolically active structural (skeletal) cells by age, according to equation A.27. Blue points are approximate empirical proportions from Table A.1

To the extent that this proportion is less than one, the parameter *q* will be under-estimated. This is because, in order to match the surface area of the cylindrical approximation of skeletal cells derived from the weight of these cells, given an over-estimate of the length of the cylinder (i.e., total skeletal height), the radius of the cylinder must be underestimated in order to compensate. In other words, given surface area 2*πrh* where *r* = *h*^*q*^ for 0 *< q <* 1, an increase in *h* will yield an equivalent surface area only if *r* is sufficiently reduced, which can be accomplished only through an appropriate reduction in *q*. Thus, the objective values of *q* estimated using total body height must be interpreted with caution, as they are almost certainly underestimates. However, relative comparisons of *q* derived from different growth trajectories are meaningful, such that a larger *q* indicates (given the assumptions above) a skeleton that is wider for a given height (or, complementarily, shorter for a given weight), with a correspondingly greater intestinal surface area.

#### D.1.3. Fitting height and weight together

Equations 6 and 7 constitute a theoretical model for how metabolism (parameters *K* and *H*), allometry (parameter *q*), and the timing of growth processes (parameter *i*) affect growth in both height and weight simultaneously. Therefore, according to this theory, fitting equation 6 to height data and, independently, fitting equation 7 to weight data from the same individuals should yield identical estimates for the four parameters.

In practice, we have found that different combinations of values for *K, H, q*, and *i* can be chosen to produce very similar overall growth trajectories. When the model is fit to just one type of data, i.e., fitting only equation 6 or only equation 7, it can be difficult to distinguish among these trajectories, and therefore to know which set of parameter values best characterizes measured growth. However, we have found that parameter values that produce similar height trajectories often produce very different weight trajectories, and vice versa. This motivates us to fit equations 6 and 7 simultaneously to the combined record of individuals’ height and weight measurements. In this way, we have more confidence that parameter estimates better reflect processes affecting both aspects of body growth.

A complication to simultaneously fitting equations 6 and 7 is the possibility that the proportion of total body weight comprising the skeleton (specifically, living skeletal cells) is not constant during ontogeny. For instance, it is plausible that, for many individuals, an increase in muscle and fat mass at certain points during puberty may be faster than the corresponding increase in skeletal mass, thereby decreasing the proportion of total body weight represented by the skeleton. In contrast, the proportion represented by equation A.27, above, is effectively constant, at 0.05, after approximately one year of age since birth (Figure A.1). The same problem is not encountered with height data, as skeletal height is relatively unaffected by an increase or decrease in fat and muscle.

Thus, in contrast to height measurements, we assume that some variation in weight measurements does not reflect skeletal (and, by extension, intestinal) growth. Our strategy to account for this is to allow the statistical model a higher tolerance for “measurement error” when fitting equation 7 to weight, than when fitting equation 6 to height. In effect, this allows the model to fit weight trajectories relatively poorly compared to height trajectories, while still allowing weight data to inform parameter estimates (though to a lesser extent than height data). See Appendices D.2 and D.3 for a description of how this is implemented in the statistical model. See Figures 2 and A.9 to note the difference in fit of estimated trajectories to height and weight data. See Appendix F.1 for the same model without the bias for greater weight measurement error when fit to both height and weight data. See Appendix F.2 for the model fit only to height data.

### D.2. Estimating the baseline trajectory

To fit the composite model (equations 8 and 9) to U.S. and Matsigenka height and weight measurements (raw data plotted in Figure A.6), our strategy is to first derive baseline human height and weight trajectories from which individual U.S. and Matsigenka children’s height and weight trajectories can diverge. Estimating a baseline human trajectory around which to center informative priors increases the efficiency of Stan’s Hamiltonian Monte Carlo fitting algorithm, which can then limit its search of the posterior parameter space to regions plausibly within the range of human variation in growth.

**Figure A.6.**
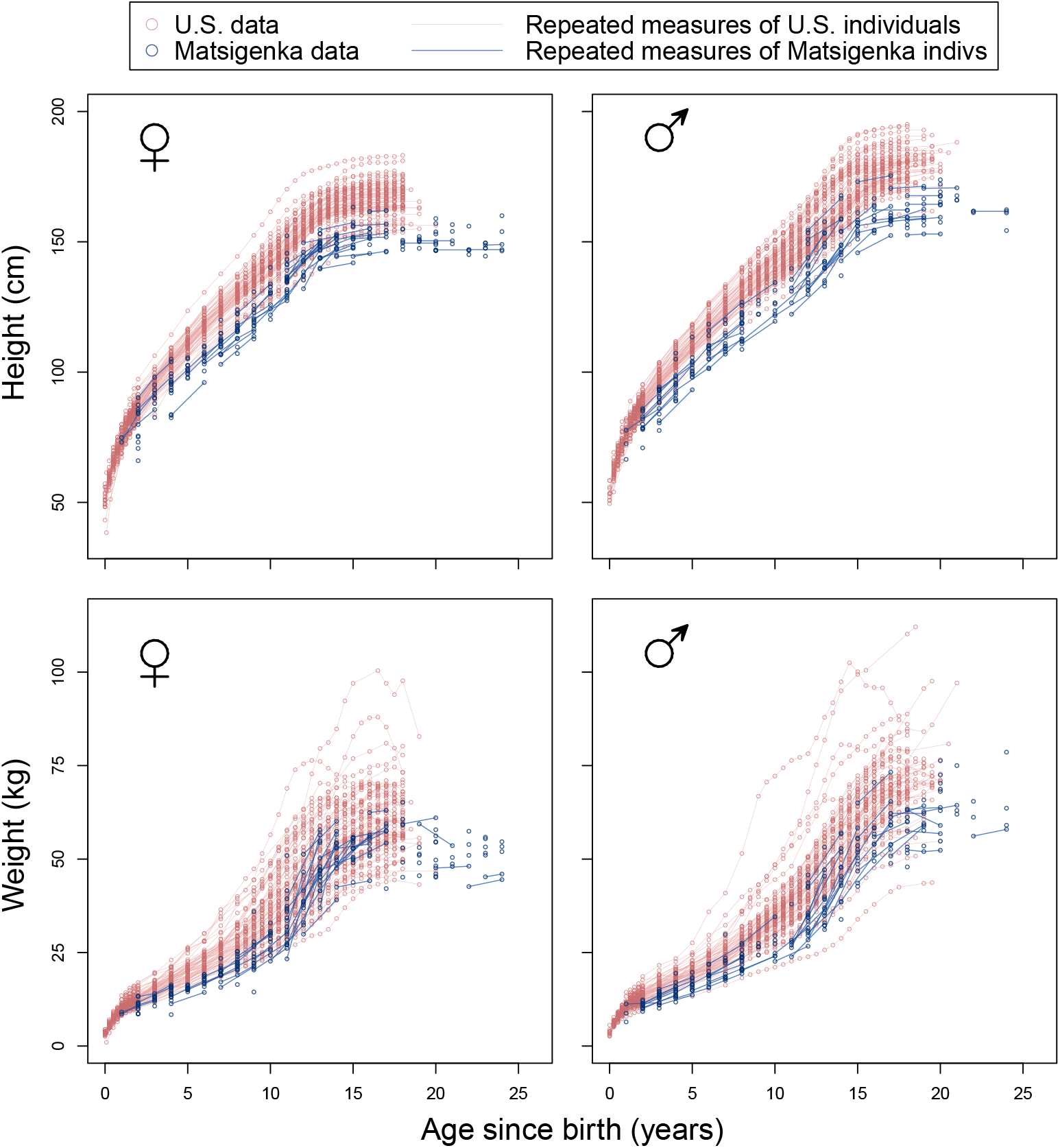
Height and weight measurements by ethnic group. Shown are the height (upper row) and weight (lower row) measurements of U.S. (red) and Matigenka (blue) children used in this study. Measurements from the same individual at different ages are connected by lines. The temporally-dense U.S. data are taken from Tuddenham and Snyder (1954), while the temporally-sparse Matsigenka data were collected by the authors.

To derive the baseline trajectory, we fit the composite model simultaneously to U.S. children’s height and weight measurements (treated as cross-sectional data), thereby approximating the mean height and weight trajectories of these children. We use the U.S. dataset for this purpose only because there is so much data, not because U.S. children are necessarily representative humans. Rich datasets from other populations would serve just as well for the derivation of a baseline growth trajectory. Importantly, this analysis strategy makes no assumption that the baseline trajectory represents “healthy” or “normal” growth. Furthermore, we neither present nor interpret any comparison of individual or population-mean trajectories with this baseline trajectory.

We fit the composite height and weight model to the U.S. dataset, performing separate analyses for females and males. To represent the fact that observed heights and weights can never be negative, we take their log. The means (ln (*η*_*t*_) and ln (*µ*_*t*_)) of the Normal likelihoods are the logs of the composite height and weight functions for age *t*, and the scales of these likelihood distributions represent measurement error (see equations 10 and 11). Observed height (*h*_*t*_) and weight (*m*_*t*_) at time *t* are modeled as:

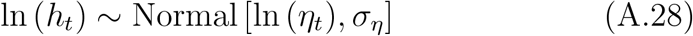

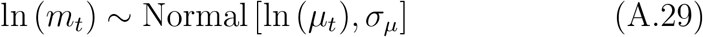

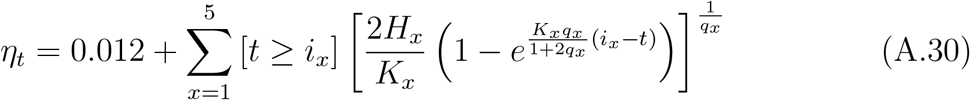

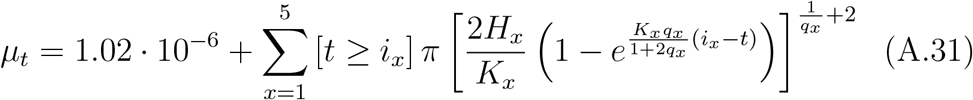

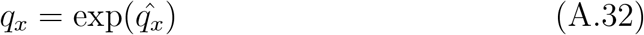

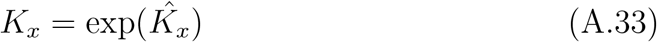

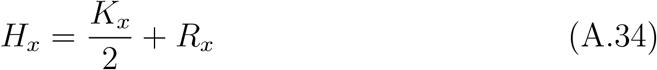

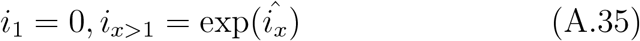

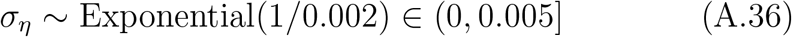

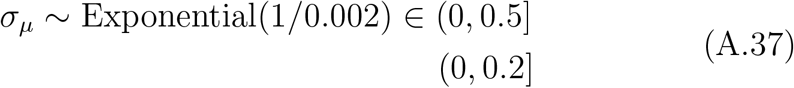

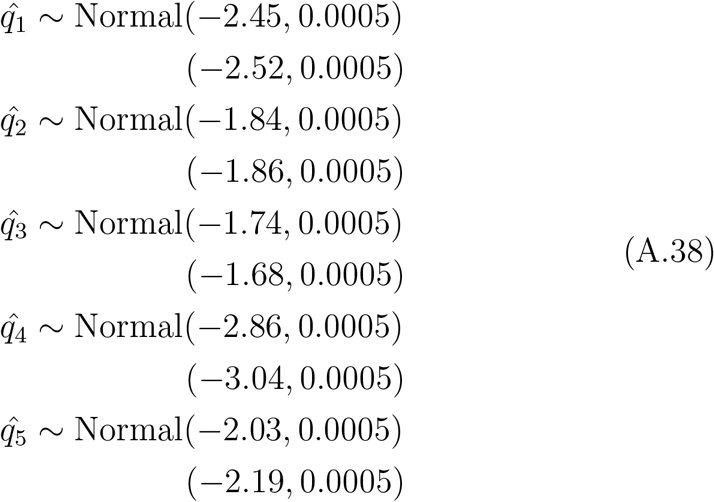

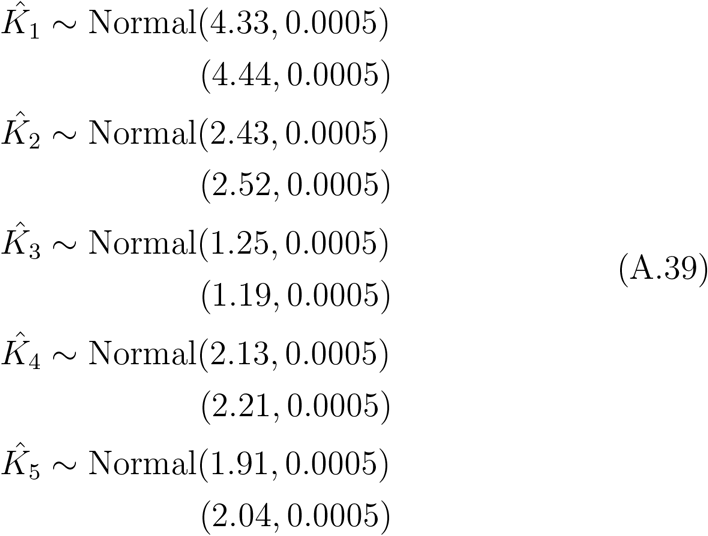

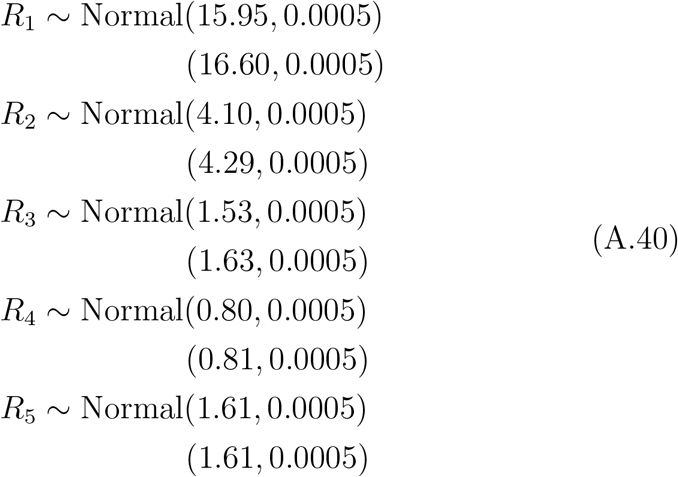

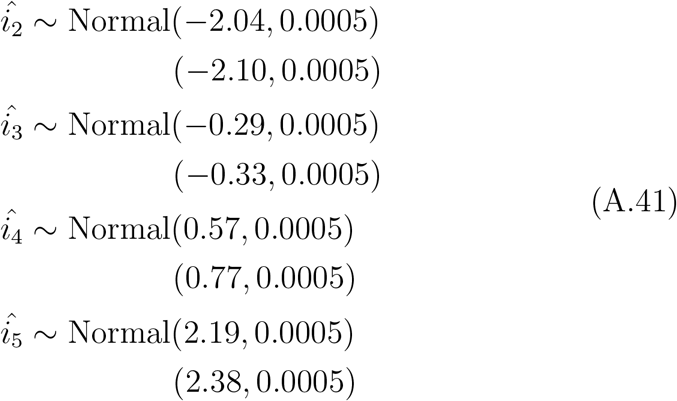

In equations A.37–A.41, values for priors used to fit males are shown beneath those used to fit females. The first terms in equations A.30 and A.31 represent a fertilized egg at conception with diameter (i.e., height) 0.012 cm (Leary et al. 2014) and weight 1.02*·*10^−6^ g (Hatton et al. 2023, https://humancelltreemap.mis.mpg.de/, for ovum mass). To increase the efficiency of the fitting algorithm, parameters in the composite model are transformed (e.g., exponentiated: equations A.32, A.33, and A.35 or decomposed into sums: Equation A.34). Priors on the scales (equations A.36 and A.37) represent measurement error, where the parameter of the exponential distribution is 1/mean. These standard deviations on the log scale are expressed on the scale of the data as, for example, 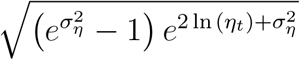, where *σ*_*µ*_ is transformed analogously. Prior means of 0.002 correspond, on the scale of the data, to measurement errors such that approximately 95% of observations fall within 2*stdev = 0.4 cm of the actual height at 100 cm, and within 200 g of the actual weight at 50 kg. As explained in Appendix D.1.3, fitting the model to height and weight data simultaneously requires allowing measurement error for weight (*σ*_*µ*_) to be greater than that for height (*σ*_*η*_). Thus, the bounds in equation A.37 are wider than in equation A.36.

Because there is limited pre-existing knowledge about the values of the *q, K, H*, and *i* parameters in this model, initial prior means for these parameters were chosen by hand to make the height and weight trajectories look as similar as possible to human height and skeletal cell weight (Appendix D.1.1) trajectories. The complexity of the parameter space makes fitting the model very difficult without small standard deviations on the Normal prior distributions for these parameters (equations A.38–A.41). This strategy introduces the risk that the model would fit the data better using combinations of parameter values that fall outside of the highly constrained parameter space explored in this analysis. At present, we have been unable to find a satisfactory alternative approach to fitting this otherwise unidentified model. However, using this baseline trajectory in a multilevel version of the model (Appendix D.3) yields posterior mean parameter values that correspond, in a general sense, to reasonable human metabolic rates (Appendix D.4). Models were fit in R (R Core Team 2022) and Stan (Stan Development Team 2022) using the cmdstanr package (Gabry and Cesnovar 2021). Because this is a preliminary model simply to provide reasonable priors for the means of the *q, K, H*, and *i* baseline parameters, we are less concerned with running the model to convergence as indicated by 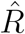 values of 1.00 (McElreath 2020). Thus, we fit the model using only two chains of 1000 samples each, half of which were warm-up.

Data and analysis scripts are available at https://github.com/jabunce/bunce-fernandez-revilla-2022-growth-model.

### D.3. The multilevel model

We fit the following model of the height and weight of person *j* at time *t* to the combined U.S. and Matsigenka datasets (females and males fit separately), with ethnic group-level (*g*) and person-level (*p*) offsets to the previously-estimated mean parameter values (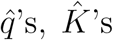, and *R*’s) for the baseline human height and weight trajectories (above). We allow both sets of offsets to covary within and across growth phases. Person *j*’s observed height (*h*_*jt*_) and weight (*m*_*jt*_) at time *t* are modeled as:

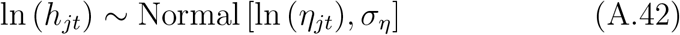

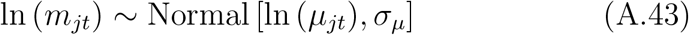

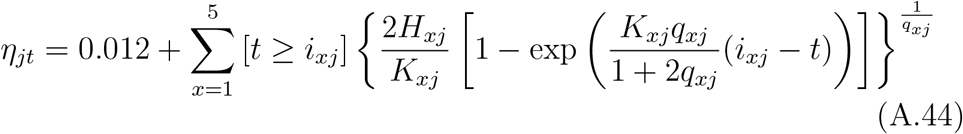

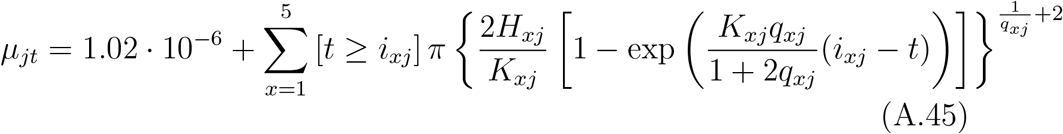

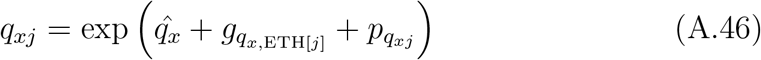

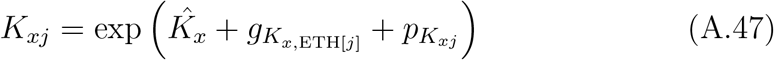

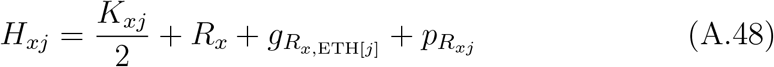

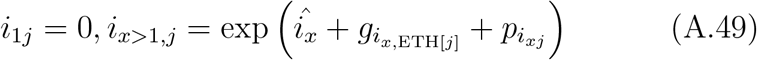

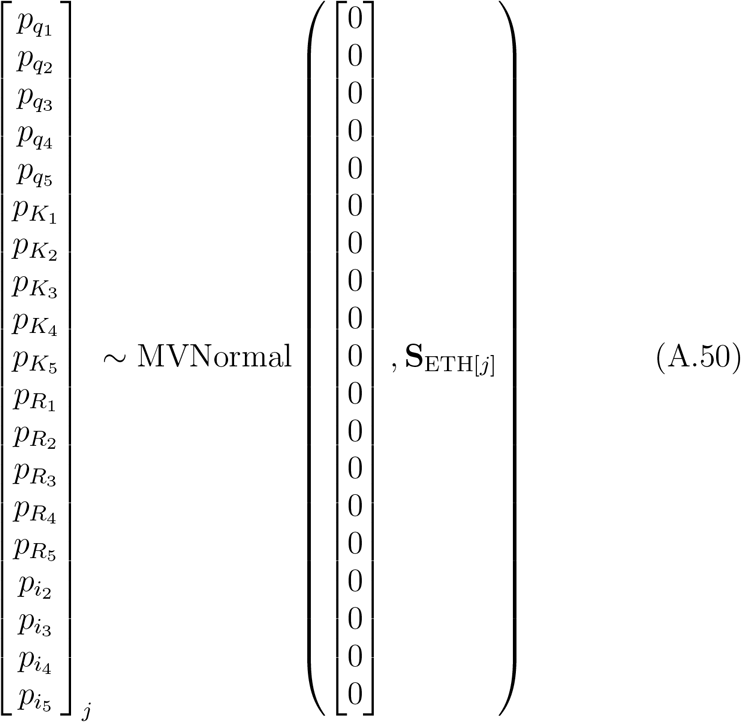

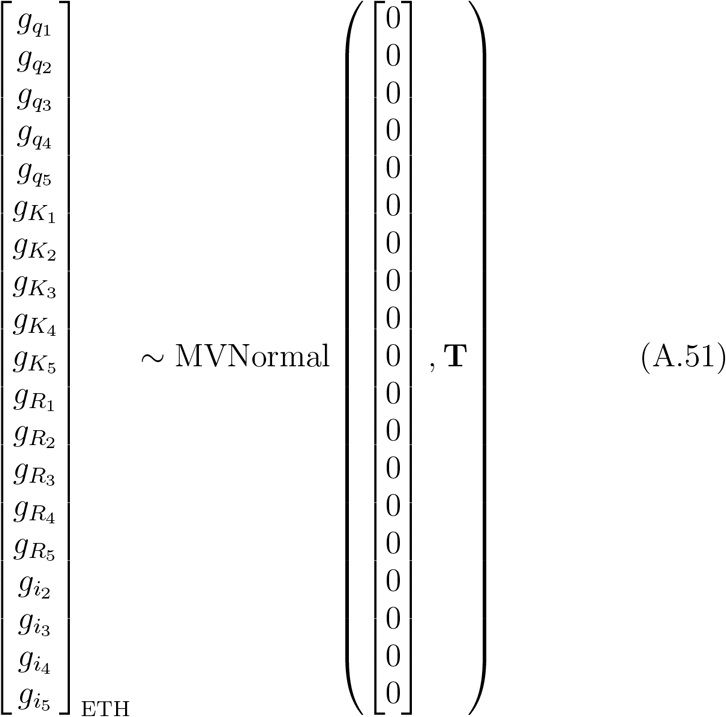

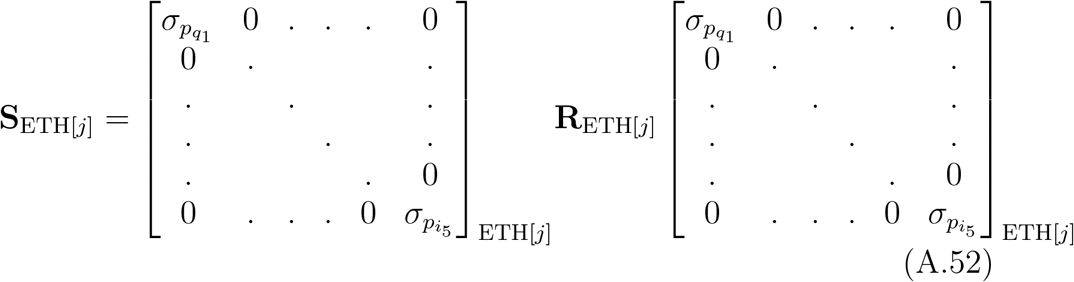

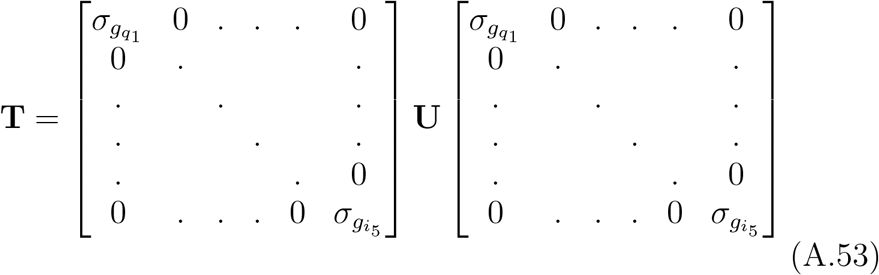

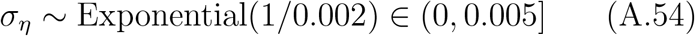

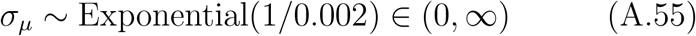

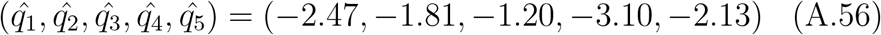

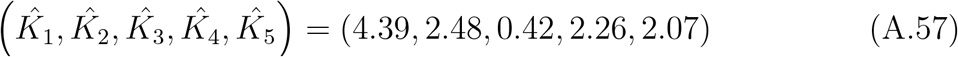

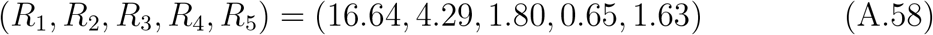

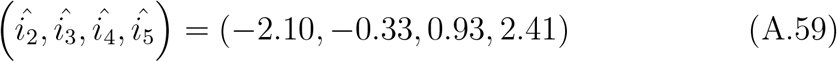

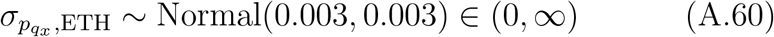

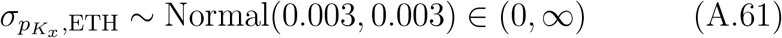

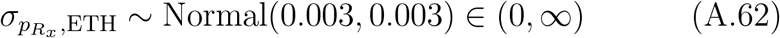

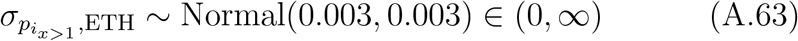

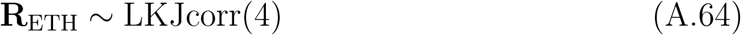

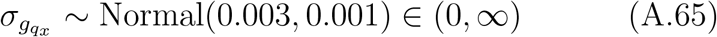

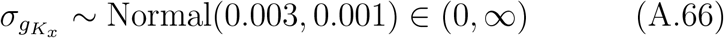

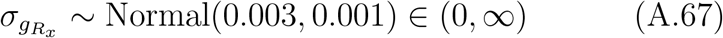

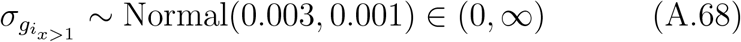

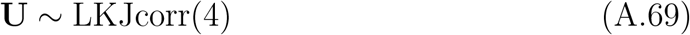

where the subscript ETH[*j*] refers to the ethnicity (U.S. or Matsigenka) of individual *j*. In practice, the covariance matrices **S** and **T** are fit using Cholesky decomposition (McElreath 2020). As explained in Appendix D.1.3, fitting the model to height and weight data simultaneously requires allowing measurement error for weight (*σ*_*µ*_) to be greater than that for height (*σ*_*η*_). Thus, the bounds in equation A.55 are wider than in equation A.54. An upper bound of 0.005 on *σ*_*η*_ corresponds, on the scale of the data, to measurement error such that approximately 95% of measurements fall within 2*stdev = 1 cm of the actual height when height = 100 cm. Priors on these measurement error terms are explained in Appendix D.2. Priors on the transformed parameters, 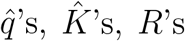, and 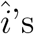, are derived from the posterior means for these parameters after fitting the baseline model (Appendix D.2) to data from males. The same priors were used to fit this multilevel model separately to males and females.

The ethnic group-specific individual-level standard deviations (*σ*_*p*_’s), as well as the the group-level standard deviations (*σ*_*g*_’s), are sampled from Gaussian distributions truncated at zero. The means of these prior distributions are chosen simply to yield adequate model convergence and fit, as we currently have limited prior information about how much variation exists between different human populations with respect to *q, K, H*, and *i*. Priors on the correlation matrices **R** and **U** are chosen to bias against extreme correlations (McElreath 2020).

Models were fit in R (R Core Team 2022) and Stan (Stan Development Team 2022) using the cmdstanr package (Gabry and Cesnovar 2021). Convergence was indicated by 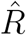 values of 1.00. This usually required four chains of 4000 to 5000 samples each, half of which were warm-up. Data and analysis scripts are available at https://github.com/jabunce/bunce-fernandez-revilla-2022-growth-model.

Posterior parameter estimates for each of the five growth processes are shown in Figures A.7 and A.8. Posterior mean standard deviations for height (*σ*_*η*_) for females and males are both approximately 0.005 (i.e., the upper bound in equation A.54). Posterior mean standard deviations for weight (*σ*_*µ*_) for females and males are 0.15 and 0.18, respectively. These correspond, on the scale of the data, to measurement errors for weight such that approximately 95% of measurements fall within 15.2 kg and 18.6 kg, respectively, of the actual weight, when weight equals 50 kg.

**Figure A.7.**
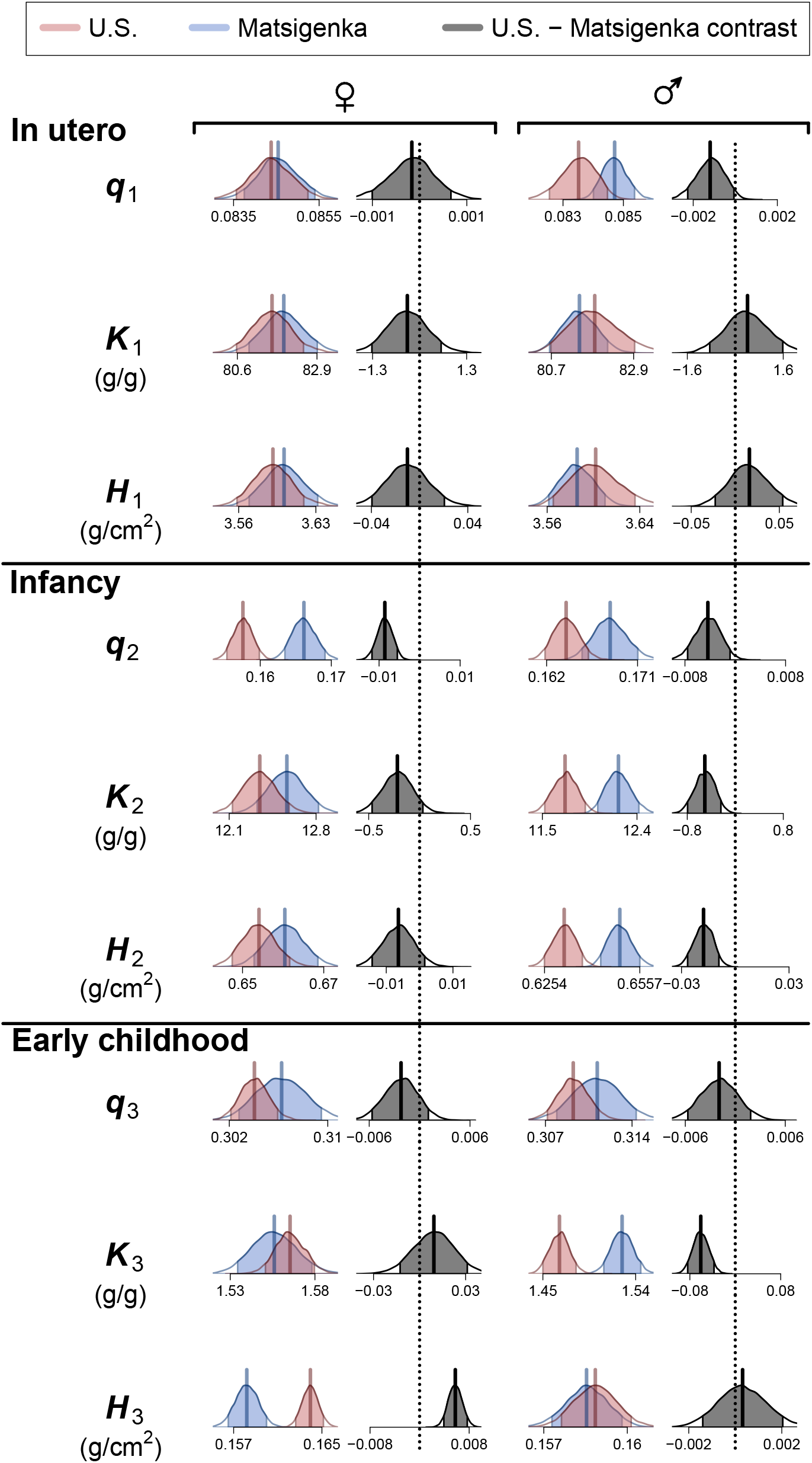
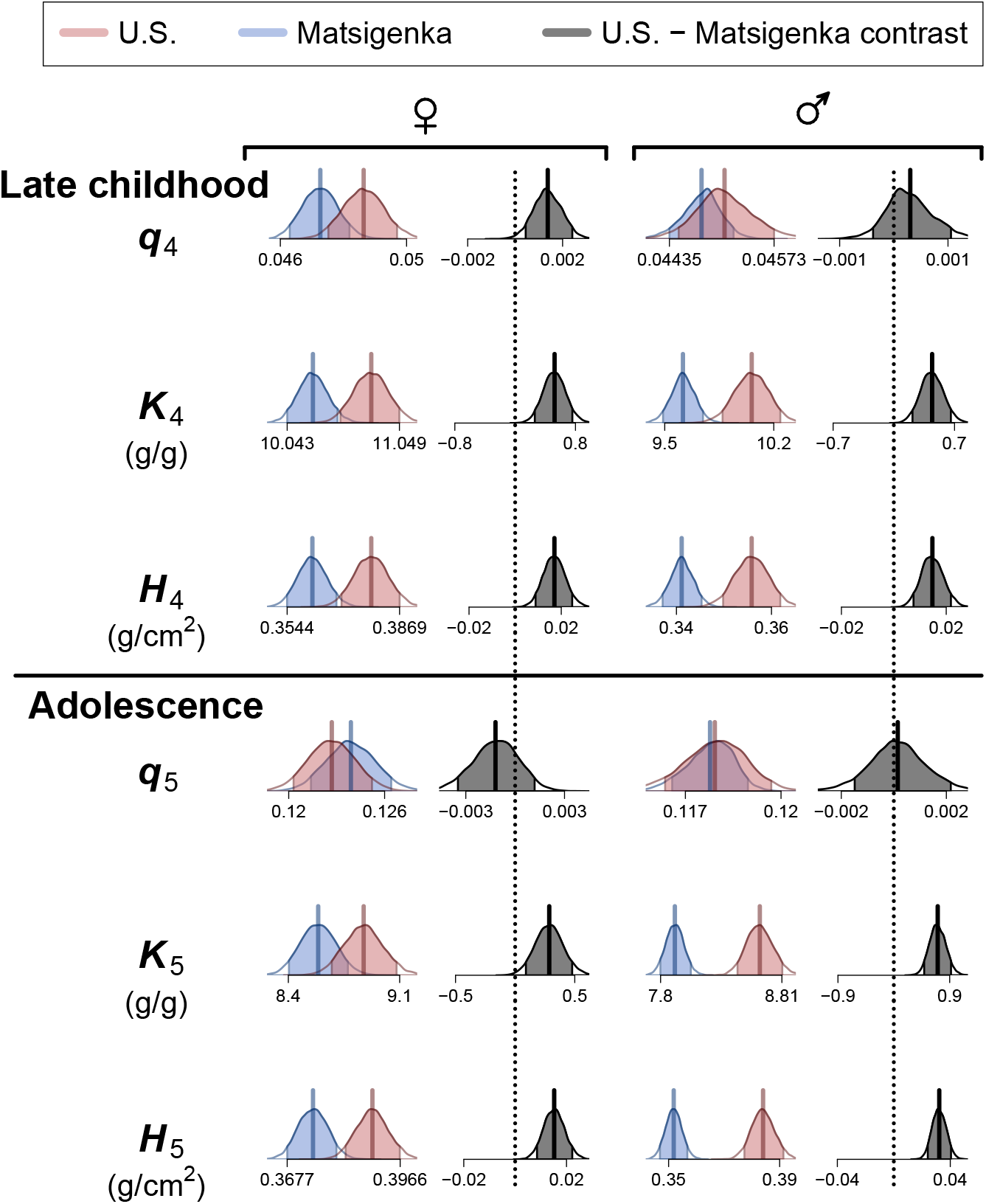
Estimates of mean parameter values by ethnic group. Shown are 90% highest posterior density intervals (HPDI) for posterior distributions of model parameters used to construct the mean height and weight trajectories (Figures 2 and A.9) of U.S. (red) and Matsigenka (blue) children, by growth process. Posterior density outside of this HPDI is shown as white tails on a distribution. Distribution means are shown as solid vertical lines. The 90% HPDI of the U.S. - Matsigenka contrast (difference) is shown in grey. HPDIs of contrasts that do not overlap zero (dotted vertical lines) indicate a detectable ethnic-group difference in parameter estimates. Interpretation of parameters and their estimated values is only meaningful under the strict assumptions used to derive the theoretical growth models, explained in the text.

**Figure A.8.**
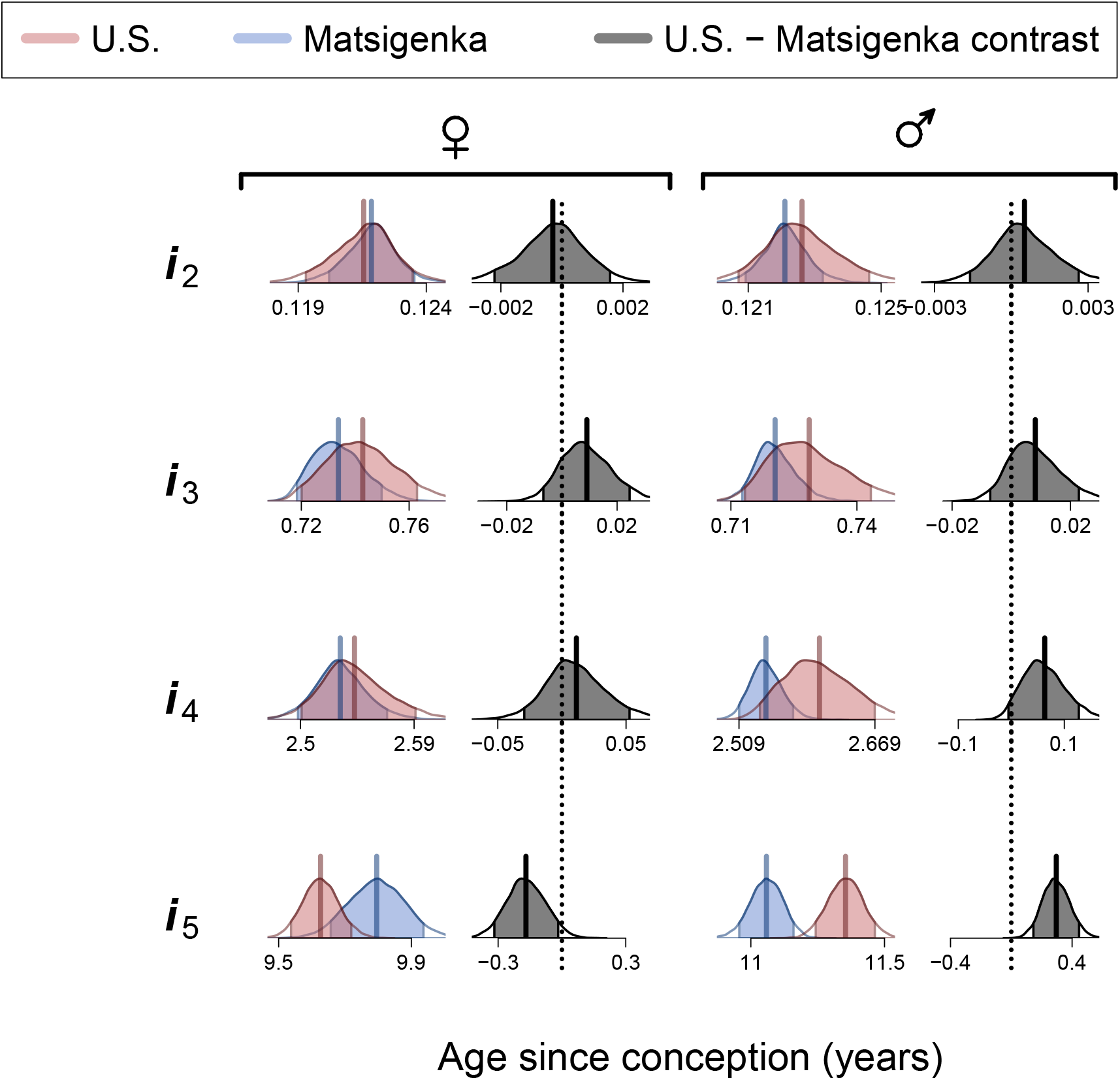
Estimates of mean *i* parameter values by ethnic group. The parameter *i*_1_ is fixed at zero, and represents the start of the growth process that begins at conception. Plot features are analogous to Figure A.7.

Growth trajectories calculated from posterior parameter estimates are shown in Figure 2 (height trajectories) and Figure A.9 (weight trajectories). Figure A.10 incorporates uncertainty in the estimated U.S. and Matsigenka mean growth trajectories in order to compute population-level contrasts of three important descriptive characteristics of these trajectories: maximum achieved height, maximum achieved growth velocity during puberty, and the age at maximum velocity during puberty. Consistent with Figure 2, U.S. and Matsigenka children are, on average, often predicted to be reliably distinguishable on the basis of these growth characteristics. In particular, Matsigenka boys and girls have lower estimated mean maximum height and lower mean maximum velocity during puberty. The model suggests that Matsigenka girls reach maximum pubertal growth velocity at later ages than U.S. girls, while the corresponding ages for U.S. and Matsigenka boys are not reliably distinguishable. An analysis of residuals to asses model fit is presented in Appendix G.

**Figure A.9.**
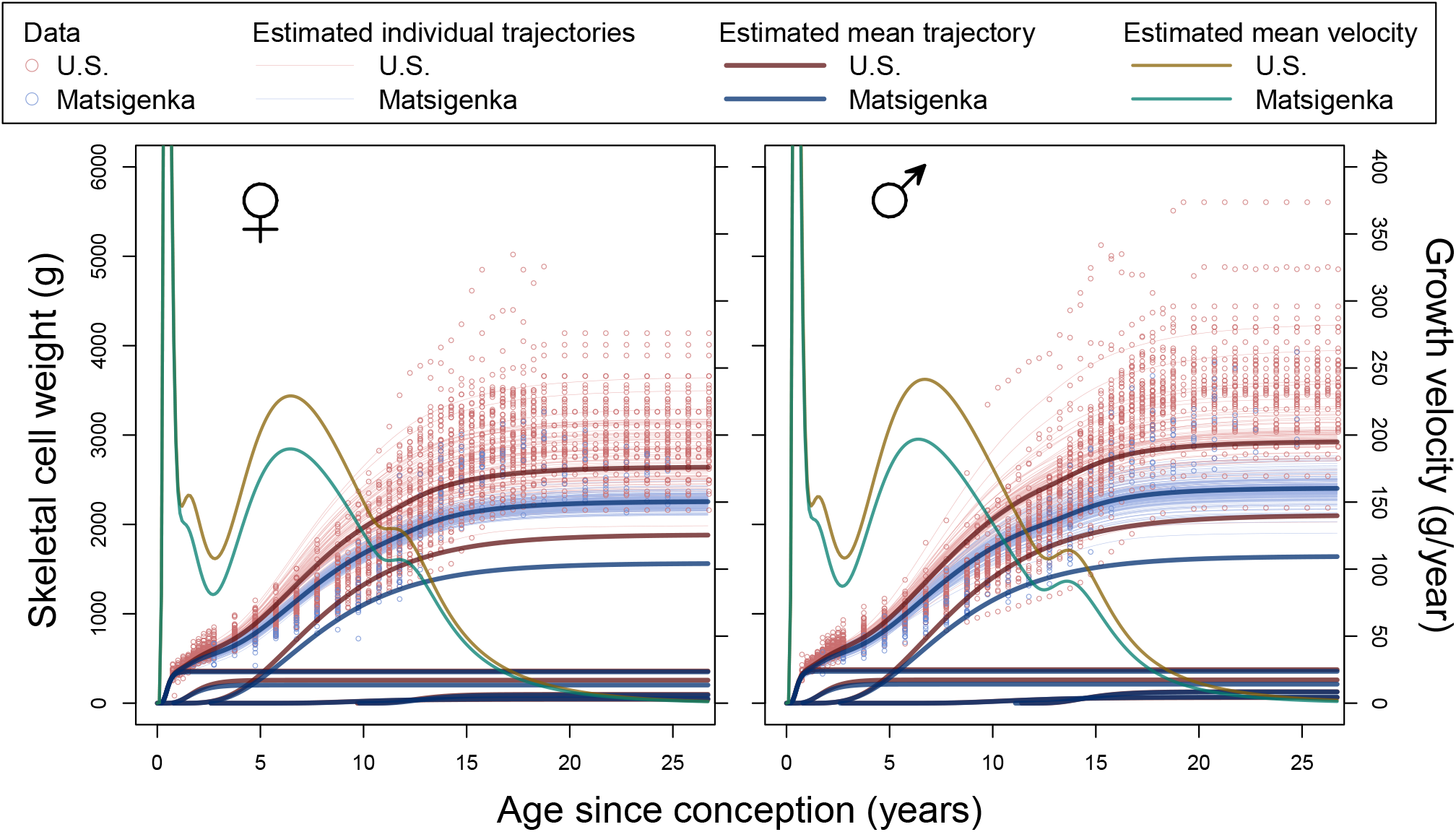
Estimated weight trajectories. This is analogous to Figure 2, using posterior parameter estimates from the multi-level model in equations A.42–A.69 to construct weight trajectories.

**Figure A.10.**
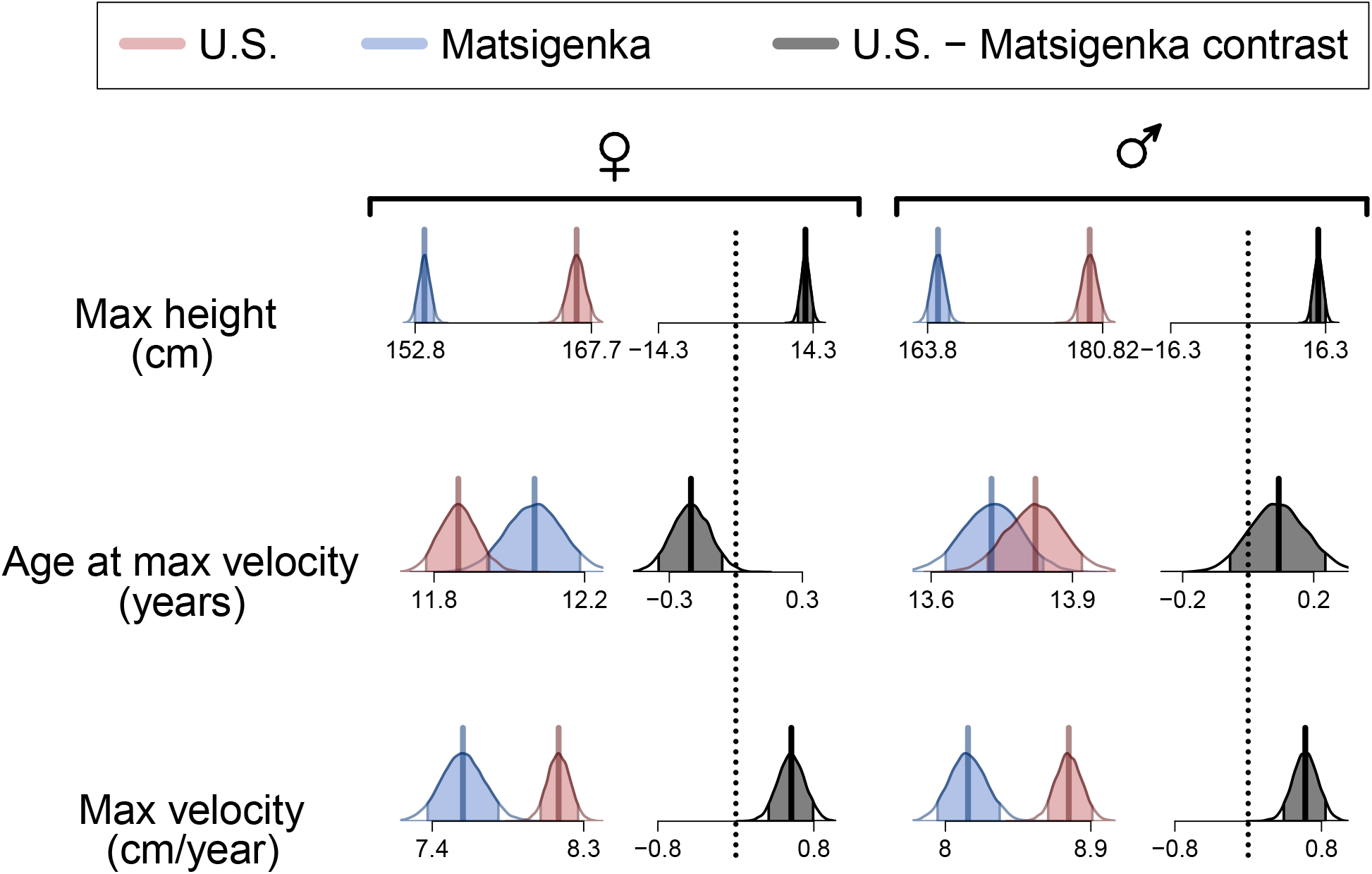
90% highest posterior density intervals (HPDI: McElreath (2020)) for posterior distributions of characteristics of the mean height trajectories of U.S. (red) and Matsigenka (blue) children. Plot features are analogous to Figure A.7. Maximum velocity and age at maximum velocity are calculated during puberty, rather than in utero when velocity is much greater (Figure 2).

### D.4. Metabolic and allometric trajectories

The overall (i.e., body-wide) value of an allometric or metabolic parameter (*q, K*, or *H*) at age *t* is the sum of the relevant parameters of each component process that has started at or before age *t*, weighted by the proportion of overall height at age *t* contributed by that process. We assume this to be proportional to the number of skeletal cells contributed to overall height by each process at age *t*.

For metabolic parameters *K* and *H*, this weighted sum at age *t* is further weighted by the proportion of metabolic activity being realized by cells in each process. The assumption is that, when a process ends (i.e., approaches its asymptote), “active” red marrow cells have been converted into marrow fat cells (International Commission on Radiological Protection 1995) which, though metabolically active (Pham et al. 2020), metabolize at 25% the rate of red marrow cells. We emphasize that this percentage is an assumption, given the current absence of data (to our knowledge), chosen to qualitatively reflect known patterns of marrow cell turnover, and results in reasonably realistic quantitative estimates for *H* and *K*.

Given

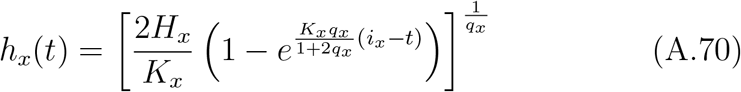

is the derived equation for height at age *t* contributed by component process *x* = 1…5, then equation 8 represents total height at age *t, h*_*tot*_(*t*). The asymptotic maximum height of component process *x, h*_*x,max*_ (see equation A.11) is

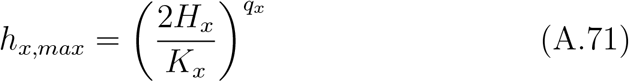

The overall values of the three parameters at age *t* are then:

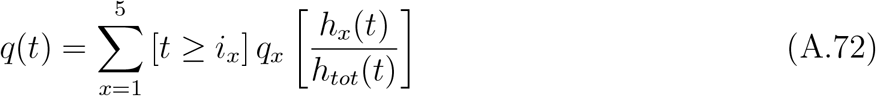

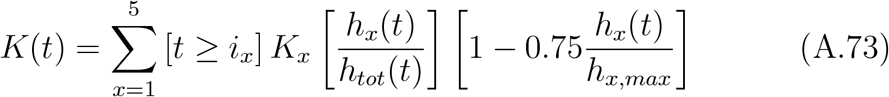

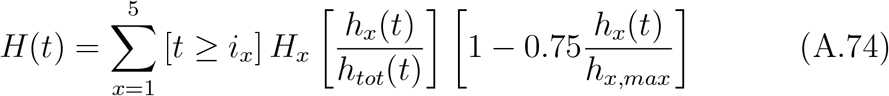

The three functions represented in equations A.72–A.74 are plotted for Matsigenka and U.S. girls and boys in Figure 3. Tables A.2–A.4 present example calculations of the mean posterior values of overall *q, K*, and *H* (respectively) for boys at age 15 years since conception, as plotted in Figure 3.

**Table A.2.** Calculations for mean overall *q* for males at 15 years since conception.

| Group | Parameter | Value<sup>a</sup> | Prop Total Height<sup>b</sup> | Product<sup>c</sup> | Sum<sup>d</sup> |
| --- | --- | --- | --- | --- | --- |
| Matsigenka | $q_1$ | 0.085 | 0.374 | 0.032 | |
| | $q_2$ | 0.168 | 0.153 | 0.026 | |
| | $q_3$ | 0.311 | 0.295 | 0.092 | |
| | $q_4$ | 0.045 | 0.094 | 0.004 | |
| | $q_5$ | 0.118 | 0.083 | 0.010 | 0.163 |
| U.S. | $q_1$ | 0.084 | 0.360 | 0.030 | |
| | $q_2$ | 0.164 | 0.166 | 0.027 | |
| | $q_3$ | 0.309 | 0.314 | 0.097 | |
| | $q_4$ | 0.045 | 0.082 | 0.004 | |
| | $q_5$ | 0.118 | 0.078 | 0.009 | 0.167 |

<sup>a</sup>Mean of the posterior distribution for the population mean value of $q$ for the respective growth process.
<sup>b</sup>Proportion of overall body height at age 15 contributed by the respective process (1 through 5).
<sup>c</sup>The product of the two columns immediately to the left.
<sup>d</sup>From equation A.72.

**Table A.3.** Calculations for mean overall *K* for males at 15 years since conception.

| Group | Parameter | Value<sup>a</sup> | Prop Total Height<sup>b</sup> | Prop Metab<sup>c</sup> | Product<sup>d</sup> | Sum<sup>e</sup> |
| --- | --- | --- | --- | --- | --- | --- |
| Matsigenka | $K_1$ | 81.452 | 0.374 | 0.250 | 7.617 | |
| | $K_2$ | 12.224 | 0.153 | 0.250 | 0.468 | |
| | $K_3$ | 1.527 | 0.295 | 0.286 | 0.129 | |
| | $K_4$ | 9.617 | 0.094 | 0.363 | 0.329 | |
| | $K_5$ | 7.919 | 0.083 | 0.519 | 0.342 | 8.886 |
| U.S. | $K_1$ | 81.861 | 0.360 | 0.250 | 7.359 | |
| | $K_2$ | 11.716 | 0.166 | 0.250 | 0.488 | |
| | $K_3$ | 1.466 | 0.314 | 0.294 | 0.135 | |
| | $K_4$ | 10.058 | 0.082 | 0.340 | 0.282 | |
| | $K_5$ | 8.626 | 0.078 | 0.514 | 0.345 | 8.609 |

<sup>a</sup>Mean of the posterior distribution for the population mean value of $K$ for the respective growth process.
<sup>b</sup>Proportion of overall body height at age 15 contributed by the respective process (1 through 5).
<sup>c</sup>Proportion of the original metabolic activity of the respective process at age 15 (1 through 5).
<sup>d</sup>The product of the three columns immediately to the left.
<sup>e</sup>From equation A.73

**Table A.4.**
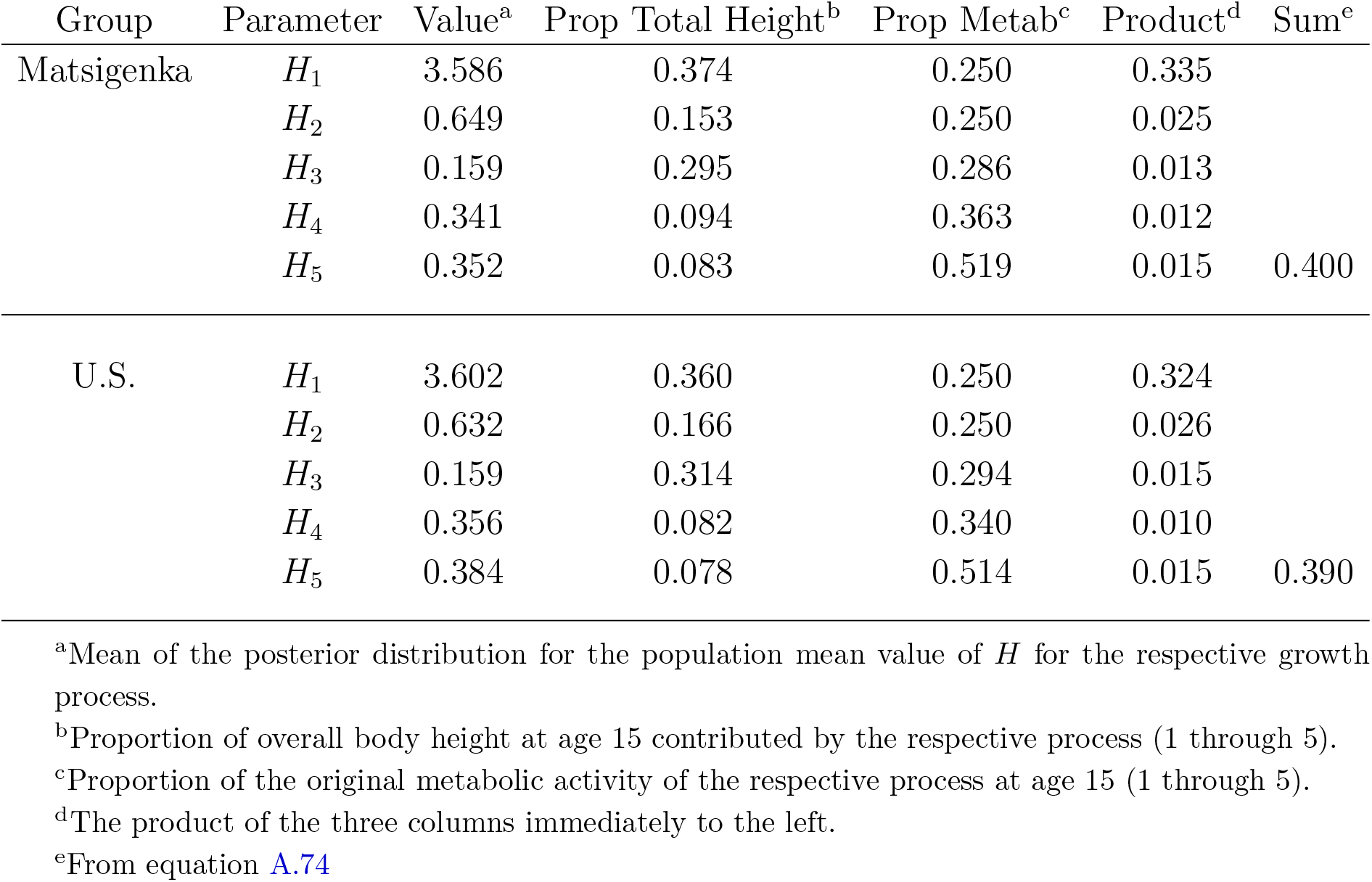
Calculations for mean overall *H* for males at 15 years since conception.

| Group | Parameter | Value<sup>a</sup> | Prop Total Height<sup>b</sup> | Prop Metab<sup>c</sup> | Product<sup>d</sup> | Sum<sup>e</sup> |
| --- | --- | --- | --- | --- | --- | --- |
| Matsigenka | $H_1$ | 3.586 | 0.374 | 0.250 | 0.335 | |
| | $H_2$ | 0.649 | 0.153 | 0.250 | 0.025 | |
| | $H_3$ | 0.159 | 0.295 | 0.286 | 0.013 | |
| | $H_4$ | 0.341 | 0.094 | 0.363 | 0.012 | |
| | $H_5$ | 0.352 | 0.083 | 0.519 | 0.015 | 0.400 |
| U.S. | $H_1$ | 3.602 | 0.360 | 0.250 | 0.324 | |
| | $H_2$ | 0.632 | 0.166 | 0.250 | 0.026 | |
| | $H_3$ | 0.159 | 0.314 | 0.294 | 0.015 | |
| | $H_4$ | 0.356 | 0.082 | 0.340 | 0.010 | |
| | $H_5$ | 0.384 | 0.078 | 0.514 | 0.015 | 0.390 |

<sup>a</sup>Mean of the posterior distribution for the population mean value of $H$ for the respective growth process.
<sup>b</sup>Proportion of overall body height at age 15 contributed by the respective process (1 through 5).
<sup>c</sup>Proportion of the original metabolic activity of the respective process at age 15 (1 through 5).
<sup>d</sup>The product of the three columns immediately to the left.
<sup>e</sup>From equation A.74

### D.5. Independent checks on parameter estimates

We can compare posterior estimates of overall allometric (*q*) and metabolic parameters (*K* and *H*) to approximations calculated using data that are independent of U.S. and Matsigenka heights and weights.

#### D.5.1. Allometric exponent (q)

Note that the values of 0.16 and 0.17 estimated in Table A.2 are considerably lower than the values of 0.43 to 0.47 estimated from head circumference of German children (Figure A.2) and waist circumference of U.S. children (Figure A.3). As explained in Appendix D.1.2, this is an expected consequence of using total body height as an approximation of height contributed only by the living cells of the skeleton (i.e., excluding bone matrix).

For anabolism and catabolism, we assume that all molecules absorbed through the intestine are broken down into smaller molecules (via catabolism), which then serve as raw materials with which to construct different, structural molecules (via anabolism). Thus, the total weight of dietary carbohydrates, protein, and fat consumed over the course of a year constitutes the weight of both destructed and constructed molecules.

#### D.5.2. Catabolic rate (K)

The catabolic rate is expressed in grams of destructed molecules per gram of body cells per year. Assume a 55 kg adolescent boy consumes approximately 152 kg of digestible carbohydrates, protein, and fat per year (https://www.ers.usda.gov/Data/FoodConsumption/, Table 3). The skeleton weighs approximately 7.5 kg (International Commission on Radiological Protection 1995, pg 14). Living skeletal cells weigh approximately 0.05*55 = 2.8 kg (Figure A.5). The weight of non-cellular water in the body is approximately 10 L * 1 kg/L = 10 kg (https://www.merckmanuals.com/home/hormonal-and-metabolic-disorders/water-balance/about-body-water).

Thus, the total weight of living cells is 55 - (7.5 - 2.8) - 10 = 40.3 kg. Fat cells (adipocytes) are metabolically active themselves (Lafontan 2012) and affect the metabolism of other cells (e.g., through leptin secretion: Kaiyala et al. (2010)). However, by weight, adipocytes are likely to have lower rates of catabolism than other types of cells, primarily because most of the volume of adipocytes comprises inert triglycerides. Assuming a 15 year old US child has a total fat mass of 17.9 kg (Borrud et al. 2010), and these fat cells have a negligible rate of catabolism compared to other body cells, then the total weight of metabolically active body cells is 40.3 - 17.9 = 22.4 kg. The rate of catabolism is then 152,000 g/year *1/22,400 g = 6.8 g/g/year. This is slightly lower than the values of 8.9 and 8.6 calculated in Table A.3 for boys at 15 years since conception. However, given the many assumptions and simplifications of the model, we are encouraged that the rate of catabolism estimated only from the shape of the growth trajectory is so close to (i.e., only 22% less than) the catabolic rate estimated independently from approximate food consumption.

#### D.5.3. Anabolic rate (H)

The anabolic rate is expressed in grams of constructed molecules per cm^2^ intestinal surface per year. Assume an adult consumes approximately 150 kg of digestible carbohydrates, protein, and fat per year (https://www.ers.usda.gov/Data/FoodConsumption/, Table 3). The surface area of the human intestine is approximately 320,000 cm^2^ (Helander and Fändriks 2014). Thus, the rate of anabolism is 150,000 g/year *1/320,000 cm^2^ = 0.47 g/cm^2^/year. This corresponds well to (i.e., is only 19% greater than) the values of 0.4 and 0.39 calculated in Table A.4 for boys at 15 years since conception.

## Appendix E. Simulating heath-care interventions

Figure A.11 shows weight trajectories, complementing the height trajectories in Figure 4, calculated for two types of healthcare intervention: 1) setting all Matsigenka metabolic parameters (*H*’s and *K*’s) at the values for U.S. children; and 2) setting Matsigenka values for *K*_2_ and *H*_3_ (girls) and *K*_2_ and *K*_3_ (boys) at the values for U.S. children. Note that the model predicts that both interventions will result in mean Matsigenka skeletal weights nearly identical to those of U.S. children, even though Matsigenka are still predicted to be shorter, on average, than U.S. children (with the exception of boys after the targeted metabolic intervention: third row of Figure 4). This suggests that, in addition to metabolism, metabolic interventions can affect the shape (i.e., allometry) of Matsigenka bodies, even though no allometric parameters (*q*’s) were modified as part of either intervention.

This can be seen by comparing the overall values of *q* by age in Figures A.12 and A.13 (representing the first and second interventions, respectively), with those in Figure 3 (pre-intervention). Note that, after both interventions, the overall mean values of *q* for Matsigenka children after birth tend to be indistinguishable or higher than the values for U.S. children, whereas they are substantially lower prior to the intervention. This is due to the fact that changing metabolic rates affect the shapes of the five component trajectories corresponding to each growth process, which in turn affect the weighting of the five *q* parameters to determine the overall *q* for a given age (Appendix D.4). In particular, note that both interventions increase the maximum asymptotic height (and thus the weighting in the overall *q* calculations) of the third (early childhood: Figure 1) growth process, for which the Matsigenka mean value of *q*_3_ is estimated to be slightly higher on average (though only marginally so) than that for U.S. children (Figure A.7). By this means, the two interventions on metabolic rates serve to increase the overall mean allometric parameter *q* for Matsigenka.

**Figure A.11.**
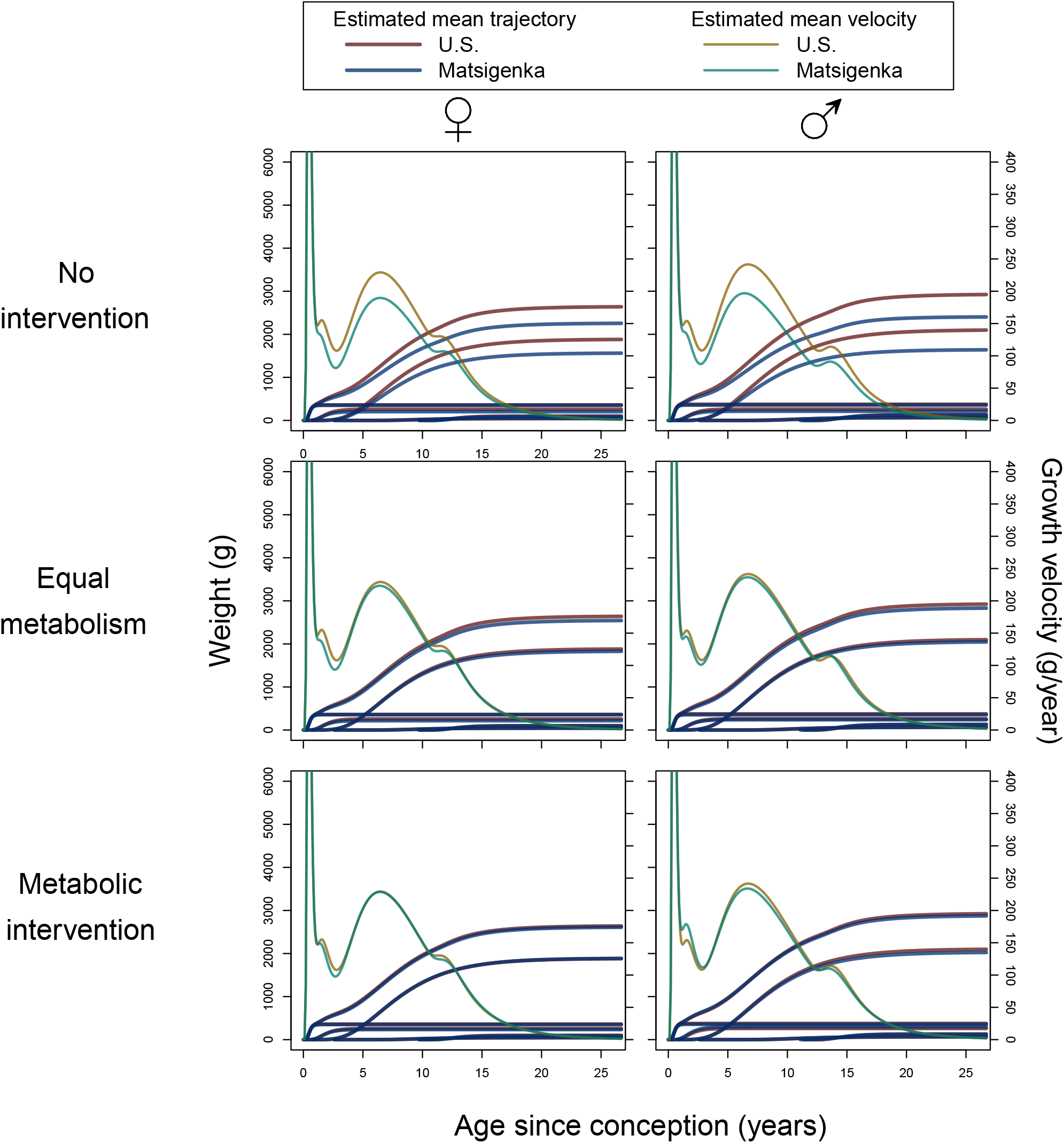
Predicted effects of healthcare interventions on Matsigenka mean weight. Shown here are mean posterior trajectories for mean overall weight and five component growth processes for U.S. (red) and Matsigenka (blue) children. Corresponding mean growth velocities are decreasing (after age 10) from left to right. The first row reproduces Figure A.9, and represents the pre-intervention state. In the second row, values of all metabolic parameters *K* and *H* for Matsigenka are modified to match mean estimates of those for U.S. children. In the third row, only Matsigenka values of *K*_2_ and *H*_3_ (girls) and *K*_2_ and *K*_3_ (boys) are modified to match mean estimates of those for U.S. children. Corresponding height trajectories are shown in Figure 4. Posterior distributions for mean characteristics of these intervention trajectories are shown in Appendix Figures A.14 and A.15.

**Figure A.12.**
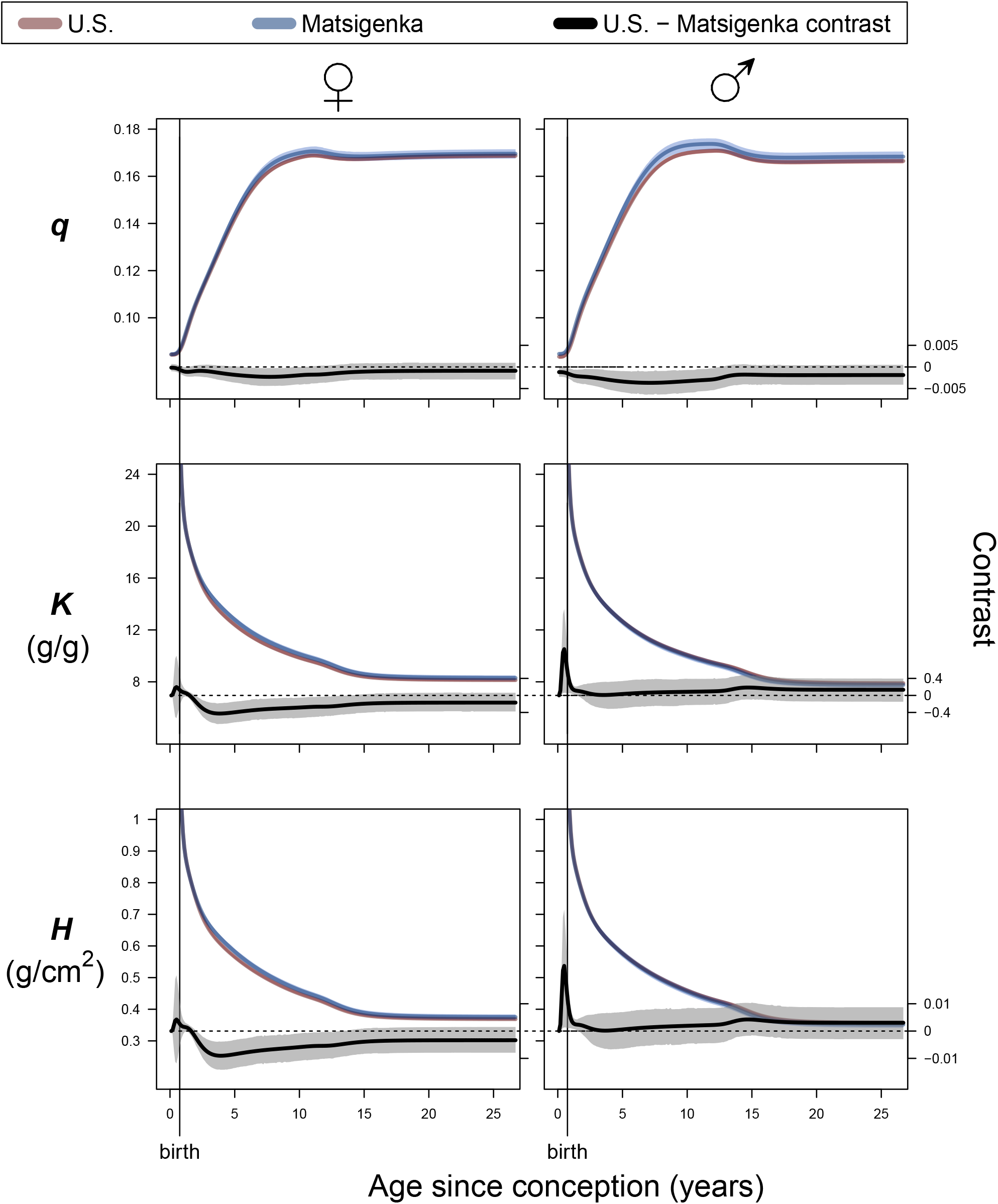
Predicted effects of the first Matsigenka healthcare intervention on mean parameter values by age. This plot is analogous to Figure 3. Estimates correspond to the first intervention (second rows of Figures 4 and A.11 in which values of all metabolic parameters (all *K*’s and *H*’s) for Matsigenka are modified to match mean estimates of those for U.S. children.

**Figure A.13.**
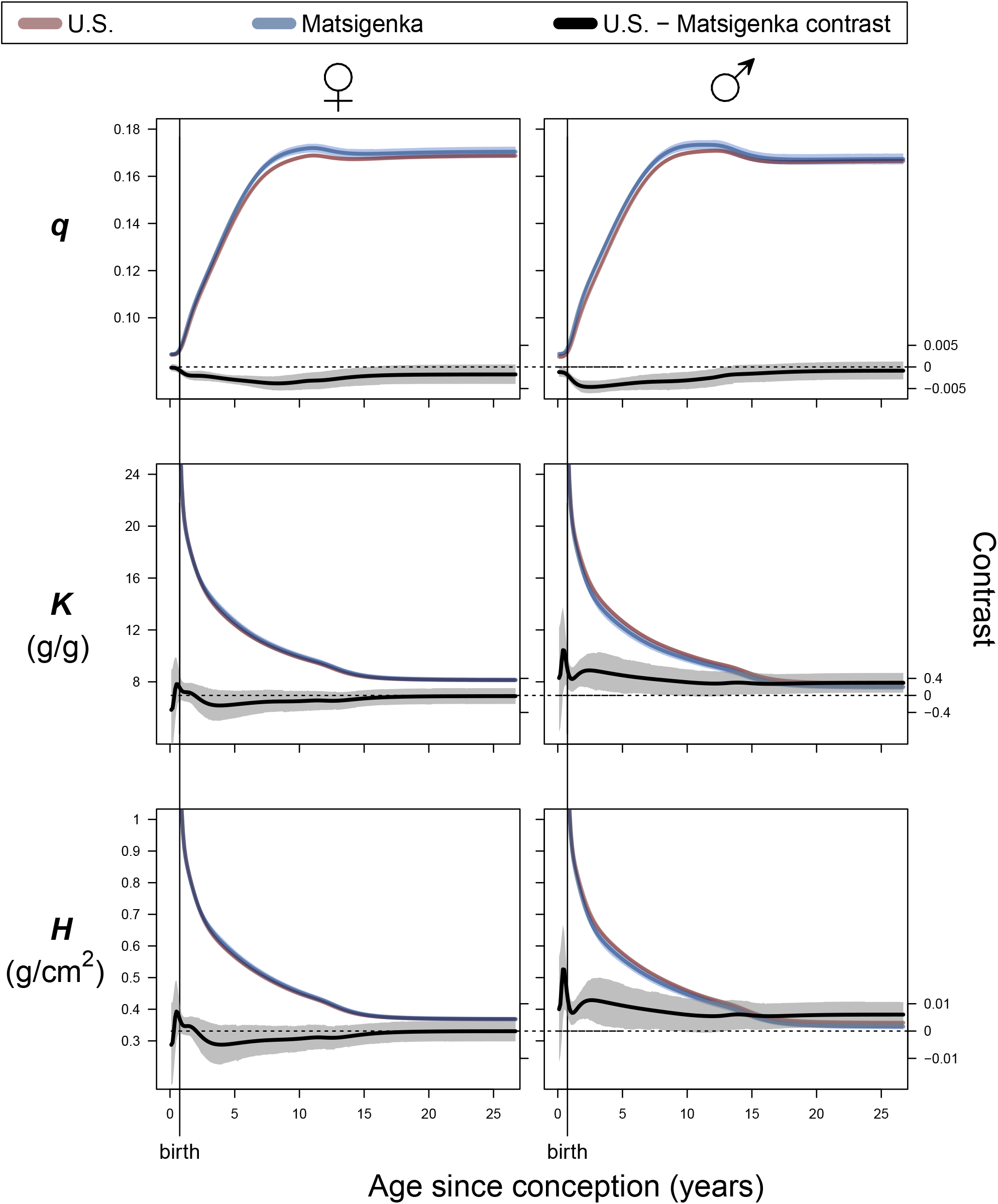
Predicted effects of the second Matsigenka healthcare intervention on mean parameter values by age. This plot is analogous to Figure 3. Estimates correspond to the second intervention (third rows of Figures 4 and A.11 in which only Matsigenka values of *K*_2_ and *H*_3_ (girls) and *K*_2_ and *K*_3_ (boys) are modified to match mean estimates of those for U.S. children.

Figures A.14 and A.15 show characteristics of the height trajectories for the first and second interventions (second and third rows of Figure 4, respectively). Comparing these against characteristics of the estimated trajectories prior to the intervention (Figure A.10), shows that both interventions are predicted to increase mean Matsigenka maximum heights, but, with the exception of boys in the second targeted intervention (Figure A.15), not enough to make them indistinguishable from mean U.S. heights. Interestingly, interventions that change only metabolic rates can also change the age at maximum growth velocity during puberty: compare values for boys in Figures A.10 and A.14. This occurs despite the fact that neither of these interventions change the ages at which the five growth processes initiate (*i* parameters). Thus, the “timing” of growth, a key parameter of the SITAR model (Cole et al. 2010), may be sensitive to metabolic rates, in addition to the age of onset of growth-promoting hormonal regimes.

**Figure A.14.**
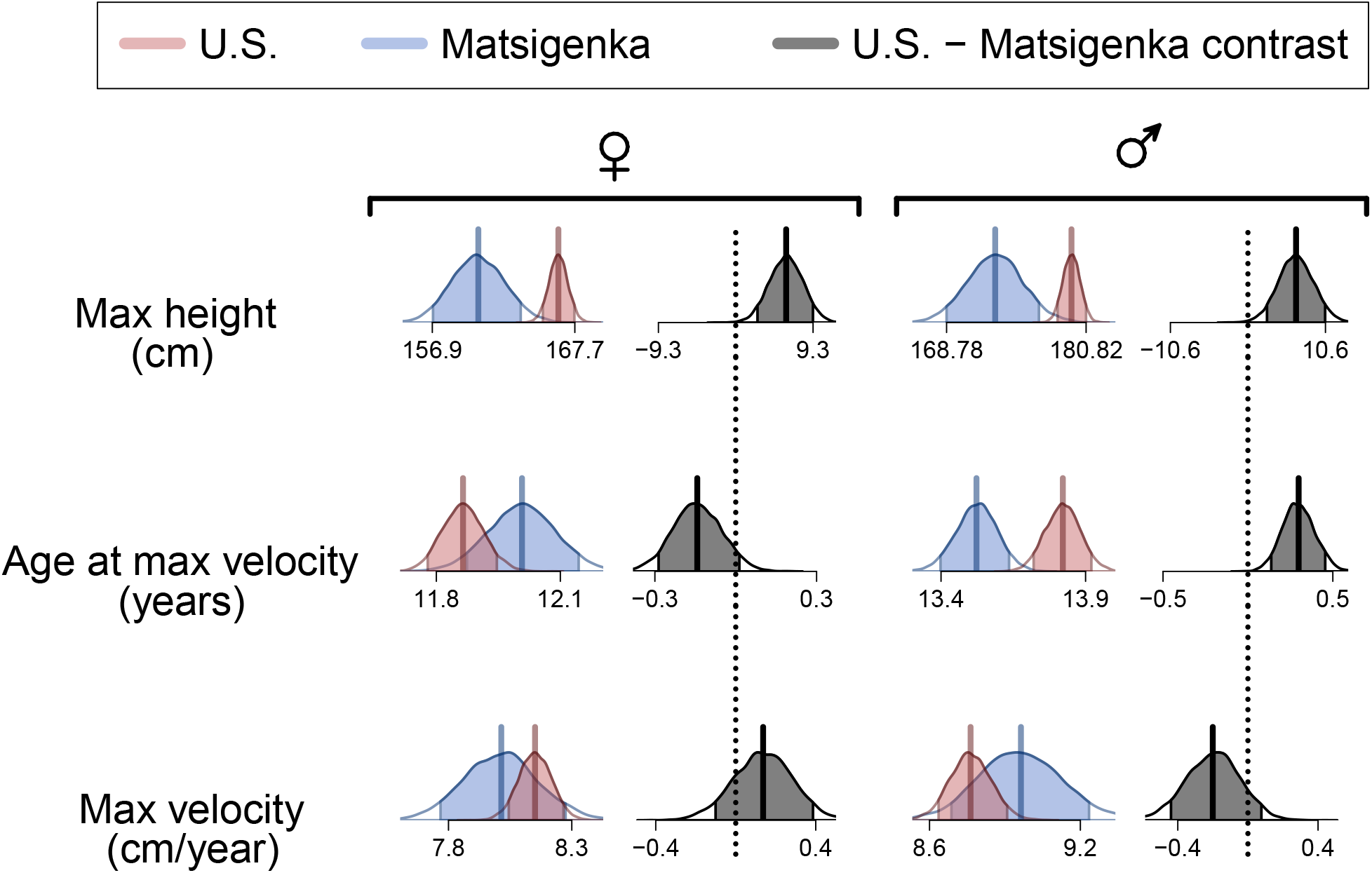
Estimated characteristics of mean height trajectories after intervention to equalize all metabolic rates. Plot features are analogous to Figure A.10. Shown are characteristics of growth trajectories in the second row of Figures 4 and A.11, corresponding to an intervention such that values of all metabolic parameters *K* and *H* for Matsigenka are modified to match mean estimates of those for U.S. children. Maximum velocity and age at maximum velocity are calculated during puberty.

**Figure A.15.**
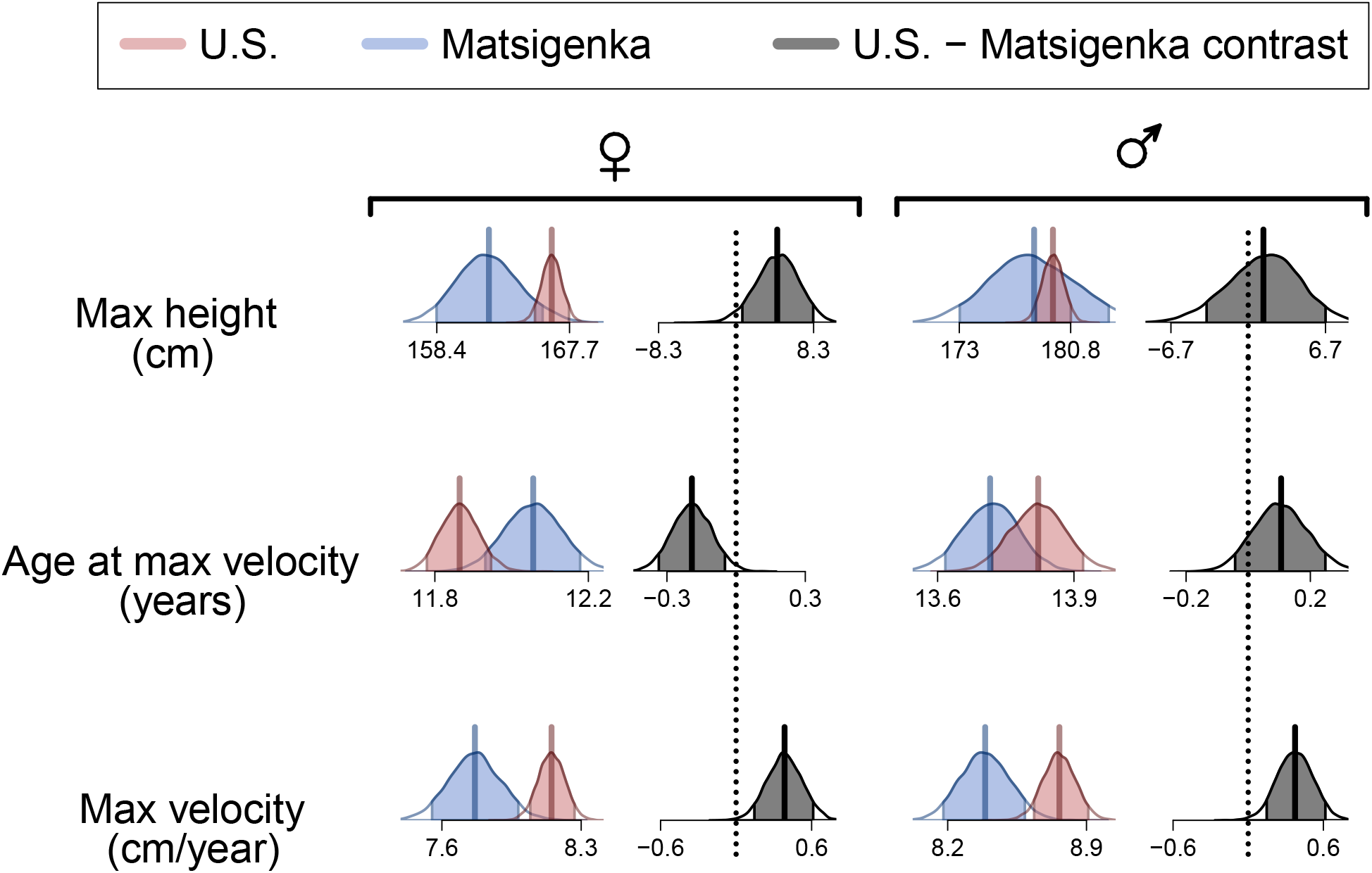
Estimated characteristics of mean height trajectories after intervention to equalize targeted metabolic rates. Plot features are analogous to Figure A.10. Shown are characteristics of growth trajectories in the third row of Figures 4 and A.11, corresponding to an intervention such that only Matsigenka values of *K*_2_ and *H*_3_ (girls) and *K*_2_ and *K*_3_ (boys) are modified to match mean estimates of those for U.S. children. Maximum velocity and age at maximum velocity are calculated during puberty.

## Appendix F. Empirical estimation of other growth models

### F.1. Unbiased height and weight model

As explained in Appendix D.1.3, we fit the main model simultaneously to height and weight data, but allowed the model to assume greater measurement error (*σ*_*µ*_) when fitting the weight data in order to account for increases in fat and muscle weight that do not necessarily correspond to increases in intestinal surface area. As an alternative, here we fit the same model but place comparable constraints on measurement error for height and weight. An upper bound of 0.005 on *σ*_*η*_ corresponds to measurement error such that approximately 95% of measurements fall within 2*stdev = 1 cm of the actual height when height = 100 cm. An upper bound of 0.01 on *σ*_*µ*_ corresponds to measurement error such that approximately 95% of measurements fall within 2*stdev = 1 kg of the actual weight when weight = 50 kg. We first derived a new baseline trajectory following the procedure in Appendix D.2. The multilevel model is identical to that in Appendix D.3, except for the following priors (priors for males shown below priors for females, where they differ):

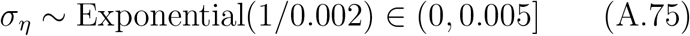

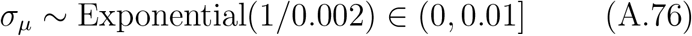

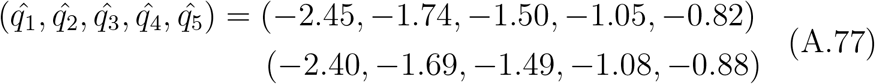

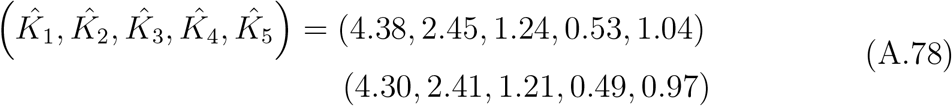

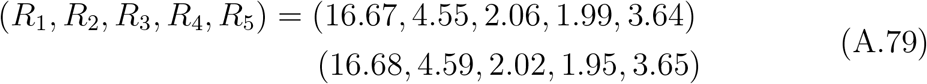

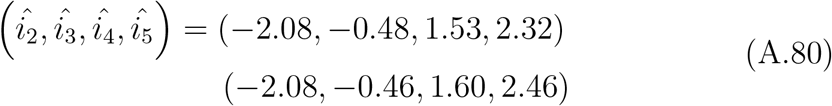

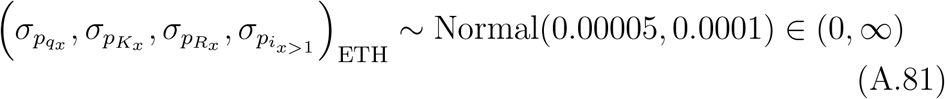

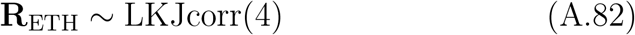

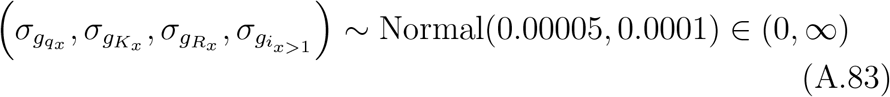

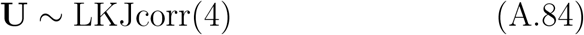

Models were fit in R (R Core Team 2022) and Stan (Stan Development Team 2022) using the cmdstanr package (Gabry and Cesnovar 2021). Convergence was indicated by 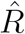 values of 1.00. This required four chains of 4000 to 5000 samples each, half of which were warm-up. Data and analysis scripts are available at https://github.com/jabunce/bunce-fernandez-revilla-2022-growth-model.

Figure A.16 shows growth trajectories calculated from mean posterior parameter estimates. Posterior mean standard deviations for height (*σ*_*η*_) for females and males are both approximately 0.005 (i.e., the upper bound in equation A.75). Posterior mean standard deviations for weight (*σ*_*µ*_) for females and males are both approximately 0.01 (i.e., the upper bound in equation A.76). Note that, as a consequence, here weight trajectories fit the weight data better than they do in Figure 2 for the model allowing greater measurement error when fitting weight. However, this comes at the cost of fitting the height data seemingly less well, with very abrupt transitions in height velocity that do not appear realistic. We expect height to be more strongly correlated with intestinal surface area than is weight, as height is not affected by increases in fat and muscle, which affect weight but not necessarily intestinal surface area (Appendix D.2). Thus, we prefer to draw inferences from the model in the main text, which biases fitting in favor of the height data, rather than from this unbiased model.

**Figure A.16.**
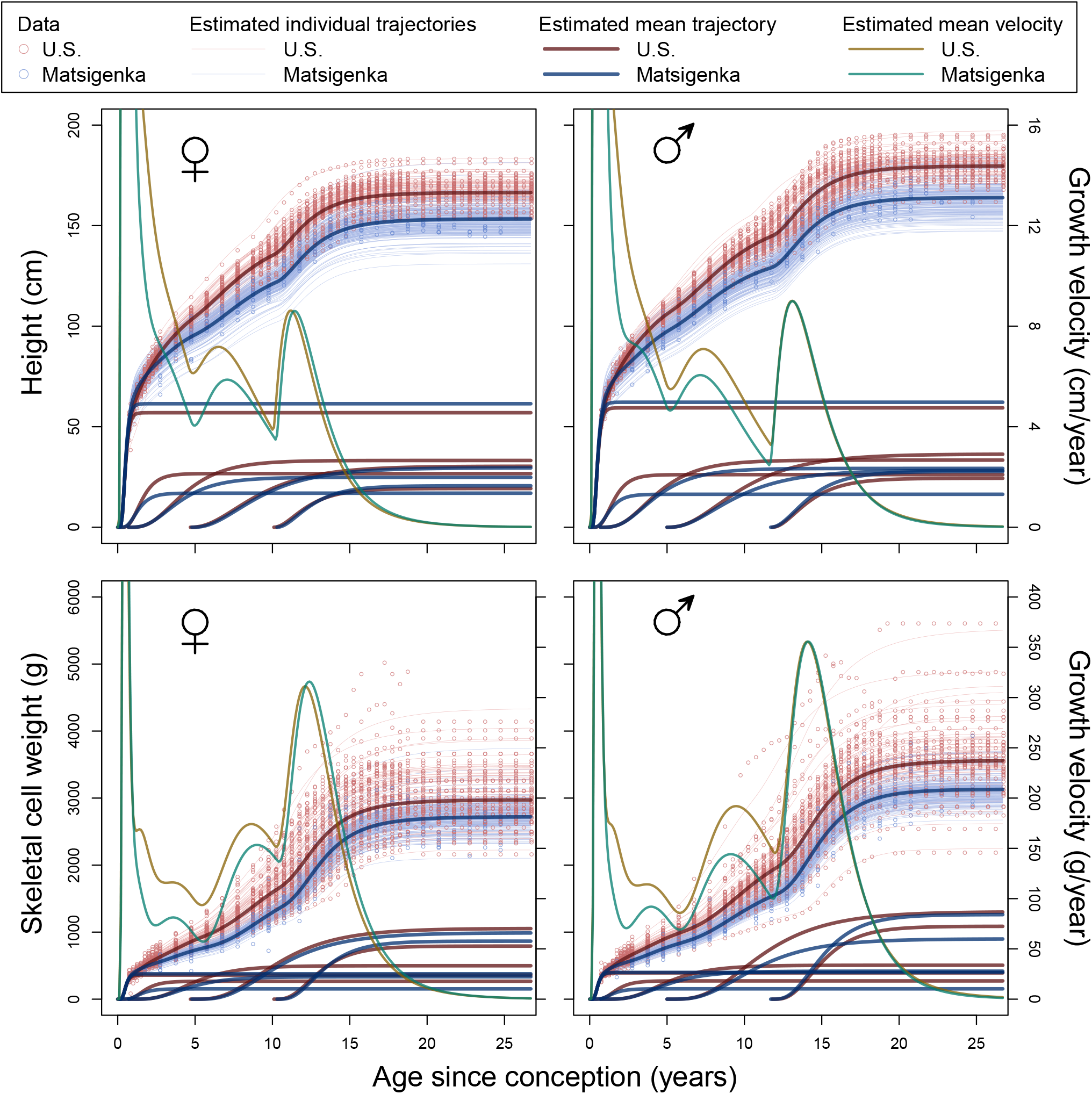
Estimated height trajectories from model fit in unbiased way to height and weight data. This is analogous to Figures 2 and A.9.

### F.2. Height–only model

For comparison we fit the model just to height data. We first derived a new baseline trajectory following the procedure in Appendix D.2. The multilevel model is identical to that in Appendix D.3, except for the absence of equations A.43, A.45, and A.55, and that the following priors are different (priors for males shown below priors for females, where they differ):

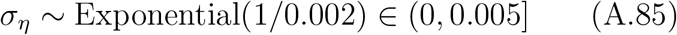

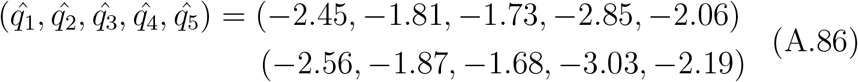

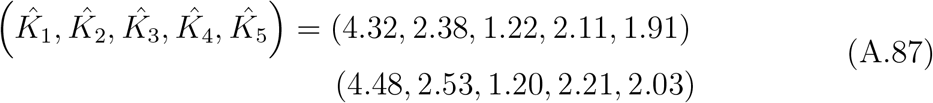

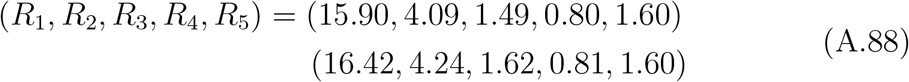

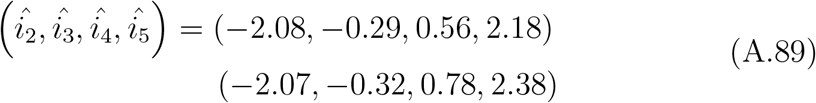

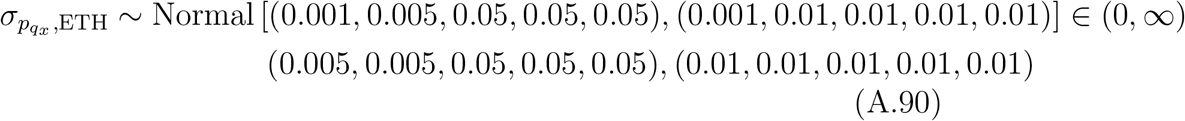

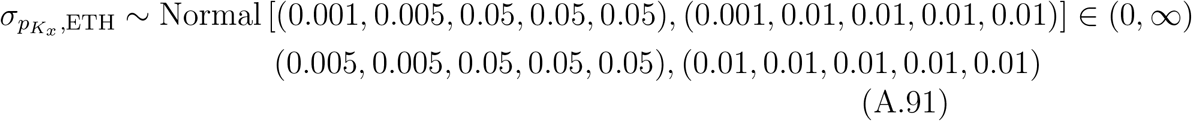

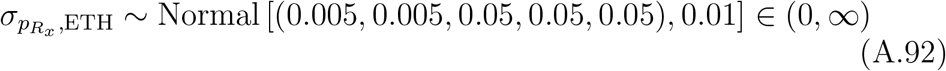

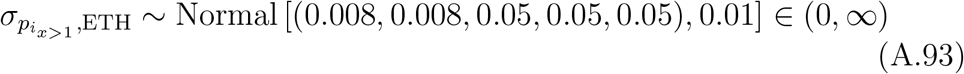

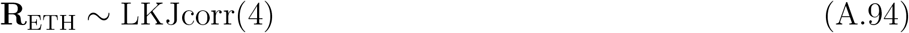

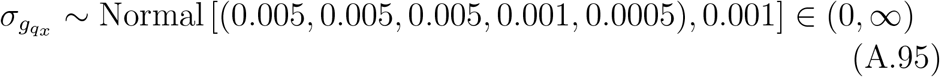

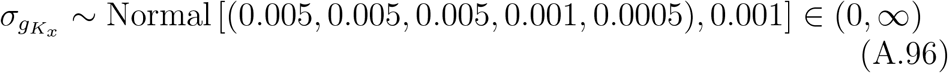

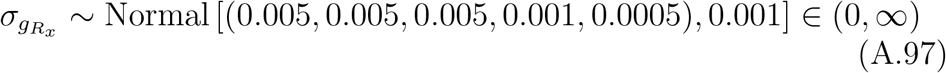

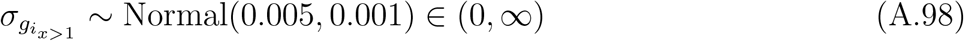

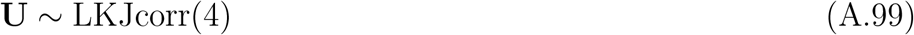

Models were fit in R (R Core Team 2022) and Stan (Stan Development Team 2022) using the cmdstanr package (Gabry and Cesnovar 2021). Convergence was indicated by 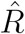 values of 1.00. This required four chains of 6000 samples each, half of which were warm-up. Data and analysis scripts are available at https://github.com/jabunce/bunce-fernandez-revilla-2022-growth-model.

Figure A.17 shows both height and weight trajectories calculated from mean posterior parameter estimates. Posterior mean standard deviations (*σ*_*η*_) for females and males are both approximately 0.005 (i.e., the upper bound in equation A.85). Note that weight trajectories fit the weight data very poorly. Thus, when model parameters are estimated using height data alone, their values tend to be different from the values estimated from a model fit simultaneously to height and weight. This is true even though the height trajectories change very little between the model fit to height alone, and the model fit to both height and weight (compare Figures 2 and A.17). As explained in Appendix D.2, this signals the importance of fitting the model simultaneously to both height and weight in order estimate parameter values that better reflect both aspects of body growth.

**Figure A.17.**
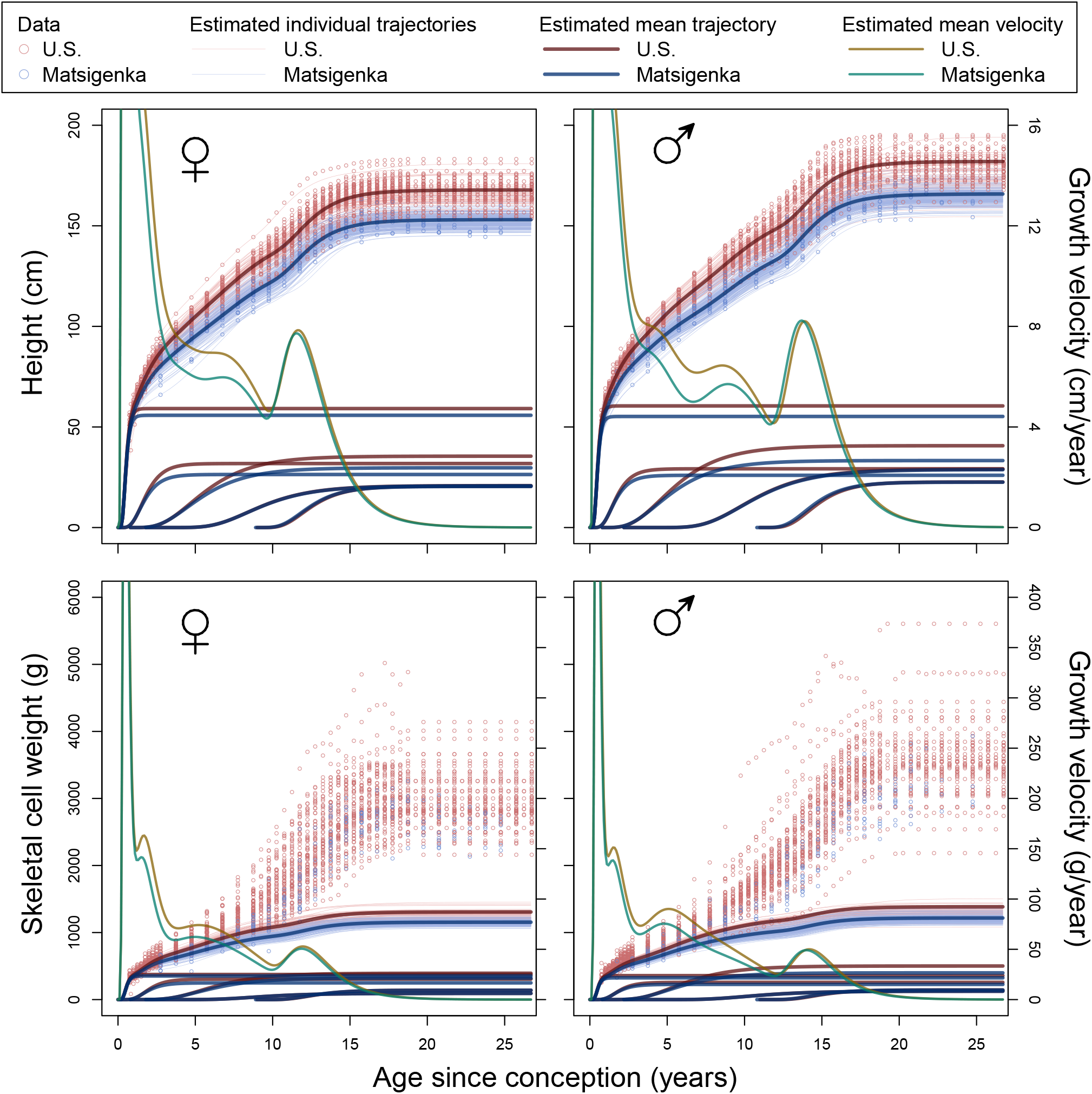
Estimated height trajectories from model fit just to height data. This is analogous to Figures 2 and A.9. Mean posterior parameter estimates are used to construct both height and weight trajectories.

### F.3. JPA-1 model

For comparison, we fit the seven-parameter JPA-1 model of Jolicoeur et al. (1992) to height data from U.S. and Matsigenka boys. Following the strategy in Appendix D.2, we first find a baseline trajectory by fitting the JPA-1 to heights of U.S. boys, treated as cross-sectional data. Prior means on the parameters are taken from Table 4 of Jolicoeur et al. (1992), which reports parameter estimates after fitting the model to data from 13 French boys. In the absence of prior knowledge, standard deviations are chosen to facilitate model convergence. Observed height at time *t, h*_*t*_, is modeled as:

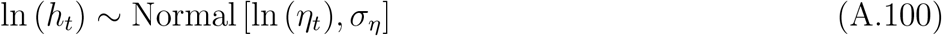

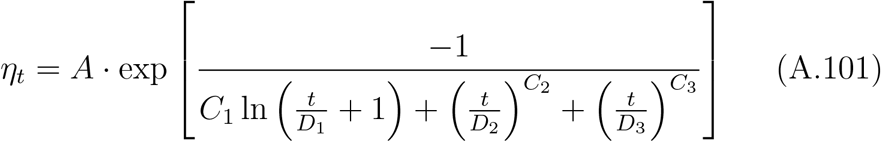

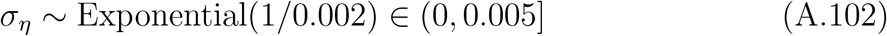

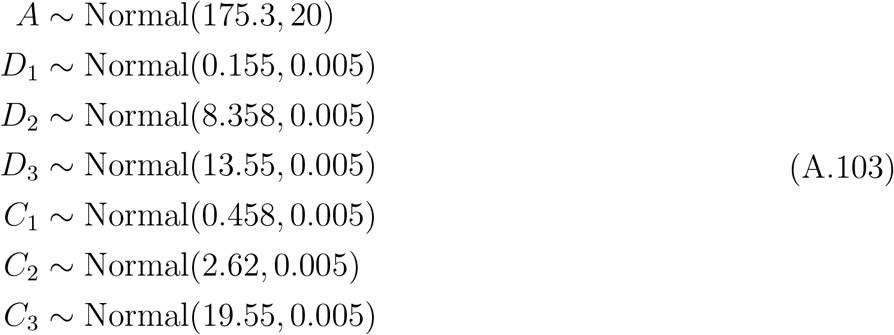

Posterior parameter means estimated from the baseline model are used as prior means for parameters in the multi-level model, which is analogous to that in Appendix D.3. In the absence of prior knowledge, priors on the standard deviations of the group- and person-level offsets are chosen to facilitate model convergence. Person *j*’s observed height (*h*_*jt*_) at time *t* is modeled as:

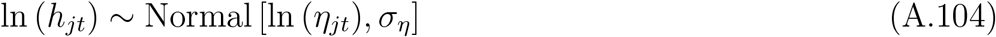

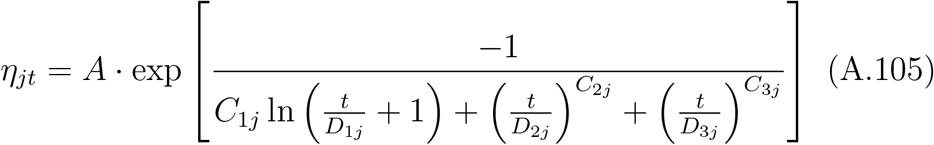

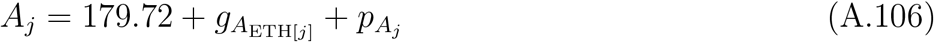

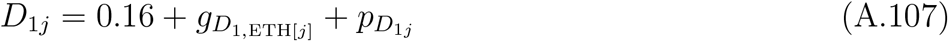

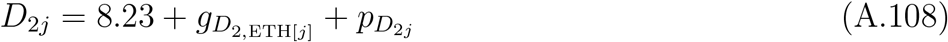

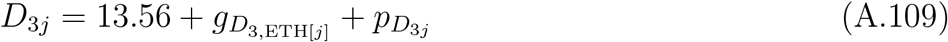

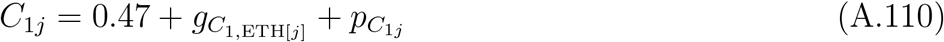

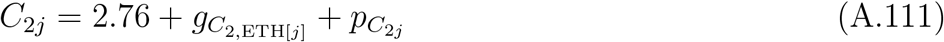

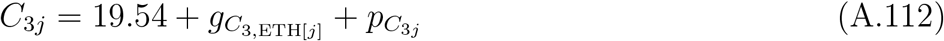

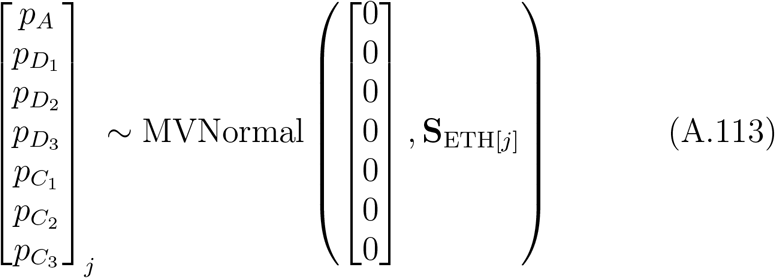

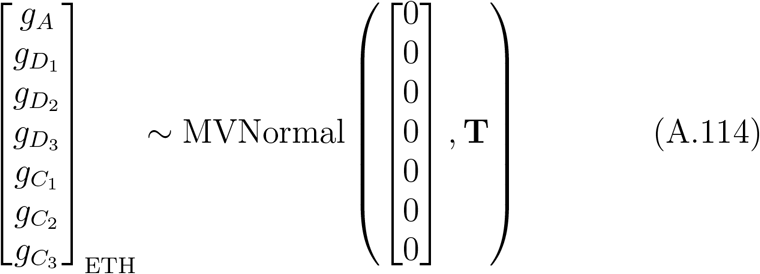

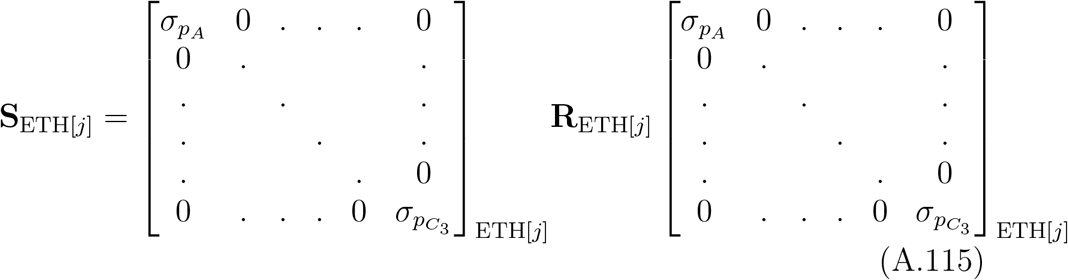

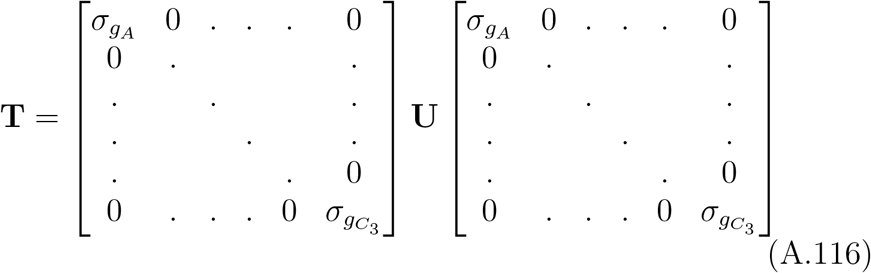

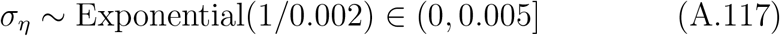

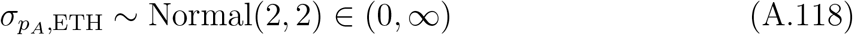

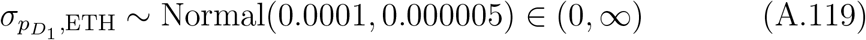

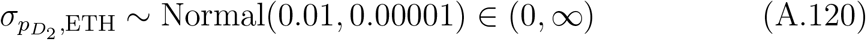

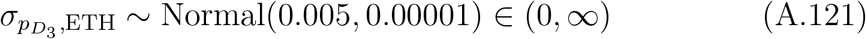

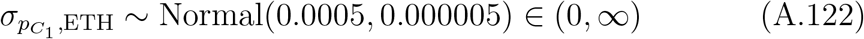

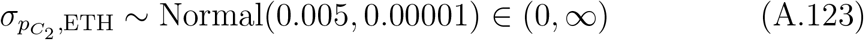

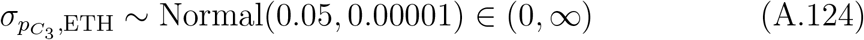

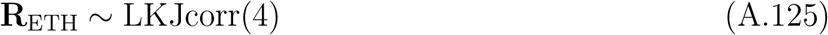

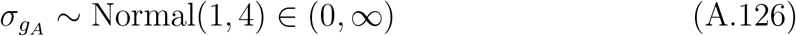

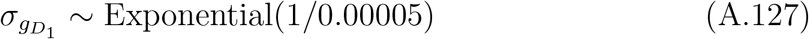

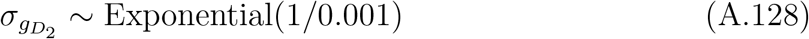

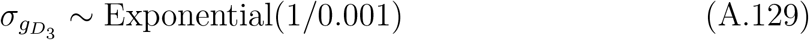

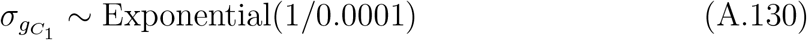

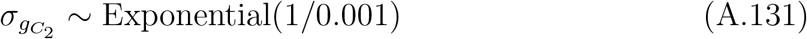

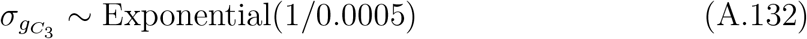

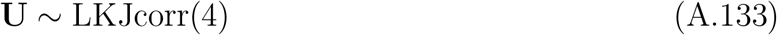

The baseline and multilevel models were fit in R (R Core Team 2022) and Stan (Stan Development Team 2022) using the cmdstanr package (Gabry and Cesnovar 2021). Convergence of the multilevel model was indicated by 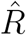 values ≤ 1.01. This required eight chains of between 6000 and 9000 samples each, half of which were warm-up. Note that we have been unable to find priors yielding 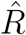 values of 1.00 for all parameters, as is the case for the other models presented here. Data and analysis scripts are available at https://github.com/jabunce/bunce-fernandez-revilla-2022-growth-model. Figure A.18 shows the posterior mean height trajectories for U.S. and Matsigenka boys.

**Figure A.18.**
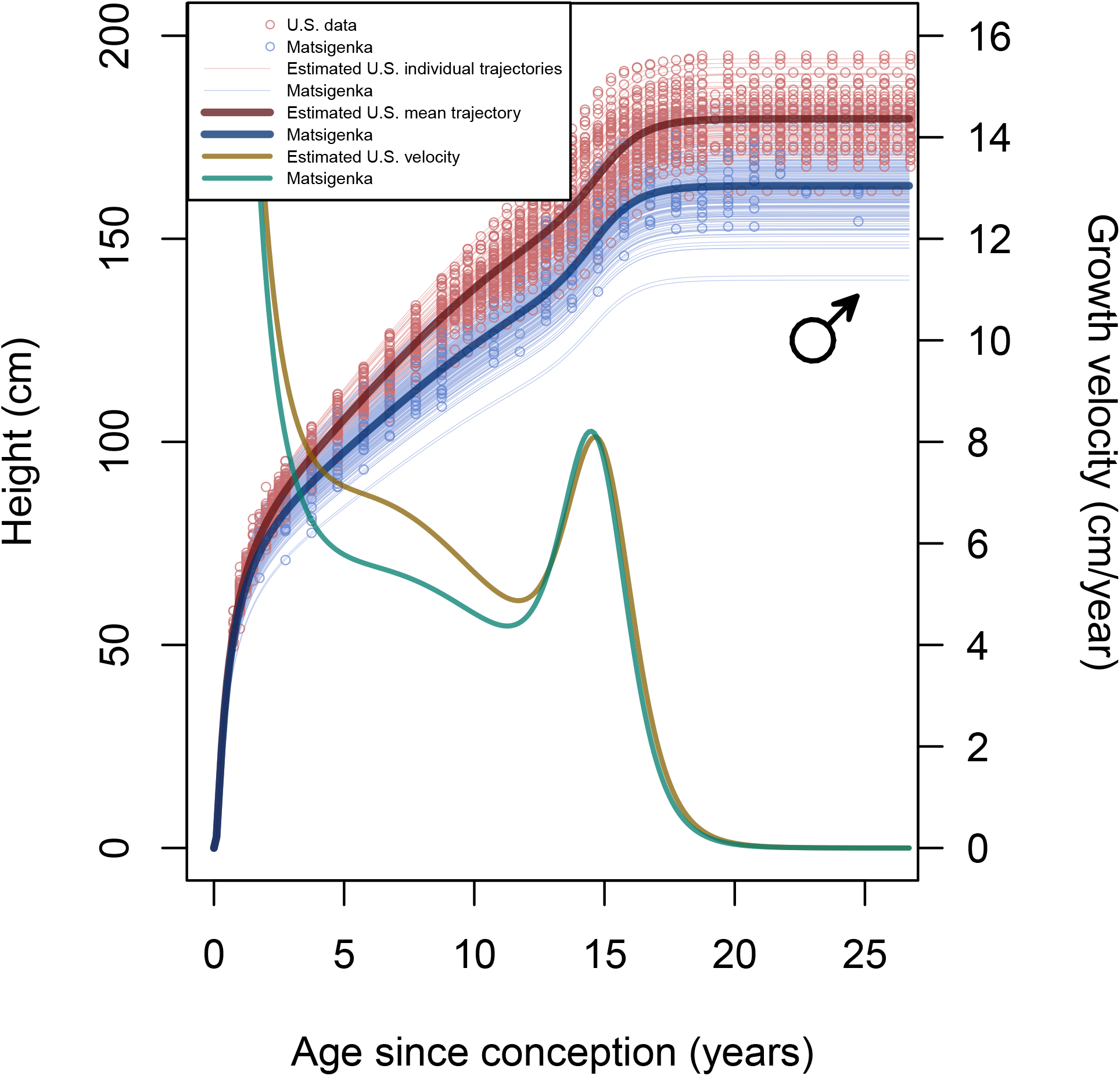
Estimated mean height trajectories using the JPA-1 model of Jolicoeur et al. (1992). Shown are the mean posteriors for the population mean trajectories from the multilevel JPA-1 model in equations A.104–A.133, fit to height measurements from U.S. (red) and Matsigenka (blue) boys. The mean posterior velocity trajectories are shown decreasing (after age 14) from left to right. Note that velocity at conception (age = 0) is undefined in the JPA-1 model.

### F.4. SITAR model

For comparison, we fit the SITAR model of Cole et al. (2010) to height data from U.S. and Matsigenka boys using the SITAR package (Cole 2021) in R. Data and analysis scripts are available at https://github.com/jabunce/bunce-fernandez-revilla-2022-growth-model.

The model uses a spline with ten degrees of freedom to fit the overall mean height trajectory across all individuals, as well as a fixed effect for ethnic group and three individual-level random effects (termed size, timing, and intensity) that relate group-mean and individual-level trajectories to the estimated overall mean trajectory. Using SITAR to simultaneously model two datasets that differ in temporal resolution, as we do here, was pioneered by Cole et al. (2016).

Note that the dataset used to fit the SITAR model differs from the dataset used to fit all other models in this paper, in that here we remove the constraint that each individual’s height is (approximately) zero at conception. We were unable to find a SITAR model that converged without errors when this constraint was in place. Consequently, this model estimates height to be negative until shortly after conception. Figure A.19 presents the estimated group-mean height trajectories for U.S. and Matsigenka boys.

**Figure A.19.**
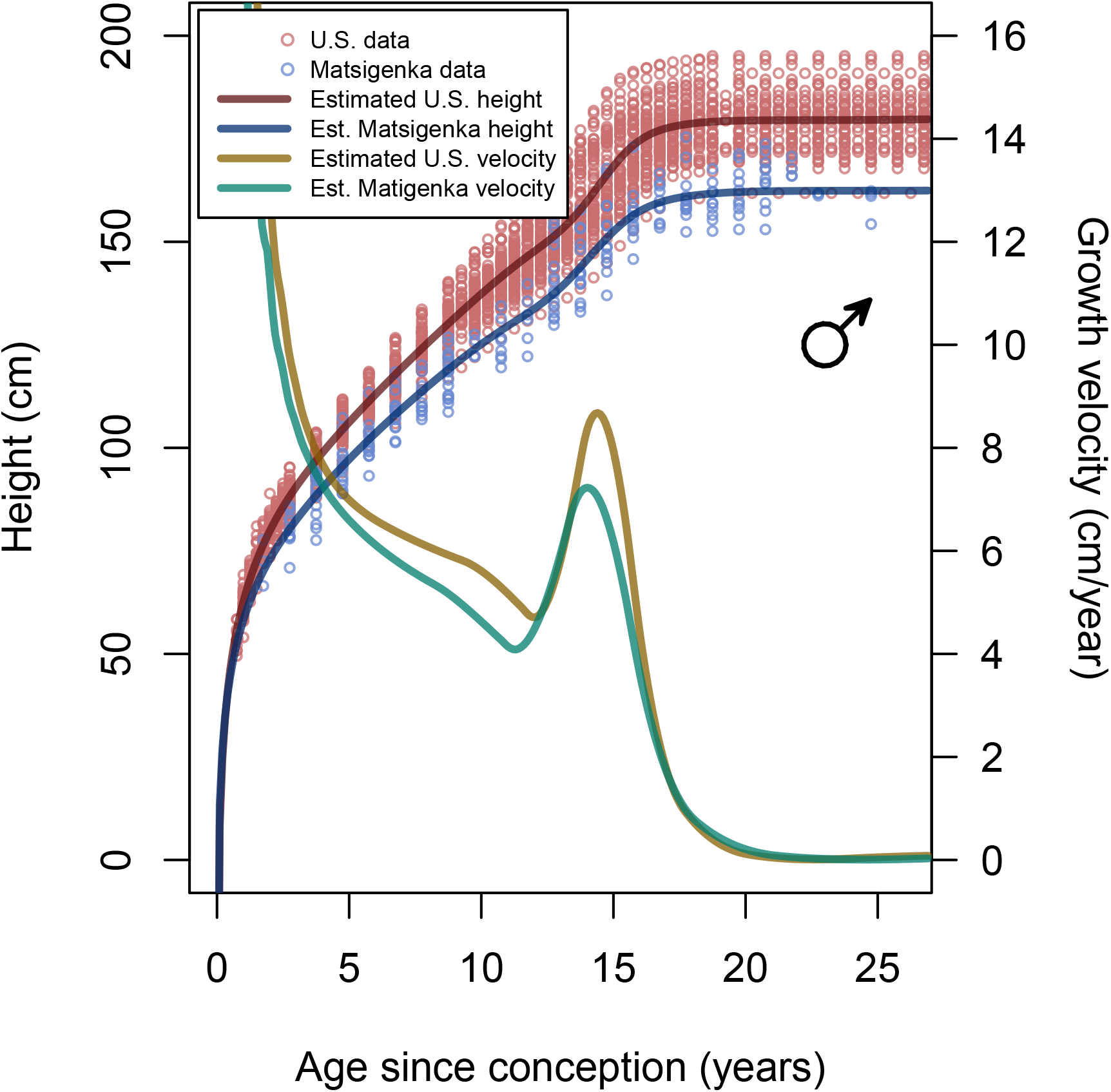
Estimated mean height trajectories using the SITAR model of Cole et al. (2010). Shown are the estimated group-mean trajectories from the SITAR model as implemented in the SITAR R package (Cole 2021), fit to height measurements from U.S. (red) and Matsigenka (blue) boys. The estimated group-mean velocity trajectories are shown decreasing (after age 14) from left to right. Note that we have been unable to find a SITAR model that convergences when forced to pass through (or very close to) the origin (i.e., height approximately 0 cm at conception). At age = 0.001 years since conception, this model estimates height to be negative.

## Appendix G. Analysis of model fit

### G.1. Comparison with other models

Comparing mean height trajectories as estimated by the composite model in the main text, the JPA-1 model (Appendix F.3), and the SITAR model (Appendix F.4), shortcomings in all three models are apparent upon visual inspection of the estimated mean trajectories and the residuals when fit to data from U.S. boys. Importantly, however, we note that the data from the U.S. population does not comprise actual measurements of children. Rather these data have been manually smoothed by taking a moving weighted average of the actual measurements (Tuddenham and Snyder 1954, pg 193–198). Thus, it must be kept in mind that there is more variability in the measurements than is reflected in the U.S. data to which these models are fit and compared, and any corresponding analysis of residuals must be interpreted with caution.

Figure A.20 suggests that the composite model tends to underestimate mean U.S. male height at both the beginning and end of the pubertal growth spurt. Figures A.20 and A.19, show that the only SITAR model we have found that converges satisfactorily does not pass through the origin (height approximately 0 cm at conception), and consequently estimates height to be negative until shortly after conception. Figure A.21 suggests that the SITAR model may overestimate mean height (length) at birth (observed − predicted height *<* 0).

Comparing the mean U.S. male height velocities estimated by these three models (Figure A.20), all three models show a diminution of deceleration (Tanner and Cameron 1980) after age five. The JPA-1 and SITAR models show very similar velocities. Compared to these, the composite model estimates a lower growth velocity at ages 4 and 12, a higher velocity after age 18, and an earlier age for the peak velocity during the pubertal growth spurt.

The composite model suggests that Matsigenka boys, on average, attain a peak velocity at an earlier age during the pubertal growth spurt, than do U.S. boys (Figures 2 and A.10). This is consistent with both the JPA-1 model (Figure A.18) and the SITAR model (Figure A.19). As can also been seen in these plots, the composite model and the SITAR model estimate Matsigenka mean peak pubertal height velocity to be lower than that of U.S. boys, while the JPA-1 model estimates this velocity to be slightly higher than that of U.S. boys.

**Figure A.20.**
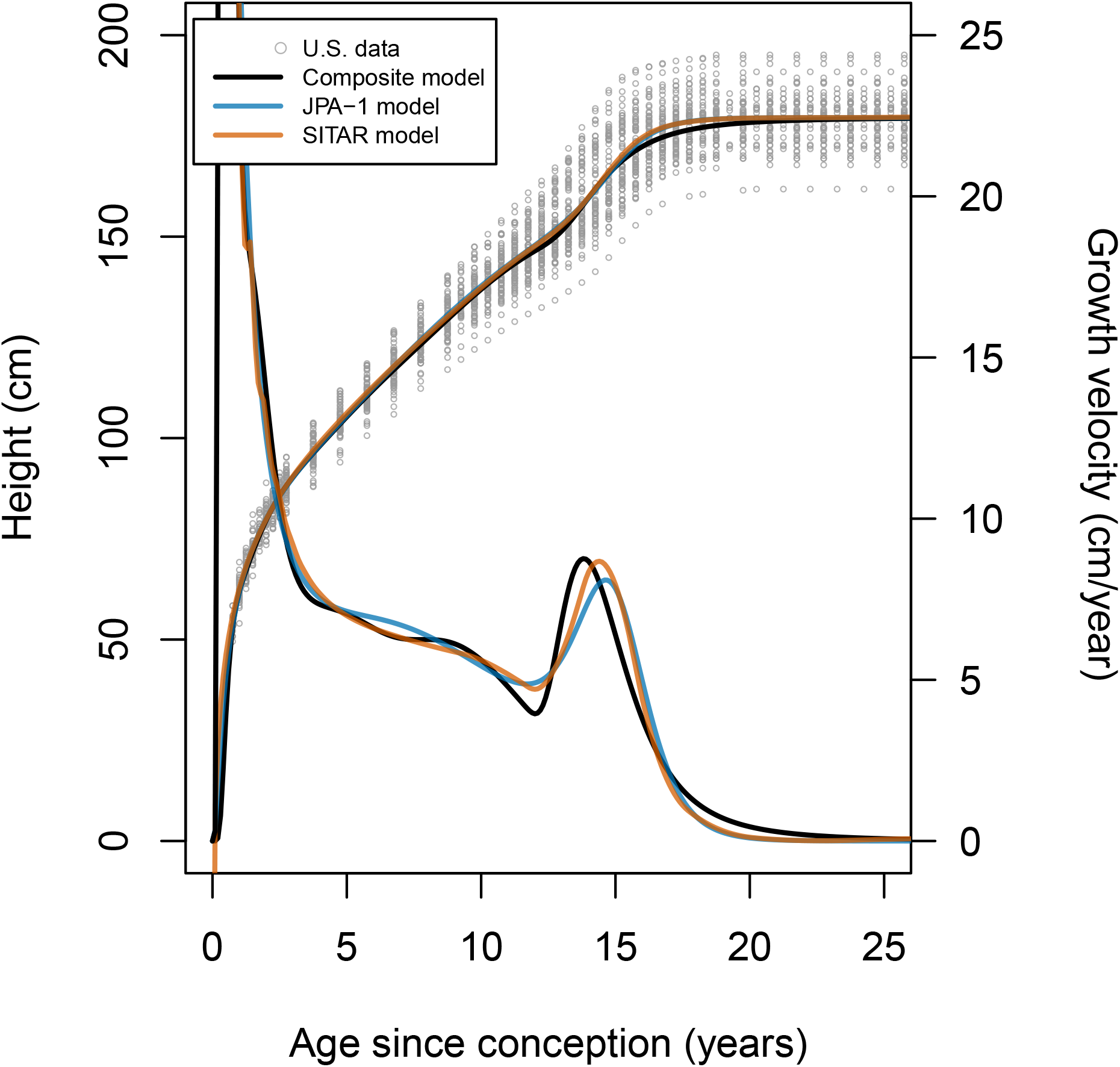
Comparison of growth models. Three alternative models estimate a mean growth trajectory using height measurements from U.S. boys. The composite model is that presented in the main text and Appendix D.3. The parametric JPA-1 model of Jolicoeur et al. (1992), and the spline-based shape-invariant SITAR model of Cole et al. (2010) are described in Appendices F.3 and F.4, respectively. Estimated height trajectories are ascending, and the corresponding velocity curves are all descending after age 14, from left to right.

**Figure A.21.**
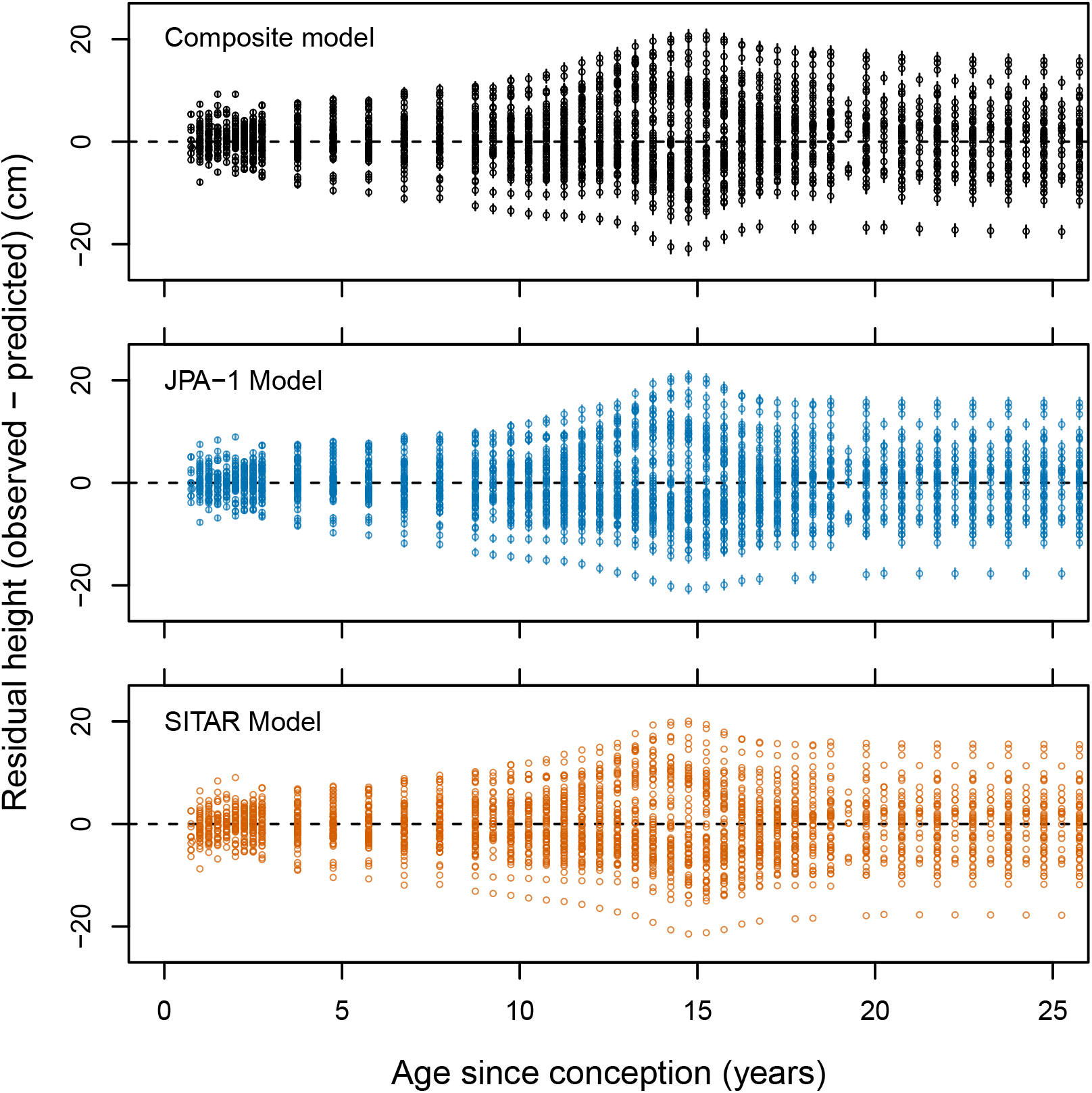
Cross-model comparison of residuals from the estimated group mean height trajectory. Residuals are the measured heights of U.S. boys minus the predicted height of the mean U.S. boy at the corresponding age, taken from the posterior mean trajectory of the composite model in the main text (upper), the JPA-1 model described in Appendix F.3 (middle), and the SITAR model described in Appendix F.4 (lower). Bars for the Composite and JPA-1 models represent the 90% HPDI of each residual estimate, and points represent the means. There are no error bars for the SITAR model, as it was not fit in a Bayesian framework. If a model represents the group mean trajectory accurately, the residuals at each age should be distributed approximately symmetrically about zero.

### G.2. Analysis of residuals

The principle aim of the empirical analysis in the main text is to compare the mean height and weight trajectories of U.S. and Matsigenka children. Thus, it is of interest to know whether the inaccuracies in the estimated mean growth trajectories of the composite growth model with respect to observed heights and weights manifest similarly for the two ethnic groups. If the patterns of inaccurate prediction are different for U.S. and Matsigenka children, then it would be inappropriate to compare their population-mean growth trajectories using these models. Figure A.22 plots the posterior means of the residuals of the estimated mean U.S. and Matsigenka height and weight trajectories. Importantly, the patterns of asymmetry around zero for residuals at different ages of U.S. children are echoed for Matsigenka children, though with fewer data points. Note the increase in inaccuracy of height predictions around the pubertal growth spurt, which shows much inter-individual variability in onset, length, and intensity. Also note the generally greater inaccuracy in weight predictions compared to height predictions, as expected and designed (Appendix D.1.3).

Figure A.23 plots the posterior means of the residuals of the estimated individual U.S. and Matsigenka height and weight trajectories (i.e., incorporating both group-level and individual-level offsets from the multi-level model). If a model represents individual growth trajectories accurately, the residuals at each age should be as close to zero as possible and distributed approximately symmetrically around zero. As is apparent, there is much room for improvement of the composite model. Importantly, the distributions of such individual-level residuals constitute one metric by which future improvements in within-sample model prediction (fit) can be judged (complementing metrics of out-of-sample model prediction, such as WAIC: McElreath (2020)). To improve the accuracy of future model estimates, we favor strategies that aim to refine the theory (described in the main text) upon which these growth models are based.

Note, in particular, that inaccuracy in individual height predictions increase markedly during the pubertal growth spurt for both U.S. and Matsigenka children. This suggests that the model is not yet flexible enough to track individual-level differences in the onset, duration, and intensity of the growth spurt. Additionally, the model tends to overestimate height of individual U.S. children after puberty. Potential solutions include widening the priors on individual-level offsets to model parameters. At present, we haven’t been able to find wider priors that still permit model convergence. However, this is an area for future work. As above, the generally greater inaccuracy in weight predictions compared to height predictions is by design, and, on theoretical grounds, is not necessarily expected to improve (Appendix D.1.3).

**Figure A.22.**
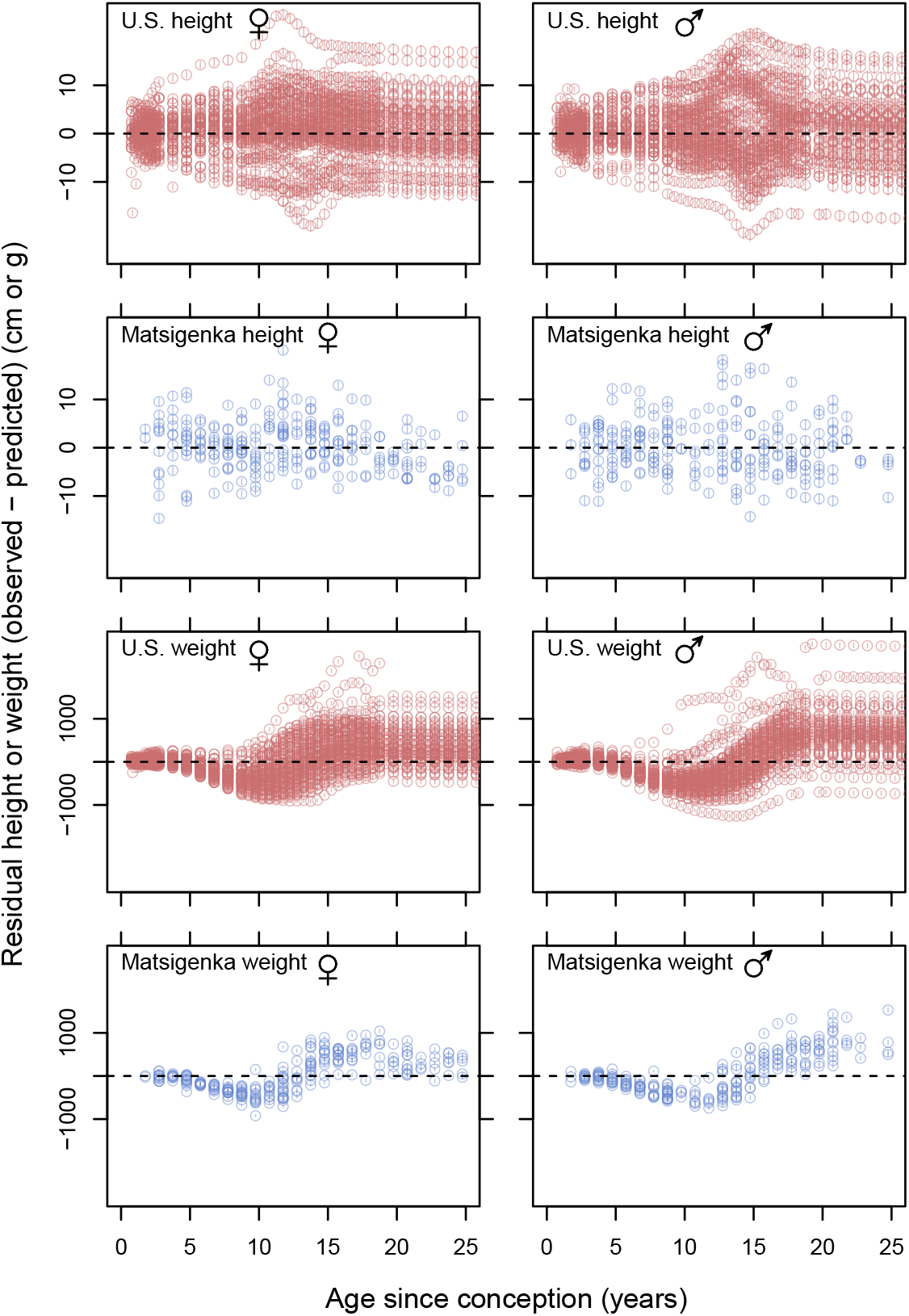
Residuals from estimated group mean height and weight trajectories. Residuals are the measured heights and weights of U.S. and Matsigenka children minus the mean posterior predicted height or weight of the mean U.S. or Matsigenka child at the corresponding age, taken from the posterior mean trajectory of the multi-level composite model in the main text. If a model represents the group mean trajectory accurately, the residuals at each age should be distributed approximately symmetrically about zero. Note that the pattern of asymmetries about zero for the U.S. estimates is echoed in the Matsigenka estimates, though with fewer data points. This suggests that relative comparisons of estimated U.S. and Matsigenka growth trajectories are possible.

**Figure A.23.**
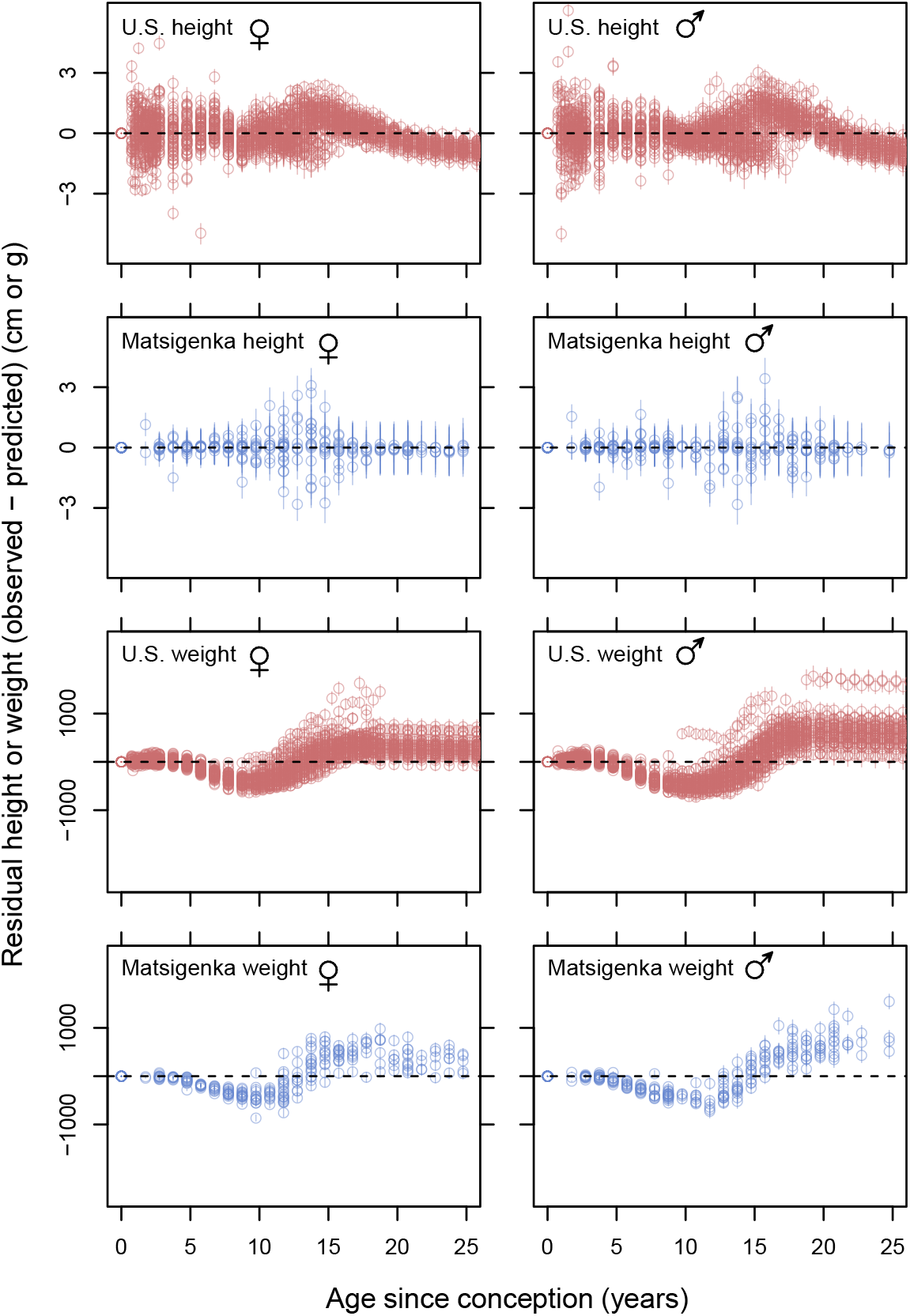
Residuals from the estimated individual mean height and weight trajectories. This figure is analogous to Figure A.22, except that residuals now represent differences between measured heights and the mean posterior predicted height or weight of the corresponding individual U.S. or Matsigenka child at the corresponding age (i.e., incorporating both group-level and individual-level offsets from the multi-level model). If a model represents individual growth trajectories accurately, the residuals at each age should be as close to zero as possible and distributed approximately symmetrically about zero. This forms the basis for judging future improvements in model fit, resulting from refinement of the theory upon which the growth model is based.

### G.3. Comparison with weight-for-height

Figure A.24 plots weight for height for U.S. and Matsigenka children. Note that, starting at a height of approximately 130 cm (around ages 10–13: Figure 2), Matsigenka girls and boys tend to weigh more than U.S. children for a given height. This contrasts with higher overall estimates of the *q* parameter for U.S. children compared to Matsigenka children (Figure 3). Recall that the composite growth model is intentionally permitted to fit weight data poorly compared to height data (Appendix D.1.3). The discrepancy between high Matsigenka weight-for-height and low estimated mean *q* is likely due to the fact that, starting around ages 10–13, mean Matsigenka weights may be underestimated more than are mean U.S. weights (Figure A.9). Underestimation of weight for a given height leads to lower estimates of radius *r*, and, therefore, given *r* = *h*^*q*^ (equation 4), lower estimates for the allometric parameter *q*. Given the definition of *K* (destructed mass per unit of an organism’s mass), underestimation of weight would also lead to an overestimate of *K*.

**Figure A.24.**
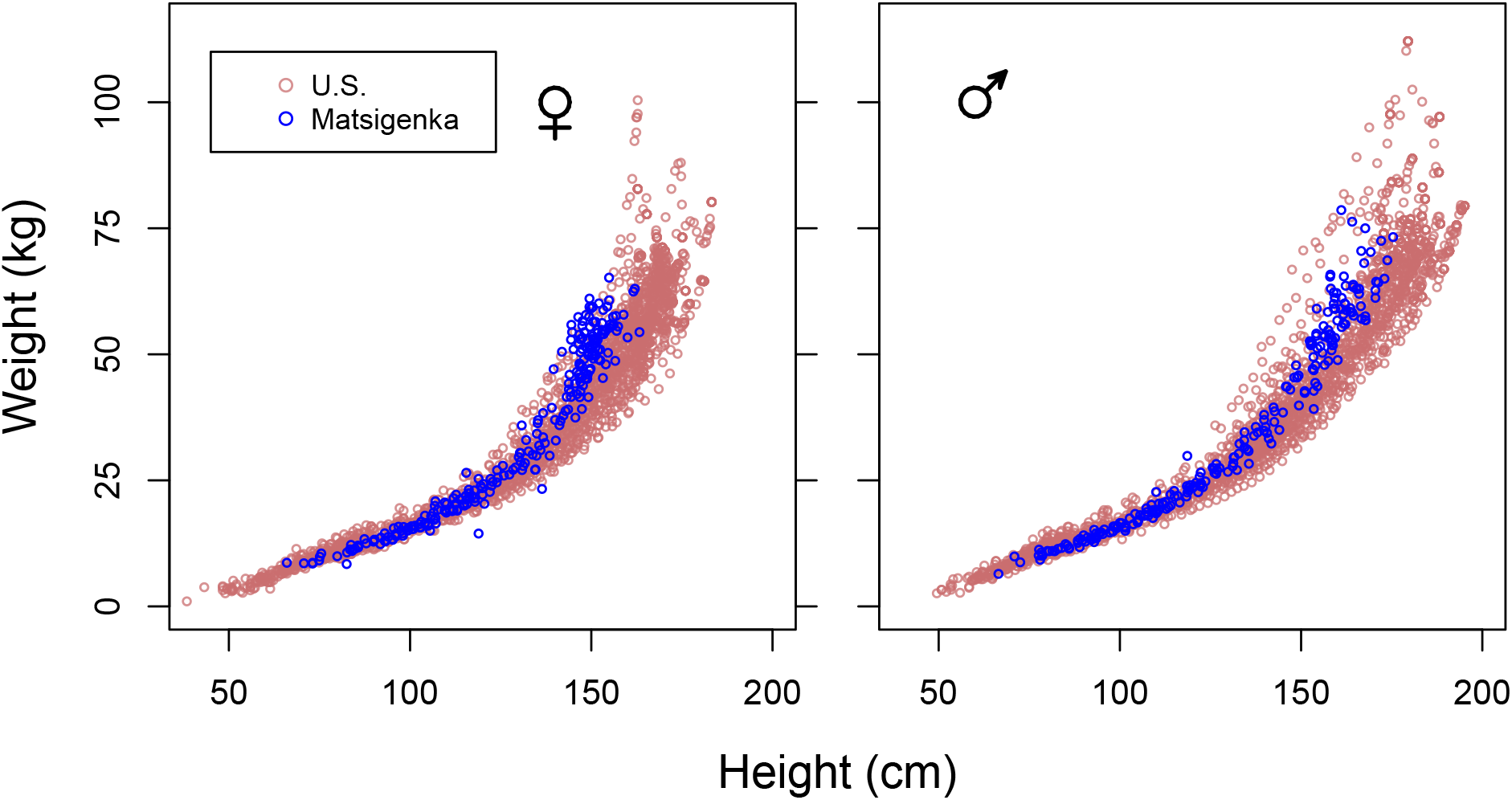
Weight for height for U.S. and Matsigenka children. Note that, starting at a height of approximately 130 cm, Matsigenka girls and boys tend to weigh more than U.S. children for a given height.

